# Spatial scenario of tropical deforestation and carbon emissions for the 21^st^ century

**DOI:** 10.1101/2022.03.22.485306

**Authors:** Ghislain Vieilledent, Christelle Vancutsem, Clément Bourgoin, Pierre Ploton, Philippe Verley, Frédéric Achard

## Abstract

Tropical forests are disappearing at an alarming rate due to human activities. Here, we provide spatial models of deforestation in 92 countries covering all the tropical moist forests in the world. Our results question the global effectiveness of protected areas in curbing deforestation and allow reinterpreting the impact of roads on deforestation in terms of both accessibility and forest fragmentation. Using our models, we derived high-resolution pantropical maps of the deforestation risk and future forest cover for the 21^st^ century under a “business-as-usual” scenario based on the deforestation rates observed in the 2010s. Under this scenario, 48% (39–56%) of tropical moist forests are expected to disappear during the course of the 21^st^ century, and 41 tropical countries will have lost all their forests by 2100. The remaining forests in 2100 will be highly fragmented and located in remote places, preferentially in protected areas, far from roads and villages, and at high elevations. We also show that future deforestation will likely concern forests with higher aboveground carbon stocks, and hence that carbon emissions from tropical deforestation are expected to increase (from 0.432–0.585 Pg/yr in 2020 to 0.583–0.628 Pg/yr in 2100). Considering also the decrease in carbon uptake in aboveground biomass (from 0.589 Pg/yr in 2000 to 0.312 Pg/yr in 2100) associated with the decrease in forest cover, tropical moist forests would become a major net carbon source in the 21^st^ century under this scenario.

## 1 Introduction

Tropical forests are at the heart of concerns when it comes to climate change and biodiversity loss, which both pose unprecedented threats to human civilization (Cardinale et al. 2012, IPCC 2014). Commonly called the “Jewels of the Earth”, tropical forests shelter 30 million species of plants and animals, representing half of the Earth’s wildlife and at least two-thirds of its plant species (Gibson et al. 2011, Wilson et al. 2012). Through carbon sequestration, they play an important role in the global carbon cycle and regulate the global climate (Baccini et al. 2017). Intact tropical forest ecosystems also prevent outbreaks of zoonoses and reduce the risk of pandemics (Tollefson 2020). At a local scale, tropical forests regulate the regional climate (Dickinson and Kennedy 1992), cooling the atmosphere (Ellison et al. 2017), and facilitate access to water (Ellison et al. 2017, Zhang and Wei 2021). They also protect against erosion and flooding (Bradshaw et al. 2007). Close to 1.6 billion people (a quarter of the world’s population) rely on forest resources for their livelihoods (FAO 2020). Despite the many ecosystem services they provide, tropical forests are disappearing at an alarming rate (Hansen et al. 2013, Achard et al. 2014, FAO 2020, Vancutsem et al. 2021), mostly because of human activities (Geist and Lambin 2002, Curtis et al. 2018). Past studies have estimated that about 10 Mha of tropical forest (including moist and dry forests) are disappearing yearly (FAO 2020). At this rate, will there still be any tropical forests left at the end of the 21^st^ century? If so, where will they be concentrated, and what will be the consequences of tropical deforestation on carbon emissions and climate in the future?

Forecasting forest cover change is paramount as it allows us to foresee the consequences of deforestation (in terms of carbon emissions, biodiversity loss, or water supply) under various technological, political, and socio-economic scenarios, and to inform decision makers accordingly (Clark et al. 2001). The “business-as-usual” scenario is of particular interest as it makes it possible to predict the likely future in the absence of change in deforestation rates, and if necessary, to alert decision-makers to an essential change of course to avoid any potential environmental disaster. While models and scenarios of carbon dioxide emission and climate change have been developed for several years by the Intergovernmental Panel on Climate Change (IPCC 2014) and are now widely used by the scientific community and known to the general public, equivalent models and scenarios for land-use change and biodiversity at the global scale are still relatively scarce (Cramer et al. 2004, Titeux et al. 2016, Pereira et al. 2020). Moreover, baseline scenarios of deforestation and associated carbon dioxide emission are necessary for implementing REDD+ (Reducing Emissions from Deforestation and forest Degradation) activities in the framework of the Paris Agreement on climate change (Goetz et al. 2015). Spatialized forest cover change scenarios are crucial because both forest carbon stocks (Avitabile et al. 2016, Baccini et al. 2017) and biodiversity (Kremen et al. 2008, Mittermeier et al. 2011) vary considerably in space at fine scale. Non-spatial scenarios of forest cover change (FAO 2020) cannot be used to forecast associated carbon emissions and change in biodiversity accurately, or for systematic conservation planning at the local scale. Spatial forecasts of forest cover change are based on spatial statistical models, which enable the estimation of a probability of change in space as a function of a set of spatial predictors (Rosa et al. 2014). In addition to forecasts, statistical models can be used to identify the main drivers of deforestation and quantify their relative effects. For example, models can be used to assess the impact of roads on the risk of deforestation (Laurance et al. 2014) and the effectiveness of protected areas in reducing deforestation (Andam et al. 2008, Wolf et al. 2021).

Few authors have attempted to provide spatialized forest cover change scenarios in the tropics at large spatial scales. The most significant studies to date have focused on modelling and forecasting forest cover change at the scale of the Amazonian basin (Soares-Filho et al. 2006, Swann et al. 2015, Aguiar et al. 2016). In this paper, we used high-resolution spatial data to model and forecast deforestation at the pantropical scale. This was made possible by the recent availability of pantropical spatial datasets of forest cover change (Vancutsem et al. 2021) and of global spatial datasets of explanatory factors related to deforestation at the required resolution (World Database on Protected Areas, SRTM Digital Elevation Database, and OpenStreetMap). We combine these extensive datasets in a spatial statistical model to test the effectiveness of protected areas in reducing deforestation and assess the impact of roads on the risk of deforestation at the pantropical scale. Assuming a business-as-usual scenario, we derive high-resolution maps of deforestation risk and future forest cover over the 21^st^ century in the humid tropics. We also estimate the carbon emissions associated with projected deforestation to assess whether tropical forests will act as a carbon source or sink in the future and conduct an uncertainty analysis.

## 2 Methods

We present below a summary of the materials and methods used in this study. A more detailed description of the workflow using the Republic Democratic of the Congo as an example can be find in the *SI Appendix*, Materials and Methods.

### 2.1 Study-areas and data

We modelled the spatial deforestation process for 119 study-areas representing 92 countries in the three tropical continents (America, Africa, and Asia), see *SI Appendix*, Fig. S1. Study-areas cover all the tropical moist forest in the world, at the exception of some islands (eg. Sao Tome and Principe or Wallis-and-Futuna). For each study-area, we derived past forest cover change maps on two periods of time: January 1^st^ 2000– January 1^st^ 2010, and January 1^st^ 2010–January 1^st^ 2020, from the annual forest cover change product by Vancutsem et al. (2021) at 30 m resolution (*SI Appendix*, Fig. S2 and Table S1). For the forest definition, we only considered *natural old-growth tropical moist forests*, disregarding plantations and regrowths. We included degraded forests (not yet deforested) in the forest definition. To explain the observed deforestation in the period 2010–2020, we considered a set of spatial explanatory variables (*SI Appendix*, Fig. S3-S6) describing: topography (altitude and slope, 90 m resolution), accessibility (distances to the nearest road, town, and river, 150 m resolution), forest landscape (distance to forest edge, 30 m resolution), deforestation history (distance to past deforestation, 30 m resolution), and land conservation status (presence of a protected area, 30 m resolution). This set of variables was selected based on a priori knowledge of the spatial deforestation process in the tropics (*SI Appendix*, Materials and Methods). Data for explanatory variables were extracted from extensive global data-sets (World Database on Protected Areas, SRTM Digital Elevation Database, and OpenStreetMap) and had a resolution close to the original resolution of the forest cover change map (30 m, see *SI Appendix*, Table S2).

### 2.2 Sampling

For each study-area, we built a large dataset from a sample of forest cover change observations in 2010–2020. We performed a stratified balanced sampling between deforested and non-deforested pixels in the period 2010–2020. Pixels in each category were sampled randomly (*SI Appendix*, Fig. S7). The number of sampled observations in each study-areas was a function of the forest area in 2010. Datasets included between 2,398 (for Sint Maarten island in America) and 100,000 (for study-areas with high forest cover such as the Amazonas state in Brazil, Peru, DRC, and Indonesia) observations. The global data-set included a total of 3,186,698 observations: 1,601,810 of non-deforested pixels and 1,584,888 of deforested pixels, corresponding to areas of 144,163 ha and 142,647 ha, respectively (*SI Appendix*, Table S3).

### 2.3 Statistical model

Using sampled observations of forest cover change in the period 2010–2020, we modelled the spatial probability of deforestation as a function of the explanatory variables using a logistic regression (*SI Appendix*, Eq. S1). To account for the residual spatial variation in the deforestation process, we included additional spatial random effects for the cells of a 10 × 10 km spatial grid covering each study-area (*SI Appendix*, Fig. S8). Spatial random effects account for unmeasured or unmeasurable variables that explain a part of the residual spatial variation in the deforestation process which is not explained by the fixed spatial explanatory variables already included in the model (such as local population density, local environmental law enforcement, etc.). Spatial random effects were assumed spatially autocorrelated through an intrinsic conditional autoregressive (iCAR) model (*SI Appendix*, Eq. S1). Variable selection for each study area was performed using a backward elimination procedure and parameter inference was done in a hierarchical Bayesian framework (*SI Appendix*, Tables S4–S9).

### 2.4 Model performance

Using a cross-validation procedure, we compared the performance of the iCAR model at predicting the spatial probability of deforestation with three other statistical models: a null model, a simple generalized linear model (equivalent to a simple logistic regression without spatial random effects), and a Random Forest model. These two last models have been commonly used for deforestation modelling (*SI Appendix*, Materials and Methods).

### 2.5 Deforestation risk and future forest cover

Using rasters of explanatory variables at their original resolution, and the fitted iCAR model for each study-area including estimated spatial random effects (*SI Appendix*, Fig. S9), we computed the spatial probability of deforestation at 30 m resolution for the year 2020 for each study-area (*SI Appendix*, Fig. S10). For each study-area, we also estimated the mean annual deforested area (in ha/yr) for the period 2010–2020 from the past forest cover change map (*SI Appendix*, Tables S14–S15). Using the mean annual deforested area in combination with the spatial probability of deforestation map, we forecasted the forest cover change on the period 2020–2110 with a time step of 10 years, assuming a “business-as-usual” scenario of deforestation (*SI Appendix*, Fig. S11 and Tables S16–S17). The business-as-usual scenario makes the assumption of an absence of change in both the deforestation intensity and the spatial deforestation probability in the future.

### 2.6 Carbon emissions associated with deforestation

We estimated the carbon emissions associated with past deforestation (2010–2020) and projected deforestation (2030–2110) using three different global or pantropical above-ground dry biomass maps at either 1 km (WUR map, Avitabile et al. 2016), 100 m (ESA CCI map, Santoro et al. 2021), or 30 m (WHRC map, Zarin et al. 2016) resolution (*SI Appendix*, Figs. S12–S13, and Table S18). We used the IPCC default carbon fraction of 0.47 (McGroddy et al. 2004) to convert biomass to carbon stocks. We assumed no change of the forest carbon stocks in the future. We estimated average annual carbon emissions for ten-year periods from 2010 to 2110. Under a “business-as-usual” scenario of deforestation, the change in mean annual carbon emissions in the future is only attributable to the spatial variation of forest carbon stocks and to the location of future deforestation. We also used annual rates of aboveground net biomass change for old-growth tropical rainforests (+1.0, +1.3 and +0.7 Mg/ha/yr for America, Africa, and Asia, respectively, Requena Suarez et al. 2019) to estimate the change in the ability of moist tropical forests to uptake carbon from the atmosphere through photosynthesis and tree growth in the future.

### 2.7 Uncertainty and alternative scenarios

To account for the uncertainty around the mean annual deforested area in our predictions, we computed the 95% confidence interval of the annual deforested area for each study area considering the deforestation observations in the period 2010–2020 (*SI Appendix*, Table S19). We thus obtained three different predictions of the forest cover change and associated carbon emissions: an average prediction considering the mean annual deforested area, and two additional predictions considering the lower and upper bound estimates of the mean annual deforested area per study area (*SI Appendix*, Figs. S14–S15, and Data S1, S2).

### 2.8 Software

To perform the analyses, we used the forestatrisk Python package (Vieilledent 2021) which has been specifically developed to model and forecast deforestation at high resolution on large spatial scales (*SI Appendix*, Materials and Methods).

## 3 Results

### 3.1 Model performance

Results of the cross-validation showed that the iCAR model had better predictive performance than the three other statistical models (*SI Appendix*, Tables S12). In particular, the Random Forest model overfitted the data and was less performant at predicting the probability of deforestation at new sites than the iCAR model. The iCAR model increased the explained deviance from 39.3 to 53.3% in average in comparison with the simple generalized linear model. Environmental explanatory variables alone explained a relative small part of the spatial deforestation process. Including spatial random effects to account for unexplained residual spatial variability strongly improved model’s fit (+14.0% of deviance explained in average) and model predictive performance (+7.4% for the TSS for example). Similar results were obtained when comparing accuracy indices between models at the continental scale (*SI Appendix*, Tables S13).

### 3.2 Effectiveness of protected areas at reducing deforestation

We found that protected areas significantly reduced the risk of deforestation for 70 study areas out of 119 (59% of the study areas). These 70 study areas accounted for 88% of the tropical moist forest in 2010 (*SI Appendix*, Table S6). But, the magnitude of this effect was relatively low: on average, protected areas reduced the risk of deforestation by 34% (Figs. 1 and *SI Appendix*, Table S5). Also, the effect of protected areas was highly variable between regions (*SI Appendix*, Table S6). For 18 study areas with a forest cover greater than 1 Mha in 2010, the effect of protected areas in reducing deforestation was not significant. Moreover, for the 47 study-areas with a forest cover greater than 1 Mha in 2010 for which the effect of protected areas was significant, the decrease in the deforestation risk within protected areas was highly variable (standard deviation = 18.72%) going from 5% (for the Bahia state in Brazil) to 82% (for Malaysia).

**Figure 1:**
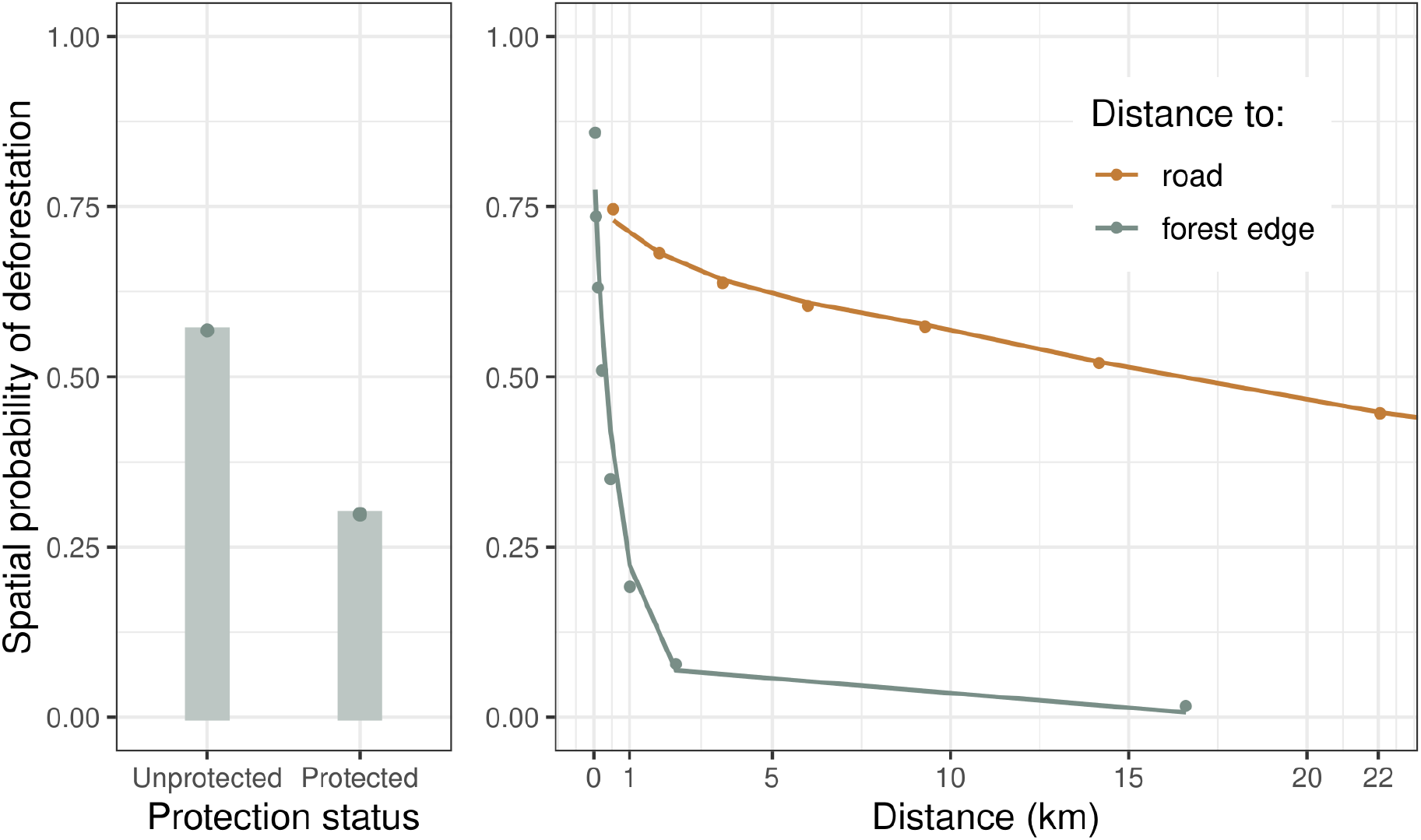
Effects of protected areas, roads, and distance to forest edge on the spatial probability of deforestation. For this figure, we used a representative dataset at the pantropical scale where the number of observations for each study area was proportional to its forest cover in 2010. We sampled 798,859 observations from the original dataset. *Left* : The dots represent the observed mean probability of deforestation in each forest protection class, either protected or unprotected. Bars represent the mean of the predicted probabilities of deforestation obtained from the deforestation model for all observations in each class. *Right* : The dots represent the local mean probability of deforestation for each bin of 10 percentiles for the distance. Lines represent the mean of the predicted probabilities of deforestation obtained from the deforestation model for all observations in each bin. Note that for distance to forest edge, the first dot accounts for three bins while for distance to road, bins for a distance > 23 km are not shown. For both left and right panels, confidence intervals for predictions were too small to be represented because of the high number of observations per class and bin.

### 3.3 Effect of distances to road and forest edge on the deforestation risk

We found that a greater distance to the road significantly reduced the risk of deforestation in 61 study areas out of 119 (51% of the study areas). These 61 study areas accounted for 90% of the tropical moist forest in 2010 (*SI Appendix*, Table S7). On average, a distance of 10 km to the road reduced the risk of deforestation by 14% (Figs. 1 and *SI Appendix*, Tables S5, S9). But the distance to forest edge was by far more important in explaining the deforestation risk than the distance to road (Fig. 1). The distance to forest edge was the most important variable in determining the risk of deforestation (*SI Appendix*, Table S5). We estimated that, on average, a distance of 1 km to the forest edge reduced the risk of deforestation by 91%, and a distance of 10 km reduced the risk of deforestation by almost 100% (Figs. 1 and *SI Appendix*, Tables S5, S9).

### 3.4 High resolution pantropical map of the deforestation risk

We obtained high resolution (30 m) maps of the deforestation risk for the year 2020 for the 119 study-areas. Combining these maps, we obtained a pantropical map of the risk of deforestation (Fig. 3 and *SI Appendix*, Fig. S10). The effect of protected areas and the effects of distances to road and forest edge on the risk of deforestation were clearly visible when looking at the map at the country and regional scales (Fig. 3 and *SI Appendix*, Fig. S10). Also, hotspots of deforestation (areas with a higher risk of deforestation), corresponding to areas with intense deforestation in the past (Fig. *SI Appendix*, Fig. S2), were clearly identifiable on the map (Fig. *SI Appendix*, Fig. S10).

**Figure 2:**
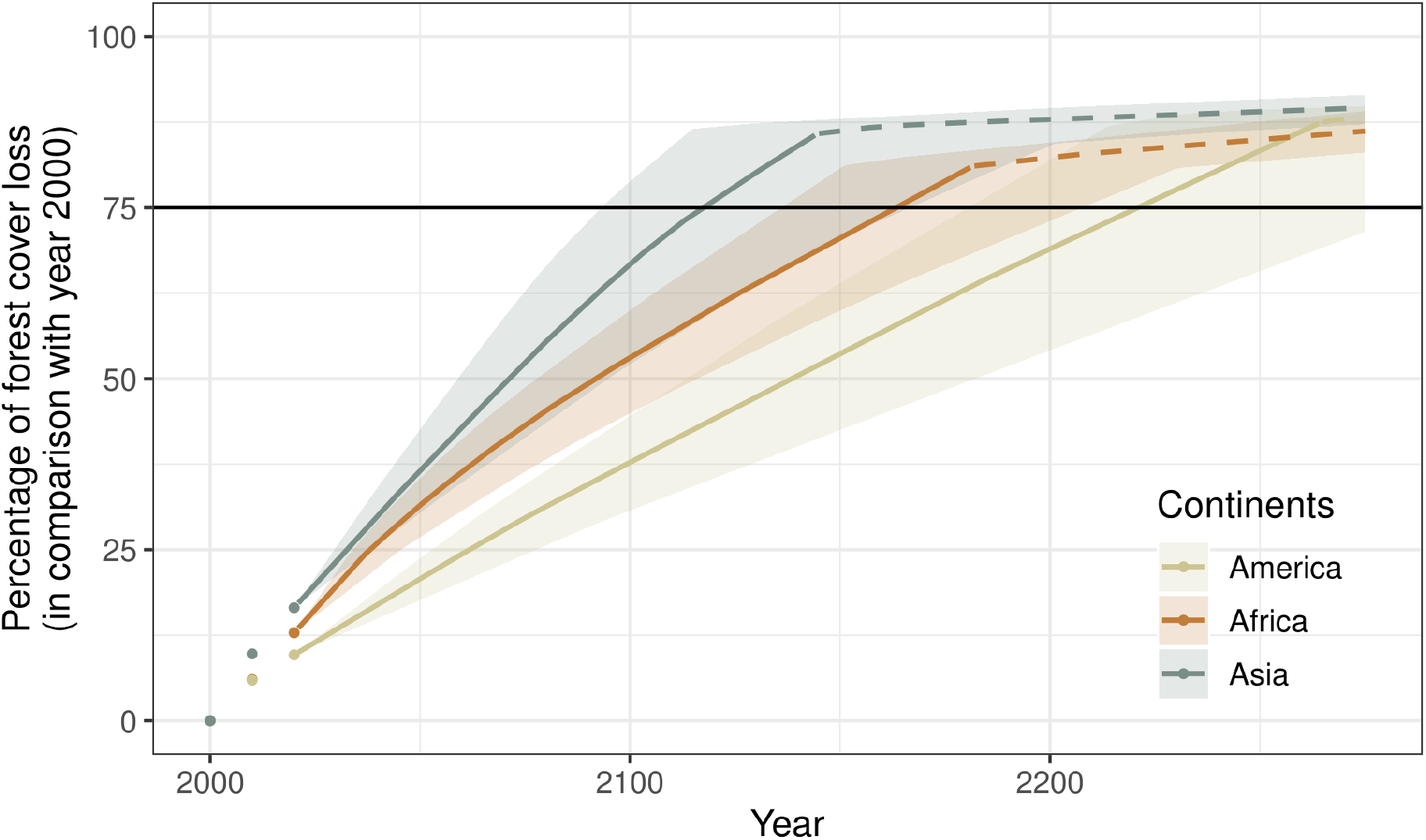
Projected forest cover loss per continent. Points represent the observed percentage of forest cover loss (in comparison with the year 2000) for the years 2000 (0%), 2010, and 2020, for America, Africa, and Asia. Lines represent the projected percentage of forest cover loss (in comparison with the year 2000) from 2020 to 2275 per continent. For the deforestation projections, we assumed no diffusion of the deforestation between countries. Under this assumption, deforestation at the continent scale is rapidly decreasing (dash lines) once large countries (Brazil for America, DRC for Africa, and Indonesia for Asia) have lost all their forests (ca. 2260, 2180, and 2140, respectively, see *SI Appendix*, Table S16). The horizontal black line indicates a loss of 75% of the continental forest cover. Under a business-as-usual scenario, this would happen in ca. 2120, 2160, and 2220 for Asia, Africa, and America, respectively. The confidence envelopes around the mean are obtained using the lower and upper bounds of the confidence intervals of the mean annual deforested areas for all study areas.

**Figure 3:**
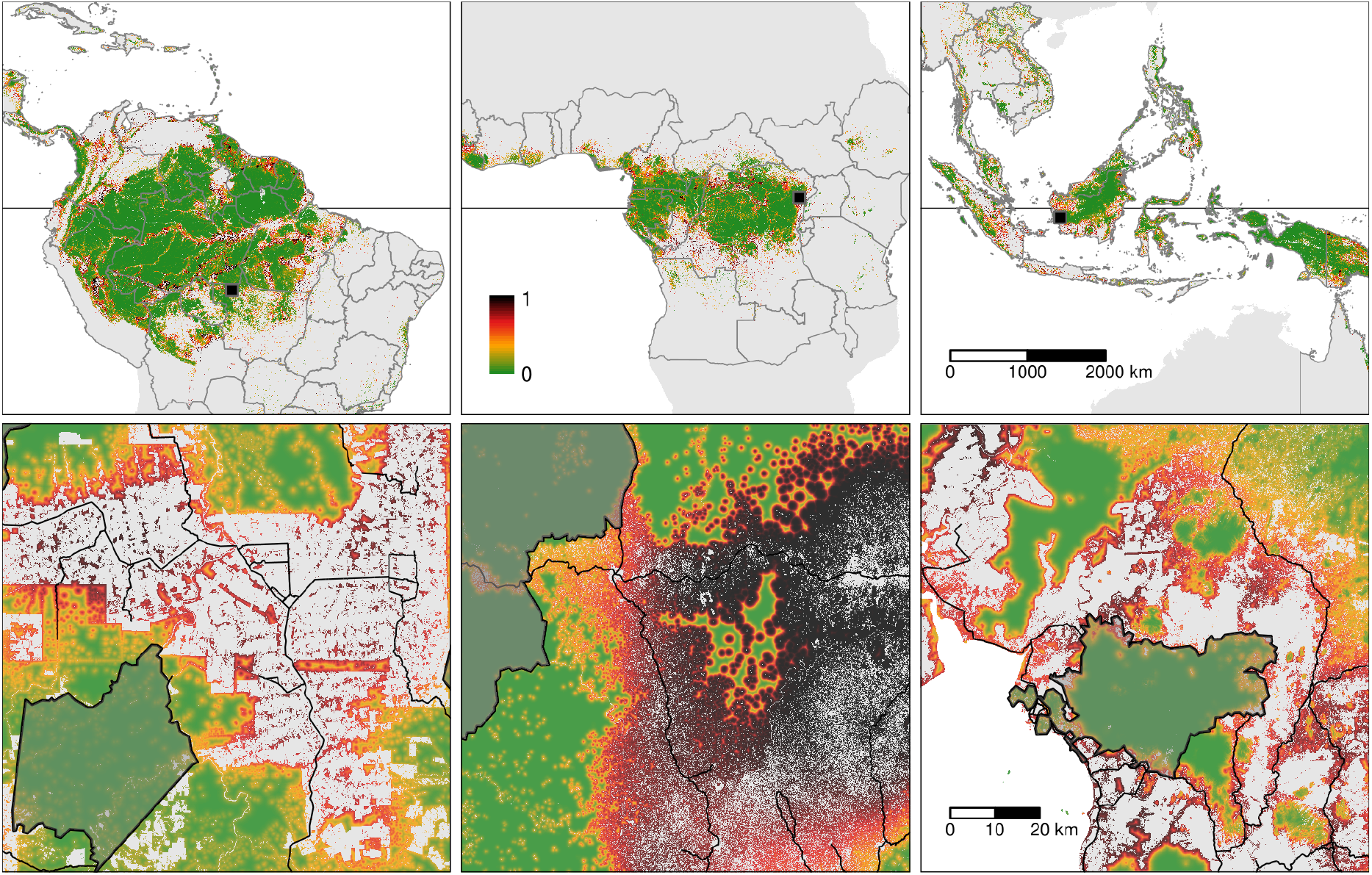
Pantropical map of the risk of deforestation. *Upper panels*: Maps of the spatial probability of deforestation at 30 m resolution for the three continents. Maps of the spatial probability of deforestation at the study area level were aggregated at the pantropical level. The horizontal black line represents the Equator. Study area boundaries are represented by dark grey lines. Coloured pixels represent forest pixels for the year 2020. Inside each study area, forest areas in dark red have a higher risk of deforestation than forest areas in green. *Lower panels*: Detailed maps for three 100 *×* 100 km regions (black squares in the upper panels) in the Mato Grosso state (Brazil), the Albertine Rift mountains (Democratic Republic of the Congo), and the West Kalimantan region (Borneo Indonesian part). Deforestation probability is lower inside protected areas (black shaded polygons) and increases when the forest is located at a distance closer to roads (dark grey lines) and forest edge. An interactive map of the spatial probability of deforestation is available at https://forestatrisk.cirad.fr/maps.html.

### 3.5 Forest cover change under a business-as-usual scenario of deforestation

We estimated that around 6.4 Mha (4.5–8.3 Mha) of tropical moist forest have been disappearing each year during the last decade (2010–2020) (Table 1). We show that under a business-as-usual scenario of deforestation, 48% (39–56%) of the world’s tropical moist forests will have disappeared over the course of the 21^st^ century (Table 1). We observed marked differences in the percentage of projected forest cover loss at continental and country scales (Fig. 2, Table 1, and *SI Appendix*, Tables S16–S17). The percentage of forest cover loss over the 21^st^ century would reach 67% (52–79%), 53% (45–60%), and 38% (31–45%) for Southeast Asia, Africa, and Latin America respectively (Table 1). Under a constant deforestation rate, three-quarters of the tropical moist forests that remained in 2000 will have disappeared around years 2120, 2160, and 2220 in Southeast Asia, Africa, and Latin America, respectively, with an average uncertainty of ±45 years (Fig. 2 and Table 1). At the country scale, we predicted that 41 countries (16 in Latin America, 21 in Africa, and four in Southeast Asia) out of the 92 we studied, plus 14 states in Brazil and one region in India, should lose all their tropical moist forests by 2100 (*SI Appendix*, Table S16).

**Table 1:**
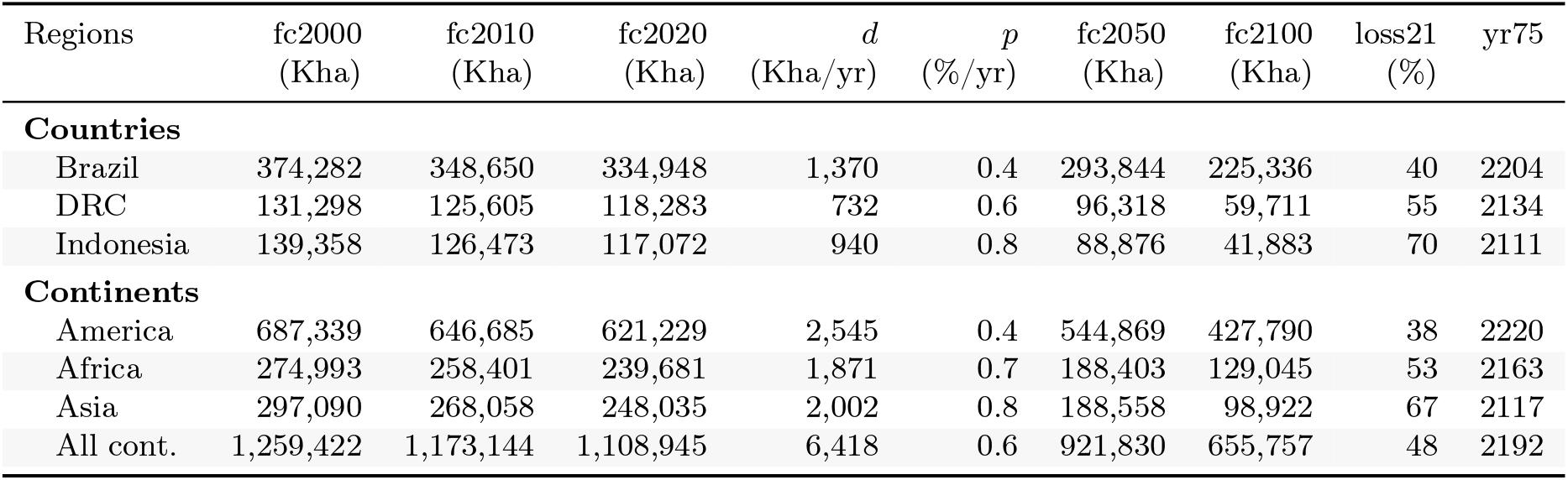
Past and predicted changes in forest cover. . We provide past and predicted forest cover for the three continents and for the three countries with the highest forest cover in 2010 for each continent (Brazil in America, the DRC in Africa, and Indonesia in Asia). Past forest cover areas (in thousand hectares, Kha) refers to their status on January 1^st^ 2000, 2010, and 2020 (“fc2000”, “fc2010”, and “fc2020”, respectively). We provide the mean annual deforested area *d* (Kha/yr) for the last ten-year period from January 1^st^ 2010 to January 1^st^ 2020, and the corresponding mean annual deforestation rate *p* (%/yr). Projected forest cover areas are given for the years 2050 and 2100 (“fc2050” and “fc2100”). Projections are based on the forest cover in 2020 (“fc2020”) and the mean annual deforested area (*d*) assuming a business-as-usual scenario of deforestation. Column “loss21” indicates the projected percentage of forest cover loss during the 21^st^ century (2100 vs. 2000). We estimate the year (“yr75”) at which 75% of the forest cover in 2000 will have disappeared.

### 3.6 Pantropical forest cover change maps for the 21^st^ century

Combining maps of the deforestation risk and the projected forest cover for years 2030, 2040, . . ., 2110 for each study area, we obtained pantropical forest cover change maps under a business-as-usual scenario of deforestation for the 21^st^ century in the humid tropics (Fig. 4 and *SI Appendix* and Fig. S11). Three large “blocks” of relatively intact tropical moist forest should remain in 2100 (Fig. 4) in the upper part of the Amazonian basin (including forests from Peru, Ecuador, Colombia, Venezuela, and the Guiana Shield), in the western part of the Congo basin (including forests from Gabon, Equatorial Guinea, Cameroon, the Central African Republic, and the Republic of Congo), and in Melanesia (including forests from Papua New Guinea, Solomon Islands, and Vanuatu). Apart from these three large and relatively intact forest blocks, the tropical moist forest remaining in 2100 should be highly fragmented (Fig. 4). In Africa for example, forests in the Democratic Republic of the Congo (DRC) will be highly fragmented (*SI Appendix*, Fig. S11) and completely separated from the large forest block located in the western part of the Congo basin (Fig. 4). The remaining forests will be concentrated in remote areas (far from roads and towns), preferentially in protected areas, and at high elevations. For example, the remaining forests of Borneo in 2100 will be concentrated in the Betung Kerihun and Kayan Mentarang National Parks (Figs. 3, 4, and *SI Appendix*, Tables S4–S9).

**Figure 4:**
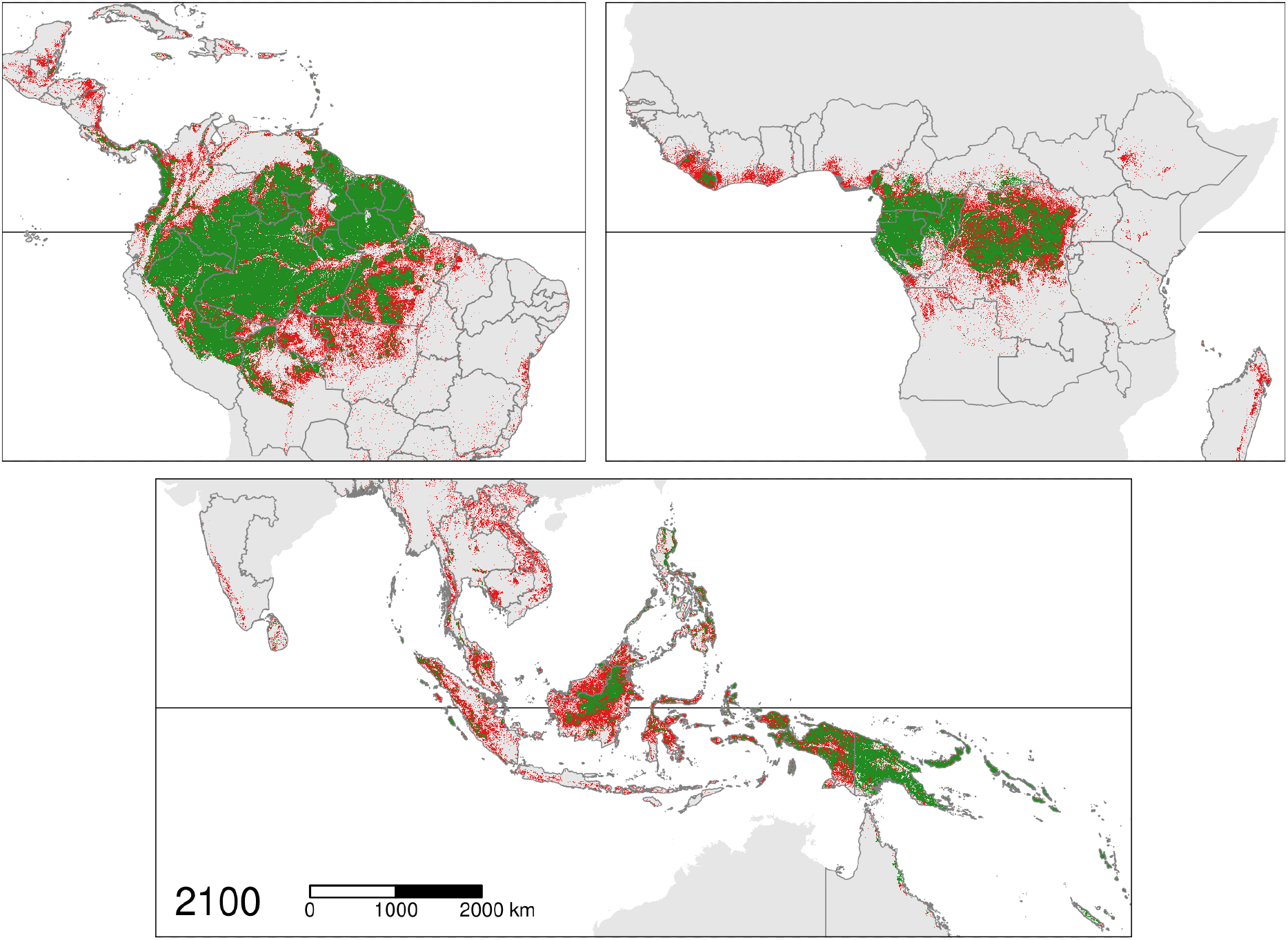
Pantropical map of the predicted change in forest cover. Maps show the predicted change in tropical moist forest cover in the three continents (America, Africa, and Asia) for the period 2020–2100 under a business-as-usual scenario of deforestation. The horizontal black line represents the Equator. Study area boundaries are represented by dark grey lines. For the deforestation projections, we assumed no diffusion of the deforestation between countries. Forest areas in red are predicted to be deforested in the period 2020–2100, while forest areas in green are likely to still exist in 2100. Several countries are expected to lose all their tropical moist forests by 2100 (including Nicaragua and Mexico in Central America, Madagascar and Ghana in Africa, and Laos and Vietnam in Asia). We predict progressive fragmentation of the remaining forest in the future, with an increasing number of isolated forest patches of smaller size (e.g., Pará state in Brazil, the Democratic Republic of the Congo, and Indonesia). These maps make it possible to identify both future hotspots of deforestation and forest refuge areas (e.g., concentrated in the heart of the Amazon, West Central Africa, and Papua New Guinea). An interactive map is available at https://forestatrisk.cirad.fr/maps.html.

### 3.7 Carbon emissions under a business-as-usual scenario of deforestation

Predictions obtained from both the ESA CCI and the WUR biomass maps suggested a substantial increase in carbon emissions associated with deforestation in the future, from 0.432 Pg/yr in 2010–2020 with the ESA CCI map (0.467 Pg/yr with the WUR map) to 0.590 Pg/yr in 2090–2100 (0.628 Pg/yr, respectively). This would correspond to a 27% (35%, respectively) increase in annual carbon emissions (Fig. 5 and *SI Appendix*, Fig. S13). Using either the ESA CCI or WUR maps, this increase in carbon emissions in the future was predicted for all three continents. This increase was not observed with the WHRC biomass map. In this case, we estimated that annual carbon emissions associated with deforestation at the global scale should remain stable throughout the 21^st^ century at about 0.600 Pg/yr (*SI Appendix*, Fig. S13). At the continental scale, whatever the biomass map we used, we predicted a decrease in annual carbon emissions starting from 2070–2080 for Southeast Asia which followed a period of increase in carbon emissions (Fig. 5 and *SI Appendix*, Fig. S13). Using annual rates of aboveground net biomass change for old-growth tropical rainforests and our estimates of forest cover change (Table 1), we estimated that the amount of carbon absorbed annually by tropical moist forests should drop by 47% (38–55%) during the 21^st^ century, from 0.589 Pg/yr in 2000 to 0.312 Pg/yr (0.267–0.363 Pg/yr) in 2100.

**Figure 5:**
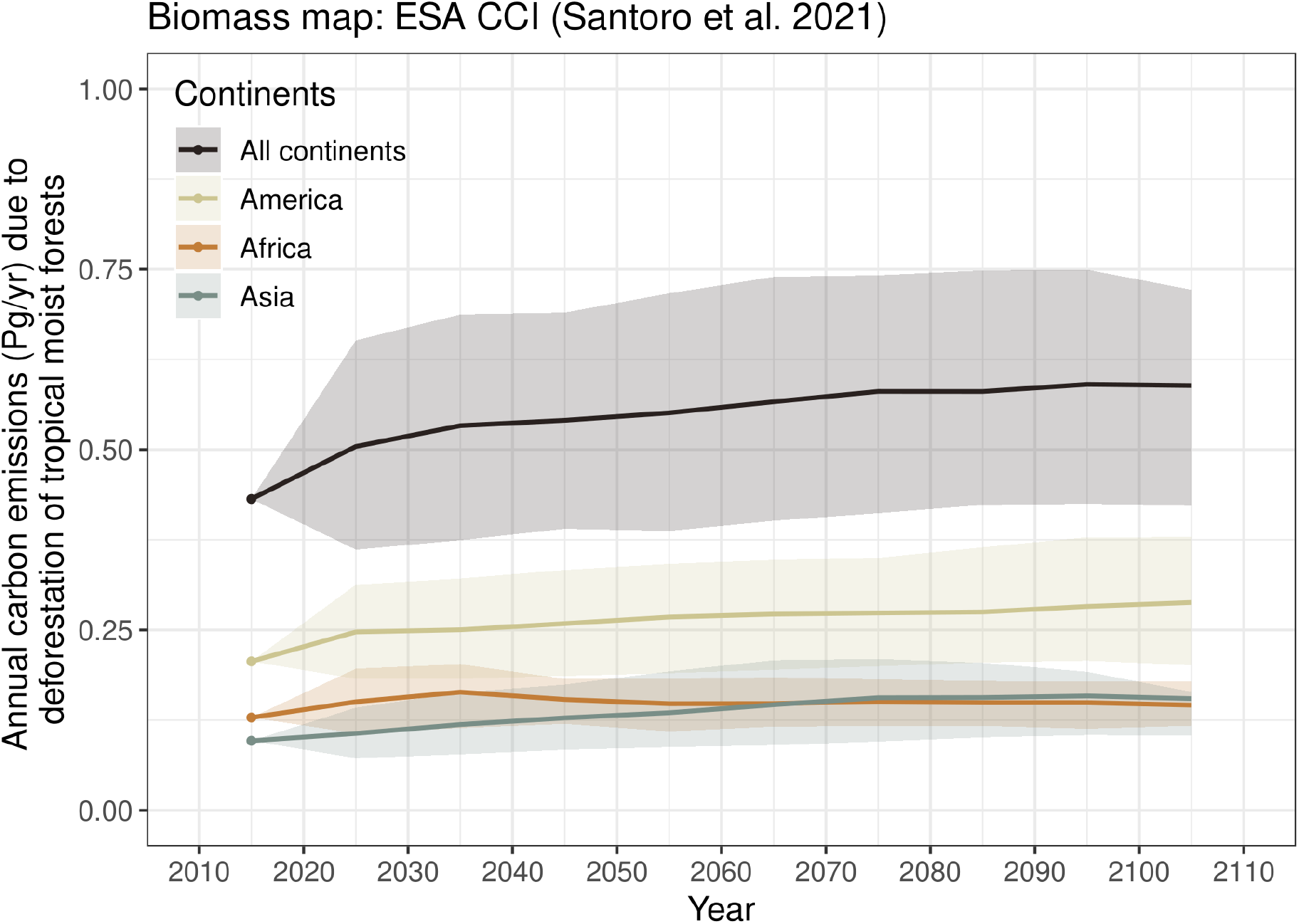
Annual carbon emissions associated with projected deforestation. Mean annual carbon emissions (Pg/yr) are estimated for ten-year intervals from 2010– 2020 to 2100–2110. The dots represent the observed mean annual carbon emissions (based on past deforestation maps) in 2010–2020, for the three continents (America, Africa, and Asia), and the three continents combined. Lines represent the projected mean annual carbon emissions based on projected forest cover change maps from 2020– 2030 to 2100–2110 per continent, and for all continents together. The confidence envelopes around the mean are obtained using the lower and upper bounds of the confidence intervals of the mean annual deforested areas for all study areas. The results shown here were obtained using the ESA CCI aboveground biomass map version 3. See *SI Appendix*, Fig. S13 for comparison with results obtained with other aboveground biomass maps.

## 4 Discussion

### 4.1 Reassessing the effects of protected areas and roads on the deforestation risk

Here we have shown that protected areas significantly reduced the risk of deforestation in 59% of the study areas, representing 88% of the tropical moist forest in 2010, but that the magnitude of this effect was relatively low (34%). In a recent global study, Wolf et al. (2021) estimated that deforestation was 41% lower inside protected areas, a value higher than the estimate in our study which was restricted to tropical moist forests. This means that protected areas do not prevent deforestation (deforestation does not stop at the boundaries of the protected areas) in the tropics and that the risk of deforestation is only reduced to some extent within protected areas. Moreover, our study has shown that the effectiveness of protected areas for reducing deforestation was very variable between study-areas, ranging from 0% to 82% reduction, and that the question of the effectiveness of protected areas must be preferentially answered on a case-by-case basis. Like several other studies reporting the effect of protected areas on deforestation (Andam et al. 2008, Wolf et al. 2021, Yang et al. 2021), our study shows that protected areas are effective on average in *displacing* deforestation outside protected areas in tropical countries, but not necessarily that protected areas play a role in *reducing* the deforestation intensity per se. Indeed, the factors that drive the intensity of deforestation at the country scale are more socio-economic or political, such as the level of economic development, which determines people’s livelihood and the link between people and deforestation (Geist and Lambin 2002), the size of the population (Barnes 1990), or the environmental policy (Soares-Filho et al. 2014). In tropical countries with weak governance (where environmental law enforcement is low) and with a low level of development (where the pressure on forests is high), it is very unlikely that protected areas will remain forested. Under a business-as-usual deforestation scenario, deforestation intensity is assumed constant over time. When all forests outside protected areas will have disappeared, deforestation is expected to occur inside protected areas (Fig. 4). In this scenario, protected areas are efficient at protecting forests with high conservation value in the medium term, i.e., forests will be concentrated in protected areas, where the probability of deforestation is lower. In the long term, forests should completely disappear from protected areas (Fig. 4). This phenomenon is already clearly visible in countries or states where deforestation is advanced, such as in Rondonia state (Brazil) in South America (Ribeiro et al. 2005), Ivory Coast (Sangne et al. 2015) and Madagascar (Vieilledent et al. 2020) in Africa, or Cambodia (Davis et al. 2015) in Southeast Asia. In these countries, several forested protected areas have been entirely deforested (e.g., the Haut-Sassandra protected forest in Ivory Coast, or the PK-32 Ranobe protected area in Madagascar) or severely deforested (e.g., the Beng Per wildlife sanctuary in Cambodia).

Regarding the effect of roads on the risk of deforestation, we have found that a distance of 10 km from a road reduces the risk of deforestation by 14% on average but that the distance to the forest edge was by far the most important variable in determining the risk of deforestation. On average, we found that a distance of 1 km from the forest edge reduced the risk of deforestation by 91%, in agreement with the results of other studies showing the importance of forest fragmentation on the risk of deforestation in the tropics (Hansen et al. 2020). Consequently, building new roads in non-forest areas but close to existing forest edges would significantly increase forest accessibility and the risk of deforestation in the nearby forest. But this negative impact would be greatly amplified if new roads were opened in the heart of forest areas. In addition to the direct deforestation associated with road building in the forest (Kleinschroth and Healey 2017), this would involve creating new forest edges and dramatically increase the deforestation risk in the area concerned. While road networks are expanding rapidly worldwide, notably in remote areas in tropical countries (Laurance et al. 2014), our results underline the importance of conserving large roadless and unfragmented forest areas.

### 4.2 Spatial scenarios of deforestation by 2100

We have estimated that around 6.4 Mha (4.5–8.3 Mha) of tropical moist forest have been disappearing each year during the last decade (2010–2020), which corresponds to an annual area of 64,000 km^2^, about the size of Greece or West Virginia. We have shown that under a business-as-usual scenario of deforestation, 48% (39–56%) of the world’s tropical moist forests should disappear over the course of the 21^st^ century. The percentage of lands covered by tropical moist forests would then decrease from 8.5% (1259 Mha) in 2000 to 4.7% (656 Mha) in 2100. Few studies have provided tropical forest cover projections for the 21^st^ century at the global scale. Using historical deforestation rates from Achard et al. (2002) and Malhi and Grace (2000) within the period 1990– 2000 (4.9–12.9 Mha/yr), Cramer et al. (2004) estimated that 46–80% of the tropical forests should disappear between 2000 and 2100 under a business-as-usual scenario. Considering historical deforestation rates for the period 1997–2002, Soares-Filho et al. (2006) estimated that the Amazonian forest would be reduced by 40% between 2003 and 2050 under a business-as-usual scenario. In our study, this percentage of loss for Latin America was not reached before 2100 (Fig. 2). These differences show first, that our predictions are rather conservative, and second, that predictions depend on the forest cover change product and on the reference period used to compute the historical deforestation rates. In our study, we used the most comprehensive and accurate forest cover change data available to date for moist tropical forests (Vancutsem et al. 2021) and performed an uncertainty analysis to see how differences in historical deforestation rates impacted forest cover change and carbon emission projections (Figs. 2, 5 and *SI Appendix*, Data S1, S2).

Despite the uncertainty surrounding the mean annual deforested area for each country (*SI Appendix*, Figs. S14–S15, and Table S19), the consequences of a business-as-usual deforestation scenario on the forest cover loss and associated carbon emissions by 2100 remain clear and alarming (Figs. 2, 5 and *SI Appendix*, Data S1, S2). Given the current global context, the business-as-usual deforestation scenario can be considered as conservative and might not be the most likely. For example, we do not account for the effect of future population growth (Raftery et al. 2012), which will likely have a major impact on deforestation, particularly in Africa, where a large part of the population depends on slash-and-burn agriculture for their livelihood (Barnes 1990, Vieilledent et al. 2020). Nor do we account for the increasing demand for agricultural commodities from the tropics, such as palm oil, beef and soybean, which will likely lead to a significant increase in deforestation (Karstensen et al. 2013, Strona et al. 2018). Our projections using high estimates of the annual deforested area for each study area, corresponding to a total deforestation of 8.3 Mha/yr at the pantropical scale, give an indication on the consequences of a 30% increase in the annual deforested area in the future. This would lead to a 56% loss of tropical moist forest cover over the 21^st^ century and the percentage of lands covered by tropical moist forests would then drop to 3.7% (554 Mha) in 2100 (*SI Appendix*, Fig. S15).

Our study provides estimates of forest cover change and associated carbon emission under a business-as-usual scenario for all countries having tropical moist forests. Our estimates are based on high resolution data (< 1 km) for both forest cover changes and carbon stocks. Previous studies providing scenarios of forest cover change and carbon emissions have generally worked at the continental scale (Cramer et al. 2004, Aguiar et al. 2016) or focused on particular countries (Soares-Filho et al. 2006) using data at coarser resolutions (*≥* 1 km). Using higher resolution data should provide more accurate results as both the deforestation risk and carbon stocks vary significantly over short distances (< 1 km, eg. effect of the distance to forest edge on the deforestation risk). Although the business-as-usual scenario might not be the most likely, it is of particular importance as it makes it possible to predict the likely future in the absence of change and can serve as a reference scenario for scenario comparisons. It is also used as the baseline scenario for the REDD+ mechanism (Goetz et al. 2015) which has been recognized as a major nature-based solution to fight climate change in the Paris Agreement on climate. The current methodology used to derive the baseline scenario in REDD+ projects has been intensively discussed recently (West et al. 2020, Guizar-Coutiño et al. 2022) in particular because of the risk of overestimating avoided deforestation and of displacement of deforestation outside the area of the project (also known as “leakage”). By providing maps of deforestation risk and forest cover change as well as carbon emissions at the jurisdictional level (national or subnational scale), our approach could help overcome these issues and provide a transparent and common methodology for implementing REDD+ at scale with more confidence.

### 4.3 Likely increase in carbon emissions under a business-as-usual scenario of deforestation

Using both the ESA CCI and the WUR maps, we predicted a substantial increase in carbon emissions associated with deforestation under a business-as-usual scenario, from 0.432–0.467 Pg/yr in 2010–2020 to 0.590–0.628 Pg/yr in 2090–2100. We obtained different results with the WHRC map annual for which carbon emissions should remain stable throughout the 21^st^ century at around 0.600 Pg/yr. Part of these differences could be explained by the fact that the WHRC map by Zarin et al. (2016) expands upon the methodology used for deriving the WHRC map by Baccini et al. (2012) which has shown higher overestimation of low biomass (*≤* 100 Mg/ha) and higher underestimation of high biomass (> 100 Mg/ha) than the ESA CCI and WUR maps (Vieilledent et al. 2016, Avitabile et al. 2016, Araza et al. 2022). These differences underline the importance of increasing the accuracy of global forest carbon maps, as underlined by previous studies (Ploton et al. 2020, Araza et al. 2022). Nonetheless, the estimates of 0.432–0.585 Pg/yr (range from the three maps) of aboveground carbon emissions due to tropical deforestation for 2010–2020 were consistent with those of several previous studies (Pan et al. 2011, Achard et al. 2014, Baccini et al. 2017, Harris et al. 2021). These studies estimated aboveground carbon emissions from tropical deforestation at 0.267– 0.669 Pg/yr within the period 2000–2019. The value of 0.432–0.585 Pg/yr from our study would represent 4.1–5.5% of the total anthropogenic carbon emissions estimated at 10.6 Pg/yr in 2011–2020 (Friedlingstein et al. 2021). Assuming a value of 0.30 for the belowground to aboveground biomass ratio (IPCC 2019), and that biomass represents 42% of the forest total carbon stock (including other compartments such as soil) (Pan et al. 2011), tropical deforestation would account for 12.7–17.0% of the total anthropogenic carbon emissions estimated in 2011–2020, in agreement with figures from previous studies (van der Werf et al. 2009, Harris et al. 2021, Friedlingstein et al. 2021).

Increase in carbon emissions predicted with the ESA CCI and WUR map is explained by the fact that future deforestation will concern forests with higher carbon stocks. Several studies have shown that elevation is an important variable in determining spatial variation of forest carbon stocks (Saatchi et al. 2011, Vieilledent et al. 2016, Cuní Sanchez et al. 2021). Forest carbon stocks are expected to be optimal at mid-elevation (Vieilledent et al. 2016) due to higher orographic precipitation at this elevation and because the climatic stress associated with winds and temperature is lower at mid-elevation than at high elevation. Here, we show that low-elevation areas have been more deforested than high-elevation areas (*SI Appendix*, Tables S4, S5). This can be explained by the fact that low-elevation areas are more accessible to human populations and by the fact that arable lands are concentrated at low elevation, where the terrain slope is usually lower and the soil is more productive (Geist and Lambin 2002). Consequently, the predicted increase in carbon emissions can be explained by the deforestation moving towards higher elevation areas where forest carbon stocks are higher. Moreover, remote forest areas less disturbed by human activities in the past have accumulated large quantities of carbon (Dargie et al. 2017, Brinck et al. 2017). The progressive deforestation of more intact forests could also explain the predicted increase in carbon emissions. We have also predicted a decrease in annual carbon emissions starting from ca. 2070 for Southeast Asia. This decrease could be explained partly by the lower carbon stocks of future deforested areas (driven by the environment, such as lower carbon stocks at very high elevation) and partly by a decrease in the total deforested area at the continental scale, as countries progressively lose all their forest. In Southeast Asia, we expect that four countries which account for a significant proportion of the annual deforested area in the continent in 2010–2020 (407,498 ha/yr out of 2,001,803 ha/yr, corresponding to 20% of the deforestation, see *SI Appendix*, Tables S14, S15) will lose all their forests between 2070 and 2110 (*SI Appendix*, Table S16). This likely explains most of the predicted decrease in carbon emissions in Southeast Asia from 2070 on.

Another consequence of tropical forest cover loss on the global carbon cycle is that the ability of tropical forests to uptake carbon from the atmosphere through photosynthesis and tree growth will decrease in the future. Using annual rates of aboveground net biomass change for old-growth tropical forests (Requena Suarez et al. 2019), we have estimated that the amount of carbon absorbed annually by tropical moist forests will drop by 47% (38–55%) during the 21^st^ century, from 0.589 Pg/yr in 2000 to 0.312 Pg/yr (0.267–0.363 Pg/yr) in 2100. These estimates are rather conservative as annual rates of carbon uptake by old-growth tropical forests are predicted to decrease in the 21^st^ century, in particular under the effect of climate change (Hubau et al. 2020). Recent studies have also shown that regrowth of tropical secondary forests could absorb large amounts of carbon from the atmosphere (Poorter et al. 2021, Heinrich et al. 2023). However, many secondary forests are not permanent and are cleared within a decade or two (Schwartz et al. 2020), so they cannot significantly contribute to carbon uptake from the atmosphere in the long term. Because carbon sequestration by tropical forests will not compensate for increasing carbon emissions from tropical deforestation (0.583– 0.628 Pg/yr in 2090–2100, range derived from the three biomass maps), tropical forests will likely act as an increasing net carbon source under a business-as-usual scenario of deforestation, thus reinforcing climate change in the future. If we consider a total deforestation of 8.3 Mha/yr at the pantropical scale, which gives an indication on the consequences of a 30% increase in the annual deforested area in the future, carbon absorption by tropical forests would drop down to 0.267 Pg/yr in 2100, and carbon emissions would increase up to 0.749–0.793 Pg/yr (Fig. 5 and *SI Appendix*, Fig. S13), turning tropical moist forests into an even bigger net carbon source. Our study provides new insights into the role of tropical forests in carbon emissions and climate change for the 21^st^ century. Although we have used lower annual deforestation rates for the projections than in previous studies (Cramer et al. 2004, Soares-Filho et al. 2006), tropical forests are likely to become a major net carbon source in the 21^st^ century both due to a decrease in forest cover, which will limit carbon uptake, and due to the deforestation of forests with higher carbon stocks in the future. These updated and forward-looking figures underline the importance of conserving tropical forests in the fight against climate change. The conservation of existing old-growth tropical forests is essential to avoid new carbon emissions and to maintain their function as a carbon sink.

## 7 Acknowledgments and Data

Our warm thanks to Rémy Dernat for help using the computing cluster of the Montpellier Bioinformatics Biodiversity (MBB) platform. We are also grateful to all the members of the Bioeconomy Unit at the JRC in Ispra for their kind support during this work. This research received fundings from the BioSceneMada project funded by FRB-FFEM (AAP-SCEN-2013 I), the Roadless Forest project funded by the European Commission, the RELIQUES project funded by CNRT, and the LabEx CeMEB funded by ANR “Investissements d’avenir” programme (ANR-10-LABX-04-01). The authors declare no conflicts of interest. *Author contribution*: GV and FA conceived the study, CV provided the forest cover change data, CB and PP helped to compute the carbon emissions, PV helped to write scripts for the HPC cluster, GV performed analysis and wrote the original draft, all authors reviewed and edited the final manuscript. *Data availability*: Raw data and products of the study are available on the ForestAtRisk website accompanying the present publication. Code is available on GitHub and is permanently archived in the Cirad Dataverse.

