## Supporting information for "Spatial scenario of tropical deforestation and carbon emissions for the 21^st^ century"

**This PDF file includes:**

Supplementary text for Materials and Methods
Figures S1 to S15
Tables S1 to S19
Legends for Data S1 and S2
SI References

**Other supplementary materials for this manuscript include the following:**

Data S1: Uncertainty around projected forest cover.
Data S2: Uncertainty around projected carbon emissions.

**5 SI References** **63**

### 1 Supplementary text for Materials and Methods

#### 1.1 Study areas

We defined 119 study areas representing 92 countries (Fig. S1) and covering all the tropical moist forest in the world, at the exception of some islands (eg. Sao Tome and Principe or Wallis-and-Futuna). Each country was identified by one unique three-letter code following the ISO 3166-1 standard (eg. MDG for Madagascar or GUF for French Guiana). Most of the countries corresponded to one unique study area, with three exceptions: Brazil, India, and Australia. Brazil, because of its large size, was divided into 26 study areas corresponding to the 26 administrative states (the state of Goias including the Federal District). For India, which is also a large country, the tropical moist forest is located in three distinct regions far from each other. We thus considered three independent study areas for India: the Western Ghats, North-East India (including the West Bengal), and the union territory of the Andaman and Nicobar Islands. For Australia, we only considered the Queensland state as a study area. Data sampling and spatial deforestation modelling were performed independently for each study area. Study area borders were obtained from version 3.6 of the Global Administrative Areas database (https://gadm.org). We used level-0 data for study areas corresponding to countries and level-1 data for study areas corresponding to states or regions. We grouped the study areas in three continents (Fig. S1): America (64 study areas for 39 countries), Africa (32 study areas for 32 countries), and Asia (23 study areas for 21 countries).

#### 1.2 Past forest cover change maps

For each study area, we derived past forest cover change maps for two time periods: January 1$^{st}$ 2000 – January 1$^{st}$ 2010, and January 1$^{st}$ 2010 – January 1$^{st}$ 2020 from the forest cover change annual product version v1_2020 by Vancutsem et al. (2021). The annual product by Vancutsem et al. (2021) classifies Landsat image pixels at 30 m resolution in 6 main categories (1: undisturbed, 2: degraded, 3: deforested, 4: regrowth, 5: water, and 6: other land cover) for each year (on the 31$^{st}$ of December) between 1990 and 2020 (31 years of data for v1_2020), and allows identifying tropical moist forest pixels at each date (Table S1). This classification is based on an expert system analyzing time-series data at the pixel level extracted from the full Landsat satellite image archive on the period 1982–2020. For our forest definition, we only considered *natural old-growth tropical moist forests*, disregarding plantations and regrowths. We included degraded forests (not yet deforested) in our forest definition. As a consequence, we considered all pixels falling in categories 1 and 2 in the annual product (Table S1), to be natural old-growth tropical moist forest pixels (simply abbreviated "forest" in this manuscript). Because several decades are usually necessary to reach the state of old-growth forest, we assumed every pixel classified as "forest" at a given date between 2000 and 2020 to be also classified as "forest" in the previous years of that period of time. We thus obtained three forest cover maps for the dates January 1$^{st}$ 2000, January 1$^{st}$ 2010, and January 1$^{st}$ 2020. We combined these three maps to obtain high-resolution forest cover change maps in the periods 2000–2010–2020 at 30 m resolution in the humid tropics (Fig. S2). We used Google Earth Engine (Gorelick et al. 2017) to process the annual product by Vancutsem et al. (2021) and derive the past forest cover change map for each study area. An interactive forest cover change map for the humid tropics is available at https://forestatrisk.cirad.fr/maps.html.

We did not consider potential forest regrowth in our forest definition for three main reasons. First, throughout the humid tropics, forest regeneration involves much smaller areas than deforestation (Hansen et al. 2013, Vancutsem et al. 2021). Second, there is little evidence of natural forest regeneration in the long term in the tropics (Grouzis et al. 2001). This can be explained by several ecological processes following deforestation such as soil erosion (Grinand et al. 2017), and reduced seed bank due to fire-induced deforestation and soil loss (Grouzis et al. 2001).

Moreover, in areas where forest regeneration is ecologically possible, young forest regrowths are more easily re-burnt for agriculture and pasture (Schwartz et al. 2020, Vieilledent et al. 2020). Third, young secondary forests generally provide more limited ecosystem services compared to old-growth natural forests in terms of biodiversity (Gibson et al. 2011) and carbon storage (Blanc et al. 2009).

##### 1.3 Spatial explanatory variables

To explain the observed deforestation during the period 2010–2020, we considered a set of spatial explanatory variables (Fig. S3) describing: topography (altitude and slope), accessibility (distances to nearest road, town, and river), forest landscape (distance to forest edge), deforestation history (distance to past deforestation), and land conservation status (presence of a protected area). This set of variables were selected based on an *a priori* knowledge of the deforestation process (Brown and Pearce 1994, Geist and Lambin 2002, Vieilledent et al. 2013). For example, the risk of deforestation is supposed to decrease with the distance to road and forest edge (lower accessibility), to increase at lower elevation and slope (higher probability to find arable lands), and to decrease in protected areas (higher level of protection).

Elevation (in m) and slope (in degree) at 90 m resolution were obtained from the SRTM Digital Elevation Database version v4.1 (http://srtm.csi.cgiar.org/). Distances (in m) to nearest road, town and river at 150 m resolution were computed from the road, town and river networks which were obtained from the OpenStreetMap (OSM) project (https://www.openstreetmap.org/). OSM country data were downloaded from two websites: Geofabric (http://download.geofabrik.de/) and OpenStreetMap.fr (https://download.openstreetmap.fr/extracts/) depending on the availability of the data for each country. To obtain the road network in each country (Fig. S4), we considered the "motorway", "trunk", "primary", "secondary" and "tertiary" categories for the "highway" key in OSM. Our dataset included a total of 3,606,841 roads. To obtain the network of populated places in each country (that we simply call "towns" in the present study), we considered the "city", "town" and "village" categories for the "place" key in OSM. To obtain the river network, we considered the "river" and "canal" categories for the "waterway" key in OSM. For a more detailed description of each category, see the OSM wiki page (https://wiki.openstreetmap.org/wiki/Tags). OSM data have been downloaded in March 2021 for all countries. Distance to forest edge was computed at 30 m resolution from the forest cover map in 2010. Distance to past deforestation in 2010 was computed at 30 m resolution from the 2000–2010 forest cover change map. To minimize border effect for the computation of distance to forest edge and distance to past deforestation, a buffer of 10 km around each study area extent was considered. Data on protected areas (Fig. S5) were obtained from the World Database on Protected Areas (https://www.protectedplanet.net, UNEP-WCMC and IUCN (2020)) using the `pywdpa` Python package (https://pypi.org/project/pywdpa/). WDPA data have been downloaded in March 2021 for all countries. For the analyses, we only retained protected areas defined by at least one polygon (we removed all protected areas defined by a point) and which had the following status: "Designated", "Inscribed", "Established", or "Proposed" before January 1$^{st}$ 2010 (we removed all "Proposed" protected areas after that date). Data included protected areas of all IUCN categories (from Ia to VI) and of all types defined at the national level (e.g. National Parks, Reserves), even if the type and IUCN category were not reported. Our dataset included a total of 89,855 protected areas. Polygons representing protected areas were rasterized at 30 m resolution.

In total, we obtained eight spatial explanatory variables to model the spatial probability of deforestation. Characteristics of each explanatory variable are summarized in Table S2 and correlation between variables are available in Fig. S6. Using data at a resolution closest to the resolution of the forest cover change map (30 m) to model deforestation ensures the capture of fine spatial scale deforestation processes. In particular, we demonstrate in this study the preponderant effect of the distance to forest edge which acts at a distance much lower than 1 km on the risk of deforestation.

##### 1.4 Data sampling for spatial modelling of deforestation

With the spatial model of deforestation, our aim was to estimate the effects of a set of variables in determining the location of the deforestation (or "allocation" census Pontius and Millones (2011)) and compute the relative probability of deforestation for each forest pixel. With the spatial model, our objective was not to estimate the intensity of the deforestation (or "quantity" census Pontius and Millones (2011)), that could be expressed in %/year or in ha/year for example. A balanced sampling between deforested and non-deforested pixels is preferable in this case (Vieilledent et al. 2013, Dezécache et al. 2017, Valle et al. 2020). Because deforestation events are rare ($\approx$ 1 %/yr), a non-stratified random sampling would lead to very few observations of deforestation events, rendering difficult a good estimation of the effects of the explanatory variables. Stratified balanced sampling provided unbiased estimates of the model's parameters, except for the model's intercept (estimated average deforestation). Having a biased model intercept (which has the same value for all forest pixels) is not a problem as we are interested in estimating a *relative* probability of deforestation between forest pixels.

As a consequence, we performed a stratified balanced sampling between (i) forest pixels in 2010 which have been deforested on the period 2010–2020 ("deforested" pixels), and (ii) forest pixels in 2010 which have not been deforested on that period of time and which represent the remaining forest in 2020 ("non-deforested" pixels). Forest pixels in each category were sampled randomly (Fig. S7). To maximize the representativity of the data, the total number of forest pixels sampled in each study area for the year 2010 was chosen proportionally to the area of forest in 2010 in that study area (2000 points for 1 Mha of forest), with the condition that this number had to be between 20,000 (to be representative of the deforestation process) and 100,000 (to limit computation time). When, for a specific study area, the total number of pixels in one of the two categories (deforested vs. non-deforested pixels) was $\leq$ 10,000, all the pixels of that category were included in the sample. This could happen for study areas with low moist forest cover such as small islands (eg. Antigua and Barbuda). For each sampled pixel, we retrieved information regarding the eight computed explanatory variables at their original spatial resolution. When the information was not complete for a given pixel (eg. elevation and slope data missing for a forest pixel located close to the sea border), the observation was removed from the dataset. Missing information affected a minority of pixels. The global dataset included a total of 3,197,942 observations: 1,601,125 non-deforested pixels and 1,587,817 deforested pixels, corresponding to an area of 144,914 ha and 142,908 ha, respectively (Table S3).

##### 1.5 Spatial deforestation model

Using observations of forest cover change in the period 2010–2020, we modelled the spatial probability of deforestation as a function of the $n$ explanatory variables using a logistic regression. We considered the random variable $y_i$ which takes value 1 if the forest pixel $i$ was deforested in the period 2010–2020 and 0 if it was not. We assumed that $y_i$ follows a Bernoulli distribution of parameter $\theta_i$ (Eq. (S1)). In our model, $\theta_i$ represents the spatial relative probability of deforestation for pixel $i$. We assumed that $\theta_i$ is linked, through a logit function, to a linear combination of the explanatory variables $X_i\beta$, where $X_i$ is the vector of explanatory variables for pixel $i$, and $\beta$ is the vector of effects $[\beta_1, \ldots, \beta_n]$ associated with the $n$ variables. All the continuous explanatory variables were normalized before fitting the model. The model includes an intercept $\alpha$. To account for the residual spatial variation in the deforestation process, we included an additional random effect $\rho_{j(i)}$ for each spatial cell $j$ of a 10 $\times$ 10 km grid covering each study area (Fig. S8). This grid resolution was chosen in order to have a reasonable balance between a good representation of the spatial variability of the deforestation process and a limited number of parameters to estimate. A sampled forest pixel $i$ was associated with one cell $j$ and one random effect $\rho_{j(i)}$. We assumed that random effects were spatially autocorrelated through an intrinsic conditional autoregressive (iCAR) model (Besag et al. 1991). This model is denoted "icar" in subsequent sections and results. In an iCAR model, the random effect $\rho_j$ associated with cell $j$ depends on the values of the random effects $\rho_{j'}$ associated with neighbouring cells $j'$. In our case, the neighbouring cells are connected to the target cell $j$ through a common border or corner (cells defined by the "king move" in chess, see Fig. S8). The variance of the spatial random effects $\rho_j$ was denoted $V_\rho$. The number of neighbouring cells for cell $j$, which might vary, was denoted $n_j$. Spatial random effects $\rho_j$ account for unmeasured or unmeasurable variables (Clark 2005) that explain a part of the residual spatial variation in the deforestation process that is not explained by the fixed spatial explanatory variables ($X_i$).

$$
\begin{aligned}
y_i &\sim \mathcal{B}ernoulli(\theta_i) \\
\text{logit}(\theta_i) &= \alpha + X_i\beta + \rho_{j(i)} \\
\rho_{j(i)} &\sim \mathcal{N}ormal(\sum_{j'} \rho_{j'}/n_j, V_\rho/n_j)
\end{aligned}
\tag{S1}
$$

#### 1.6 Variable selection

Variable selection was performed using a backward elimination procedure. All the spatial explanatory variables in our dataset should decrease the deforestation risk (having a negative effect on the probability of deforestation). For example, the probability of deforestation should decrease with the distance to the forest edge and should also decrease inside a protected area. Our variable selection procedure was thus not based on statistical significance but on background knowledge regarding the deforestation process and on the interpretability of the variable effects (Heinze et al. 2018). For each study area, we started to fit a model with the full set of explanatory variables (eight variables). At each step of the procedure, we removed the variables having a positive effect on the probability of deforestation. This was done in order to avoid unrealistic predictions of the spatial probability of deforestation at the scale of the study area. For example, it is not realistic to observe, at the country scale, a decrease of the deforestation with the distance to forest edge (higher deforestation risk in the core of the forest compared with forest edge). This might happen in a particular context for a very specific region but is very unlikely at the country level. In most of the cases, when we found a positive effect for a given explanatory variable, it was non-significant (95% credible interval including zero). On the contrary, when we found a negative but non-significant effect for a given explanatory variable, we kept this variable in the model for the predictions. This effect, albeit non-significant, was interpretable given our background knowledge of the deforestation process and was relatively lower than the effects associated with the other variables.

#### 1.7 Parameter inference

Parameter inference was done in a hierarchical Bayesian framework. We used the function `model_binomial_iCAR()` from the `forestatrisk` Python package (Vieilledent 2021) for parameter inference. This function calls an adaptive Metropolis-within-Gibbs algorithm written in C for maximum computation speed. Non-informative priors were used for all parameters: $\alpha \sim \mathcal{N}ormal(\text{mean} = 0, \text{var} = 10^6)$, $\beta \sim \mathcal{N}ormal(\text{mean} = 0, \text{var} = 10^6)$, and $V_\rho \sim 1/\mathcal{G}amma(\text{shape} = 0.05, \text{rate} = 0.0005)$. During the variable selection procedure, we run a Markov Chain Monte Carlo (MCMC) of 2000 iterations, discarding the first 1000 iterations (burn-in phase). For the final model, we repeated the parameter inference using a longer MCMC of 10,000 iterations. We discarded the first 5000 iterations (burn-in phase), and we thinned the chain each 5 iterations (to reduce autocorrelation between samples). MCMC convergence was visually checked looking at MCMC traces and parameter posterior distributions. We obtained 1000 estimates for each parameter. We used these 1000 estimates to compute the mean and 95% credible interval of each parameter (Tables S4–S7). We also back-transformed the parameters using the mean and standard-deviation of each continuous variable for each study area. Doing so, we can use Eq. (S1) to compute the change in the probability of deforestation associated with a particular change in the explanatory variables, in their original units. The value of the intercept $\alpha$ of the model is affected by the back-transformation, but not the effect of protected areas nor the variance of the spatial random effects, which are left unchanged (Tables S8, S9).

#### 1.8 Model comparison

##### 1.8.1 Alternative models

We compared the performance of the "icar" model at predicting deforestation with three other models: a null model (denoted "null"), a simple generalized linear model ("glm"), and a random forest model ("rf"). The "null" model assumes that all the slope parameters have value zero (no effect of explanatory variables), and that all the spatial random effects have also value zero (no residual regional variability in the deforestation process). For the "null" model, the probability of deforestation is only determined by a mean intercept common to every forest pixel ($\text{logit}(\theta_i) = \alpha$). The simple "glm" is a logistic regression which does not include spatial random effects (no residual regional variability in the deforestation process). For the "glm" model, the probability of deforestation is only determined by the mean intercept $\alpha$ and the parameters $\beta$ associated with each explanatory variables ($\text{logit}(\theta_i) = \alpha + X_i\beta$). Using simple "glm" models is a commonly proposed approach for spatial modelling of deforestation (Ludeke et al. 1990, Soares-Filho et al. 2002, Mas et al. 2007, Rosa et al. 2014). The random forest model (Breiman 2001) is a machine learning approach using an ensemble of random classification trees (where both observations and features are chosen at random to build the classification trees) to predict the deforestation probability for a forest pixel. Random forest has been intensively used for species distribution modelling (Thuiller et al. 2009) and is now also commonly used for spatial modelling of deforestation (Zanella et al. 2017, Grinand et al. 2020). The "glm" and "rf" models were fitted using functions `LinearRegression` and `RandomForestClassifier` respectively, both available in the `scikit-learn` Python package (Pedregosa et al. 2011). We used the same set of selected explanatory variables for the "glm" and "rf" models as those used for the final "icar" model. For the "rf" model, we set the number of random classification trees to 500.

##### 1.8.2 Percentage of deviance explained

We computed the deviance $\mathcal{D}$ of the four models ("icar", "null", "glm", and "rf") with the formula $\mathcal{D} = -2\log\mathcal{L}$, $\mathcal{L}$ being the likelihood of the model, i.e. the probability of observing the data given the model and estimated parameters. The deviance is a measure of error and a model with a lower deviance fits better the data. We also considered the deviance of the "full" model (also called the saturated model) which has as many parameters as there are observations. When $y_i = 0$, the deforestation probability predicted by the full model is 0. When $y_i = 1$ the deforestation probability predicted by the full model is 1. The deviance of the full model is then equal to 0. Considering that the "null" model explained 0% of the deviance and the "full" model explains 100% of the deviance, we then computed the percentage of deviance explained by each of the three other models: "icar", "glm", and "rf".

##### 1.8.3 Cross-validation procedure

To compare the performance of the "icar", "glm", and "rf" models at predicting correctly the relative probability of deforestation on independent observations, we also performed a five-fold cross-validation procedure. We used 70% of the observations for the model training and 30% of the observations for the model validation. We used the fitted models to predict the deforestation probability of all the forest pixels of the validation dataset. To transform the deforestation probabilities into binary values, we identified the probability threshold respecting the percentage of deforested pixels in the validation dataset (e.g., the mode of the predicted probabilities for a percentage of 50% of deforested pixels). Consequently, the predicted number of deforested pixels was equal to the observed number of deforested pixels in the validation dataset. This implies that there was no "quantity disagreement" (*sensu* Pontius and Millones (2011)) for any of the three models in the cross-validation procedure. Through this cross-validation, we only compared the ability of the models to correctly identify the pixels to be deforested, given a particular mean deforestation rate. This corresponds to estimating the "allocation disagreement" (*sensu* Pontius and Millones (2011)) for each of the three models. Using model predictions and observations in the validation dataset, we computed several accuracy indices: the Area Under the ROC Curve (AUC), the Figure of Merit (FOM), the Overall Accuracy (OA), the Specificity (Spe), the Sensitivity (Sen), and the True Skill Statistics (TSS). A detailed description of these indices can be found in Pontius et al. (2008) (for the FOM) and Liu et al. (2011) (for all the other indices). Formulas used to compute these indices are presented in Tables S10, S11.

##### 1.8.4 Model selection

For all the study areas, we found that the percentage of deviance explained for the "rf" model was much higher than for the "icar" and "glm" models (Table S12), suggesting that the "rf" model was fitting much better the data than the two other models. Nevertheless, when looking at the results of the cross-validation procedure, the "rf" model had a lower accuracy in comparison with the "icar" model (Table S12). These two results show clearly that the "rf" model was overfitting the data and was less performant at predicting the probability of deforestation at new sites than the "icar" model. This can be explained by the fact that the strong nonlinear relationship between explanatory variables and the spatial probability of deforestation, which is estimated by the "rf" model based on the training dataset, does not represent the true relationship at the landscape scale. This limitation associated with machine-learning techniques such as random forest has already been raised in previous scientific articles dealing with large scale mapping of ecological variables (Ploton et al. 2020). On the contrary, the "icar" model had the highest accuracy indices of the three models suggesting a better predictive performance than the "glm" and "rf" models. Also, the "icar" model increased the explained deviance from 39.3 to 53.3% in average in comparison with the "glm" model. This shows that environmental explanatory variables alone explain a relative small part of the spatial deforestation process and that including spatial random effects to account for unexplained residual spatial variability strongly improves model fit (+14.0% of deviance explained in average) and model predictive performance (+7.4% for the TSS for example). Same results were obtained when comparing accuracy indices for the three statistical models per continent (Table S13). We thus selected the "icar" model for predicting the spatial probability of deforestation for all the study areas.

#### 1.9 Computing the spatial probability of deforestation for the year 2020

Before computing the predictions of the deforestation probability, the spatial random effects at 10 km were interpolated at 1 km using a bicubic interpolation method (Fig. S9). This was done in order to obtain spatial random effects at a resolution closer to the original forest raster resolution of 30 m, and to smooth the deforestation probability predictions spatially.

Distance to forest edge in 2020 was recomputed from the forest cover map in 2020 at 30 m resolution. Distance to past deforestation in 2020 was recomputed from the 2010–2020 forest cover change map at 30 m resolution. All other explanatory variables (protected areas, distance to nearest road, town and river, elevation, and slope) were supposed unchanged between years 2010 and 2020. Using rasters of explanatory variables at their original resolution, interpolated spatial random effects at 1 km resolution, and the fitted "icar" model for each study area, we computed the spatial probability of deforestation at 30 m resolution for the year 2020 for each study area.

Deforestation probabilities (float values in the interval $[0, 1]$) were rescaled and transformed to integer values in the interval $[\![1, 65535]\!]$. This allowed us to record the large rasters of probabilities as UInt16 type (using zero as no-data value) and save space on disk. We then obtained a map of the relative probability of deforestation for the year 2020 at 30 m resolution (Fig. S10). An interactive global map of the spatial probability of deforestation is available at https://forestatrisk.cirad.fr/maps.html.

##### 1.10 Forecasting forest cover change for the period 2020–2110

For each study area, we computed the observed mean annual deforested area $d$ (in ha/yr) on the recent ten-year period 2010–2020 (from January 1$^{\text{st}}$ to January 1$^{\text{st}}$) using the forest cover change annual product by Vancutsem et al. (2021) (Tables S14, S15). The year 2020 was not considered to compute the recent mean annual deforested area as more disturbance events need to be observed on Landsat satellite images to disentangle degradation from deforestation for that year. As a consequence, most disturbance events are classified as degradation, and deforestation is underestimated for that specific year.

To forecast the forest cover change at a particular date $y$ in the future, we computed the estimated total deforestation $D_y$ (in ha) between years 2020 and $y$ assuming a "business-as-usual" scenario. The business-as-usual scenario makes the assumption of an absence of change in the deforestation intensity in the future (no increase in the deforestation intensity that could be attributable to a future increase in the demand of agricultural commodities for example, nor decrease in the deforestation intensity that could be attributable to new conservation policies or increase in agricultural yields for example). The business-as-usual scenario also makes the assumption that the spatial deforestation process will remain the same in the future (assuming a constant effect of the spatial explanatory variables in the future) and that areas with a higher relative probability of deforestation will remain the same in the future. To compute the total deforestation $D_y$ (in ha) between years 2020 and $y$, we projected the mean annual deforestation $d$ on the time interval $y - 2020$: $D_y = d \times (y - 2020)$.

For Brazil, which was divided into 26 study areas, the mean annual deforested area $d$ was supposed constant at the country level, not at the study area level. Indeed, we assumed that deforestation inside Brazil should spread between states and should not stop at the state administrative borders. As a consequence, when all the forest of a specific state in Brazil was deforested, the corresponding residual deforested area for that state was redistributed to the other states of Brazil still having forest. This ensured that the annual deforested area was constant for Brazil, but imply an increase of the annual deforested area with time for some states. We assumed that this contagious deforestation between study areas was only valuable for connected study areas inside a specific country (here, Brazil). On the contrary, we assumed that it was much less likely to observe contagious deforestation between two neighbouring countries. First, because of the presence of less permeable international borders, and second, because the socio-economic factors driving the intensity of deforestation between two neighbouring countries can be significantly different (see contrasting past deforestation intensity for Haiti–Dominican Republic, Congo–DRC, French Guiana–Suriname, or Indonesia–Papua New Guinea for example). We assumed no contagious deforestation for the three study areas in India which are not connected (Fig. S1).

The map of the relative spatial probability of deforestation in 2020 has a resolution of 30 m equivalent to an area of $r_{\text{ha}} = 0.09$ ha. Using this map, we computed a probability threshold $p_y$ in the interval $[\![1, 65535]\!]$ identifying the $n_y$ forest pixels in 2020 with the highest probability of deforestation so that $n_y \times r_{\text{ha}} = D_y + \epsilon$. Because deforestation probabilities have finite values in the interval $[\![1, 65535]\!]$, some forest pixels might have the same deforestation probability and it might not be possible to identify $p_y$ such that $\epsilon = 0$. We thus selected the threshold $p_y$ minimizing $\epsilon$. Because $D_y$ represents the total deforested area for several years and that few pixels had the same probability of deforestation, we always obtained negligible $\epsilon$ compared to $D_y$ ($\epsilon << D_y$). We considered those $n_y$ forest pixels in 2020 as deforested between years 2020 and $y$, and we derived the corresponding forest cover change map for the period 2020–$y$ (Fig. S11).

We projected the future forest cover for several years in the period 2030–2110 using a base time-interval of 10 years (Tables S16, S17). We also projected the future forest cover for years 2055 and 2085 as these two years are often used as pivot years in studies on future climate change. They correspond to 30-year climate averages on the periods 2040–2070 and 2070–2100, and to mid-term and long-term climate projections, respectively, for the 21$^{st}$ century (IPCC 2014). We then computed the percentage of forest cover loss in 2100 in comparison with the forest cover in 2000 for each study area and continent (Tables S16, S17). We also computed the year at which all the forest will have disappeared for each study area (Tables S16). At the continental level, it makes less sense to compute the year at which all the forest will have disappeared, as some countries might conserve forest for a very long time, even though they account for a very small proportion of the total forest area at the continental scale. Instead, we computed the estimated year at which 75% of the forest cover in 2000 will have disappeared (Table S17).

##### 1.11 Carbon emissions associated with deforestation

We estimated the carbon emissions associated with past deforestation (2010–2020) and projected deforestation (2030–2110) (Table S18) using three recent global or pantropical maps providing estimates of aboveground dry biomass (AGB, in Mg/ha) on a date before 2010 (Fig. S12).

The first map by Avitabile et al. (2016) from Wageningen University and Research (WUR) is a combination of two pantropical aboveground biomass maps by Saatchi et al. (2011) and Baccini et al. (2012). This map is representative of the aboveground biomass for the years 2000–2010 and provide AGB estimates at 1 km resolution. This fused map achieved a lower RMSE (87–98 Mg/ha representing a 5–74% decrease) and lower bias (a decrease of 90–153%) than the two input maps for all continents. This map was not covering some islands in the tropics: Mauritius, Reunion island, Fiji, New Caledonia, Solomon Islands, and Vanuatu. The second map by Zarin et al. (2016) from the Woods Hole Research Center (WHRC) provides AGB estimates at a higher resolution of 30 m, at pantropical scale, for circa the year 2000. This map expands upon the methodology presented in Baccini et al. (2012) using data from Landsat 7 ETM+ satellite imagery at 30 m resolution in place of MODIS data at 500 m resolution. The third map by Santoro et al. (2021), obtained within the framework of the Climate Change Initiative of the European Space Agency (ESA CCI), provides AGB estimates at 100 m resolution at the global scale for the year 2010. For this third map, and contrary to the two previous maps based on passive optical sensors, AGB estimates were retrieved from active radar sensors (ALOS-PALSAR and Envisat-ASAR sensors) for which the signal is not sensitive to weather conditions (eg. clouds often present over tropical moist forests).

We used the Intergovernmental Panel on Climate Change (IPCC) default carbon fraction of 0.47 (McGroddy et al. 2004) to convert aboveground dry biomass to carbon stocks. We assumed no change of the forest carbon stocks in the future while computing carbon emissions associated with projected deforestation. We estimated average annual carbon emissions for ten-year periods from 2010 to 2110. Under a "business-as-usual" scenario of deforestation (no change in the annual deforested area, in ha/yr, in the future), the change in mean annual carbon emissions in the future is only attributable to the spatial variation of the forest carbon stocks and to the location of the future deforestation. The three AGB maps were used separately to derive carbon emissions associated with deforestation and see if the changes in mean annual carbon emissions in the future were robust to change in the input data (Fig. S13).

#### 1.12 Uncertainty analysis

An uncertainty surrounds the estimate of the mean annual deforested area $d$ (in ha/yr) for each study area. We took into account this uncertainty in our predictions (Data S1 and S2). We computed the 95% confidence interval of $d$ for each study area using the deforestation observations $d_t$ for the ten years $t$ from 2010 to 2020 (Table S19). Deforestation observations $d_t$ were obtained from the forest cover change annual product by Vancutsem et al. (2021). We estimated $d'$, the lower limit of the confidence interval ($d' = \bar{d}_t - 1.96 \times \sigma(d_t)/\sqrt{n}$), and $d''$, the upper limit of the confidence interval ($d'' = \bar{d}_t + 1.96 \times \sigma(d_t)/\sqrt{n}$), with $\bar{d}_t$: the mean annual deforested area on 2010–2020 ($\bar{d}_t$=$d$), $\sigma(d_t)$: the standard deviation of the annual deforested area, and $n$: the number of observations ($n$=10, the number of years in our case). We repeated the simulations of the deforestation in the future using either $d'$ or $d''$ as the annual deforested area for each study area. We thus obtained three different predictions of the forest cover change and associated carbon emissions: an average prediction considering the mean annual deforested area $d$, a prediction considering a lower deforestation $d'$ (Fig. S14), and a prediction considering a higher deforestation $d''$ (Fig. S15).

#### 1.13 Software used: the `forestatrisk` Python package

We developed a specific package called `forestatrisk` (Vieilledent 2021) to model and forecast the tropical deforestation spatially using the Python programming language (Python Software Foundation 2020). The package has an associated website at https://ecology.ghislainv.fr/forestatrisk. The package is installable either from GitHub at https://github.com/ghislainv/forestatrisk or PyPI (The Python Package Index) at https://pypi.org/project/forestatrisk. It can be easily installed using `pip` (the package installer for Python) in any Python virtual environment created with either `conda` (recommended) or `virtualenv`. The `forestatrisk` package includes functions (i) to build a dataset from deforestation observations (function `.sample`), (ii) to estimate the parameters of several spatial deforestation models (functions `.model*`, including function `.model_binomial_iCAR` for the "icar" model), (iii) to assess the model performance (function `.cross_validation`), (iv) to derive predictive maps of the probability of deforestation (functions `.predict_raster*`), and (v) to forecast future forest cover under a given intensity of deforestation (function `.deforest`). Using functions from the `forestatrisk` Python package makes computation fast and efficient (with low memory usage) by treating large raster data by blocks. Numerical computations on blocks of data are performed with the NumPy (https://numpy.org) Python package whose core is mostly made of optimized and compiled C code which runs fast (Harris et al. 2020).

We also developed another smaller Python package called `pywdpa` (https://ecology.ghislainv.fr/pywdpa) that allows downloading the shapefiles of the protected areas for each country using the API of the World Database on Protected Areas (https://api.protectedplanet.net). The package is also available on GitHub at https://github.com/ghislainv/pywdpa or PyPI at https://pypi.org/project/pywdpa, and can also be easily installed using `pip`.

Computations for each country were run in parallel on the computing cluster of the Montpellier Bioinformatics Biodiversity (MBB) platform (https://mbb.univ-montp2.fr) provided by LabEx CeMEB (http://www.labex-cemeb.org/). Google Earth Engine (Gorelick et al. 2017) was used to process the annual product by Vancutsem et al. (2021) and derive the past forest cover change map for each country. While the raw results of our study (which implied intensive computations on large raster data) were obtained using the Python programming language, the summarized results (tables and figures) presented in this article (main text and supplementary materials) were obtained using the R software (R Core Team 2020).

#### 2 Supplementary figures

##### Fig. S1 – Study areas in the three continents

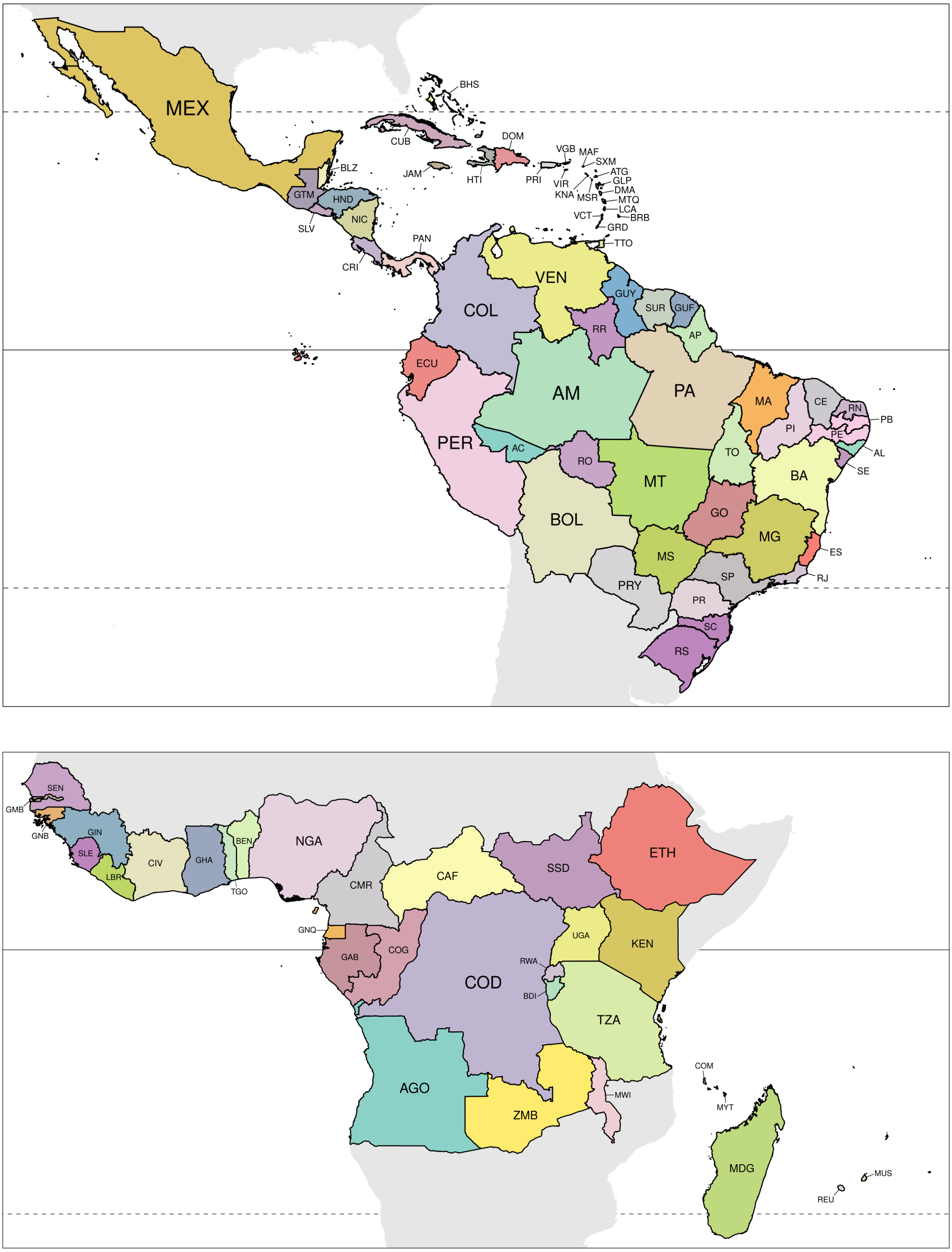

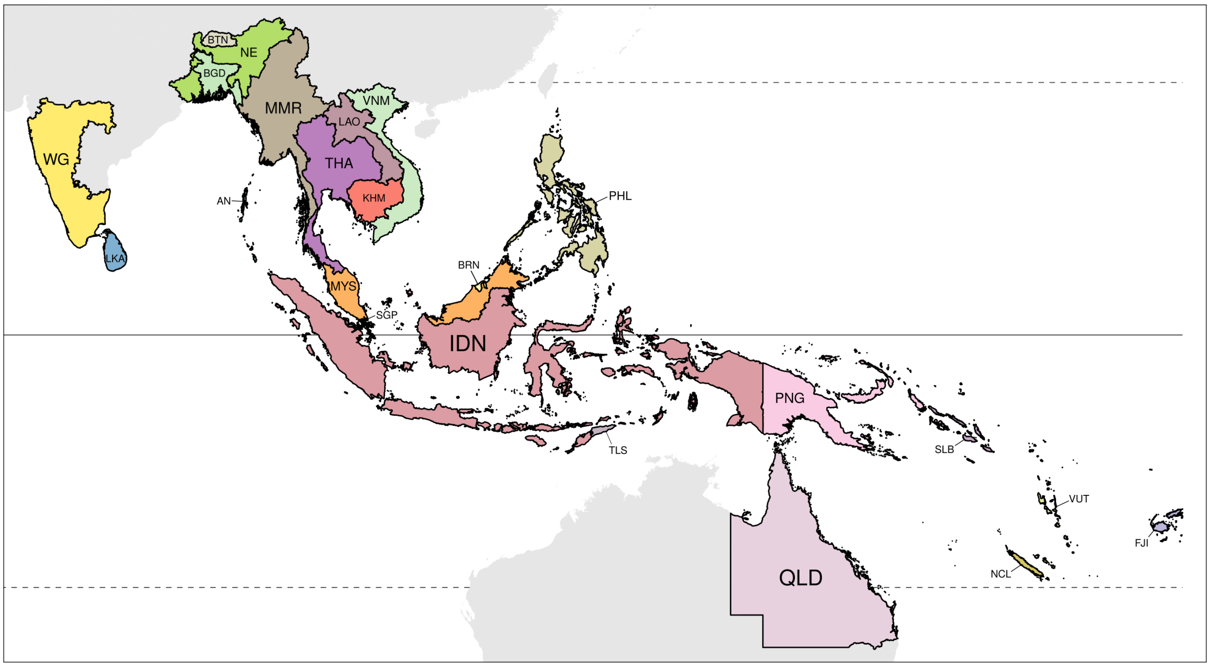

Figure S1: **Study areas in the three continents: America, Africa, and Asia**. America included 64 study areas (39 countries), Africa included 32 study areas (32 countries), and Asia included 23 study areas (21 countries). Each country was identified by one unique three-letter code following the ISO 3166-1 standard (eg. MDG for Madagascar or GUF for French Guiana). In America, Brazil was divided in 26 study areas corresponding to the 26 Brazilian states. Each Brazilian state was defined by one unique two-letter code (eg. AM for Amazonas). For India, three study areas were considered: the Whestern Ghats (WG), the North-East India (NE), and the Andaman and Nicobar Islands (AN). For Australia, we only considered the Queensland (QLD) state as a study area. In the three figures, each study area is identified by one unique code and a set of polygons with the same colour. The horizontal lines on each figure indicate the position of the Equator (plain line) and the two tropics (Cancer at the North and Capricorn at the South, dashed lines).

#### Fig. S2 – Past forest cover change map

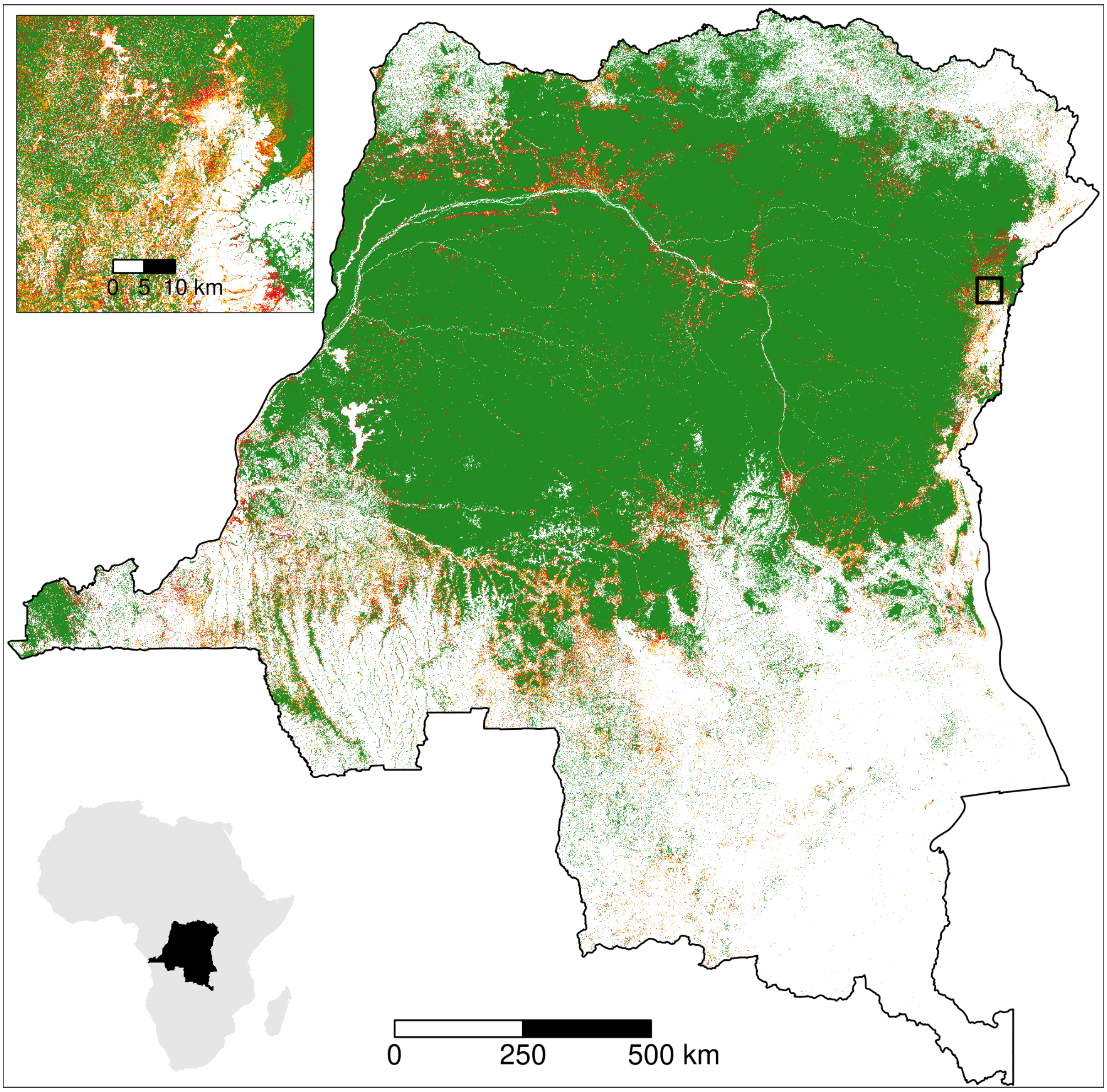

Figure S2: **Past forest cover change map**. Map of forest cover change for the period 2000–2010–2020 for the Democratic Republic of the Congo in central Africa (bottom-left inset). orange: 2000–2010 deforestation, red: 2010–2020 deforestation, green: forest cover in 2020. Forest cover change map was derived from the forest cover change annual product by Vancutsem et al. (2021). Original resolution of the forest cover change map is 30 m. The top-left inset shows a zoom of the map for an area at the North-East of the country which is close to the city of Beni and the Virunga national park. An interactive pantropical forest cover change map is available at https://forestatrisk.cirad.fr/maps.html.

#### Figs. S3–S6 – Spatial explanatory variables used for modelling deforestation

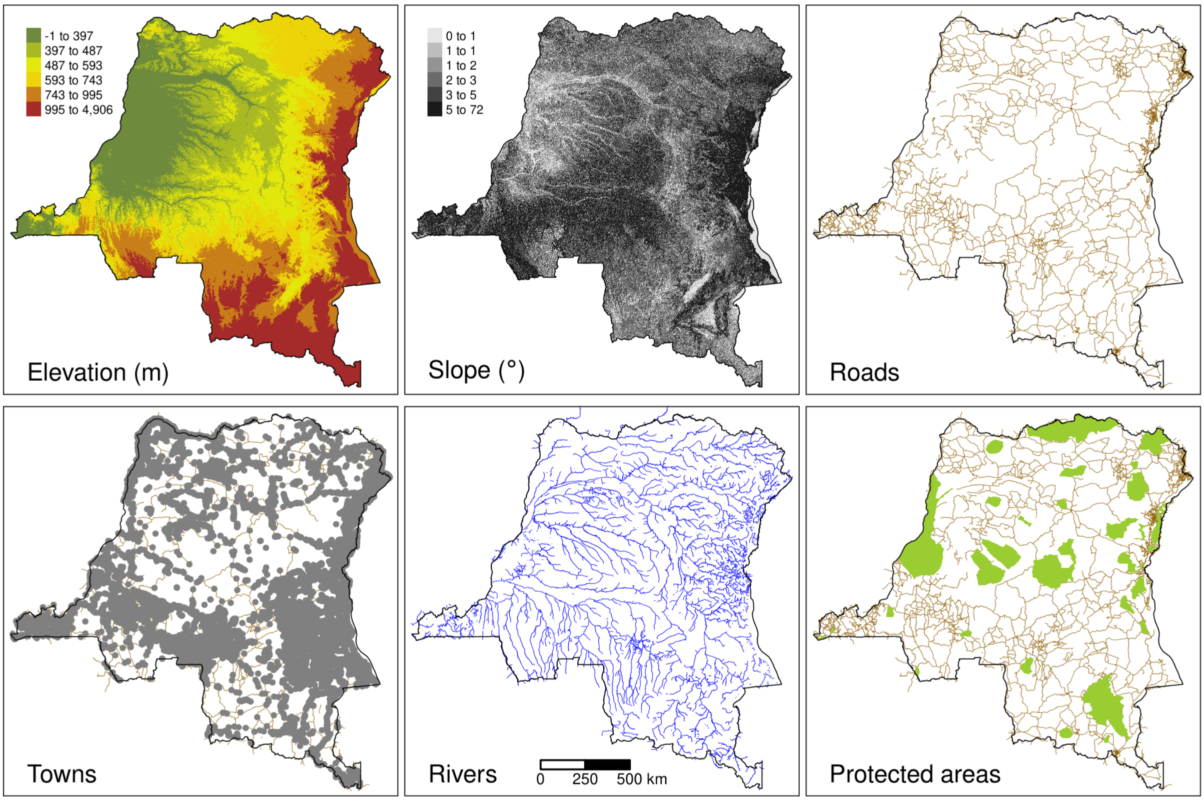

Figure S3: **Spatial explanatory variables**. Spatial explanatory variables for the Democratic Republic of the Congo in central Africa. Elevation (in m) and slope (in degree) at 90 m resolution were obtained from the SRTM Digital Elevation Database v4.1 (http://srtm.csi.cgiar.org/). Distances (in m) to nearest road, town, and river at 150 m resolution were computed from the road, town, and river network obtained from OpenStreetMap (OSM) (https://www.openstreetmap.org/). Roads include "motorway", "trunk", "primary", "secondary" and "tertiary" roads from OSM. Towns include "city", "town" and "village" categories from OSM. Rivers include "river" and "canal" categories from OSM. Protected areas were obtained from the World Database on Protected Areas (https://www.protectedplanet.net , UNEP-WCMC and IUCN (2020)). We retained protected areas defined by at least one polygon and which had the following status: "Designated", "Inscribed", "Established", or "Proposed" before January 1$^{st}$ 2010. Data included protected areas of all IUCN categories (from Ia to VI) and of all types defined at the national level (e.g. National Parks, Reserves). Two additional spatial explanatory variables (distance to forest edge and distance to past deforestation) were obtained from the past forest cover change map (Fig. S2).

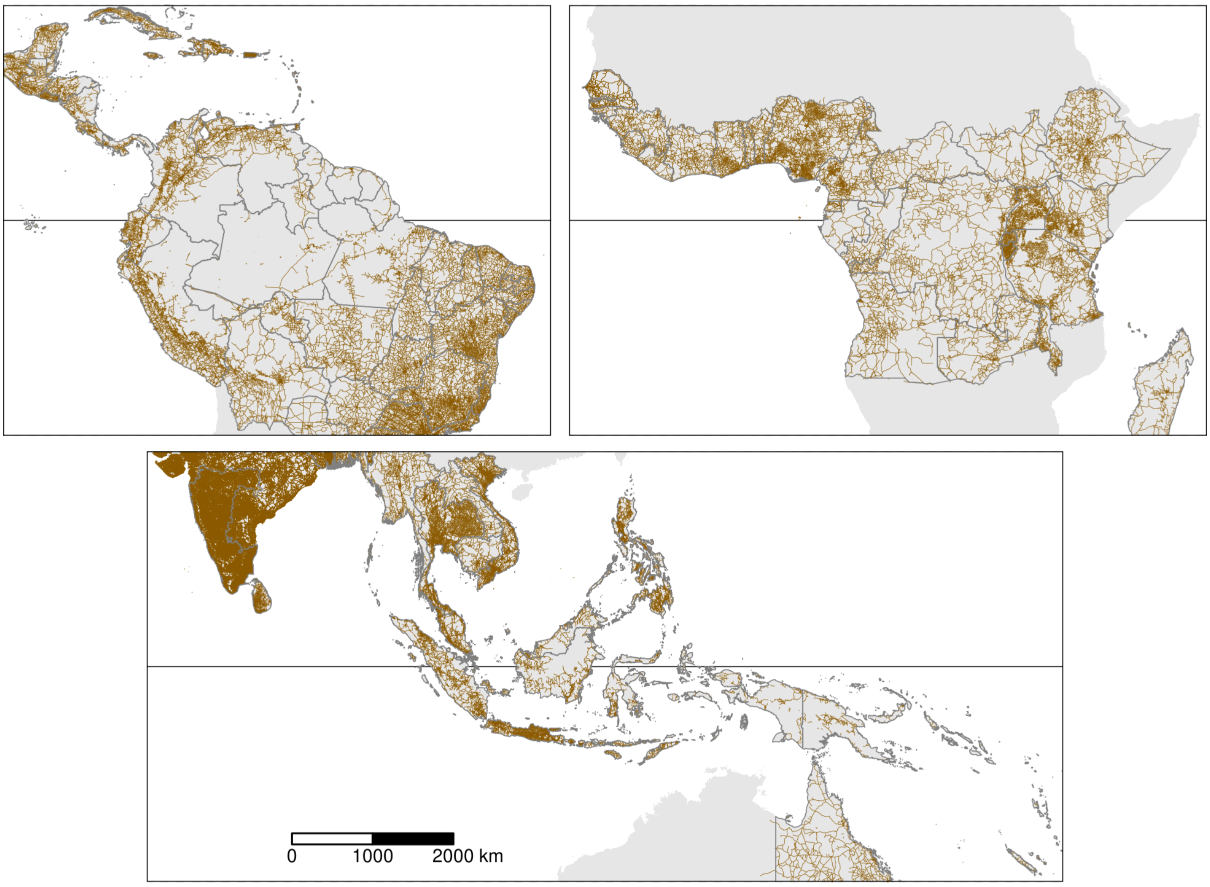

Figure S4: **Pantropical road network**. The road network was obtained from OpenStreetMap (OSM) (https://www.openstreetmap.org/). Roads included "motorway", "trunk", "primary", "secondary" and "tertiary" roads from OSM. Our dataset included a total of 3,606,841 roads.

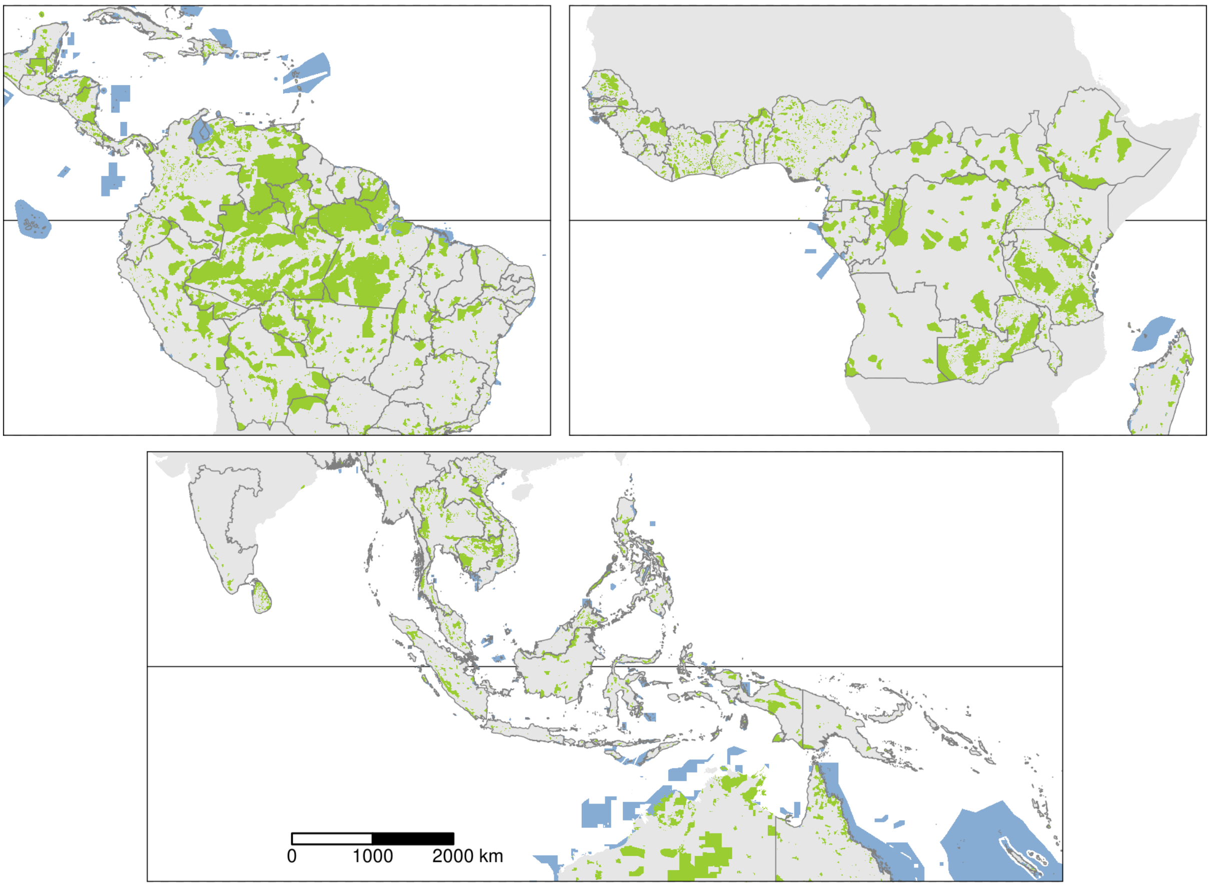

Figure S5: **Pantropical data-set on protected areas**. Terrestrial protected areas in green, marine protected areas in blue. Protected areas were downloaded from the World Database on Protected Areas (https://www.protectedplanet.net, UNEP-WCMC and IUCN (2020)) using the `pywdpa` Python package. We retained protected areas defined by at least one polygon and which had the following status: "Designated", "Inscribed", "Established", or "Proposed" before January 1$^{st}$ 2010. Data included protected areas of all IUCN categories (from Ia to VI) and of all types defined at the national level (e.g. National Parks, Reserves). Our dataset included a total of 89,855 protected areas.

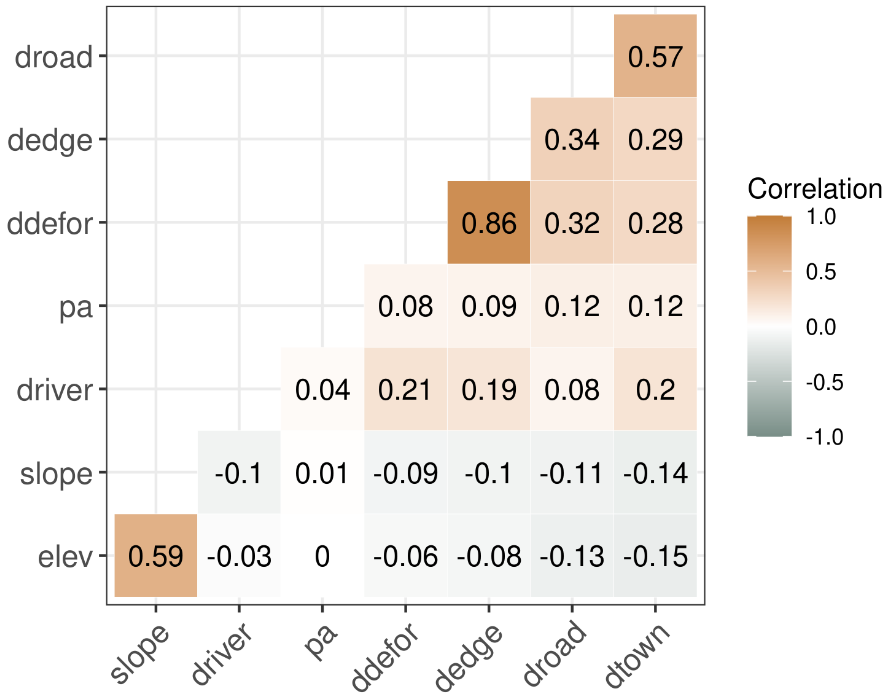

Figure S6: **Correlation between explanatory variables**. To compute the correlations, we used a representative data-set at the global scale where the number of observations for each study area was proportional to its forest cover in 2010. We used a total of 798,859 observations. We computed the Pearson's correlation matrix for the seven continuous explanatory variables used to model the deforestation: elevation ("elev"), slope ("slope"), distance to nearest road, town, and river ("droad", "dtown", and "driver", respectively), distance to forest edge ("dedge"), and distance to past deforestation ("ddefor"). For the protected areas ("pa"), which is a categorical variable for which a Pearson's correlation coefficient cannot be computed, we reported the slope coefficient of simple logistic regressions where the probability of presence of a protected area was a function of one intercept and one of the continuous variable (which was normalized).

#### Fig. S7 – Data sampling

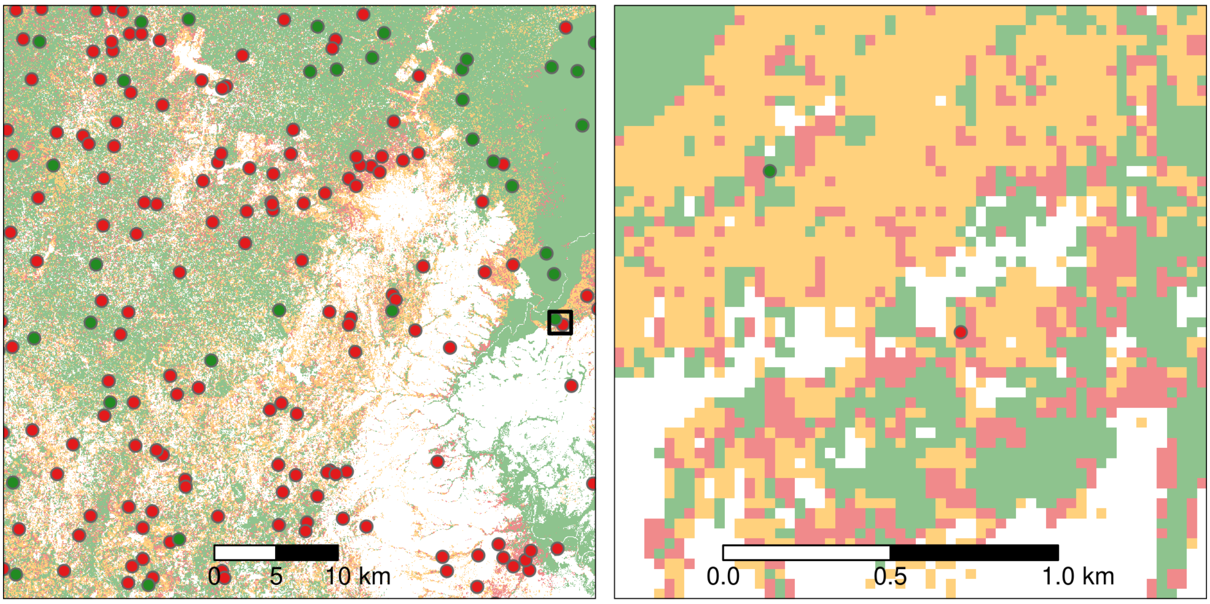

Figure S7: **Data sampling for spatial modelling of deforestation**. Map on the left corresponds to the top left inset in Fig. S2 representing a zoom of the forest cover change map in the period 2000–2010–2020 for an area at the North-East of the Democratic Republic of the Congo. Map on the right presents an inner zoom showing the delimitation of the 30 m forest pixels with two sample points. We used a stratified balanced sampling between (i) forest pixels in 2010 which have been deforested in the period 2010–2020 ("deforested" pixels in red), and (ii) forest pixels in 2010 which have not been deforested in that period of time and which represent the remaining forest in 2020 ("non-deforested" pixels in green). Forest pixels in each category were sampled randomly.

#### Fig. S8 – Grid for spatial random effects

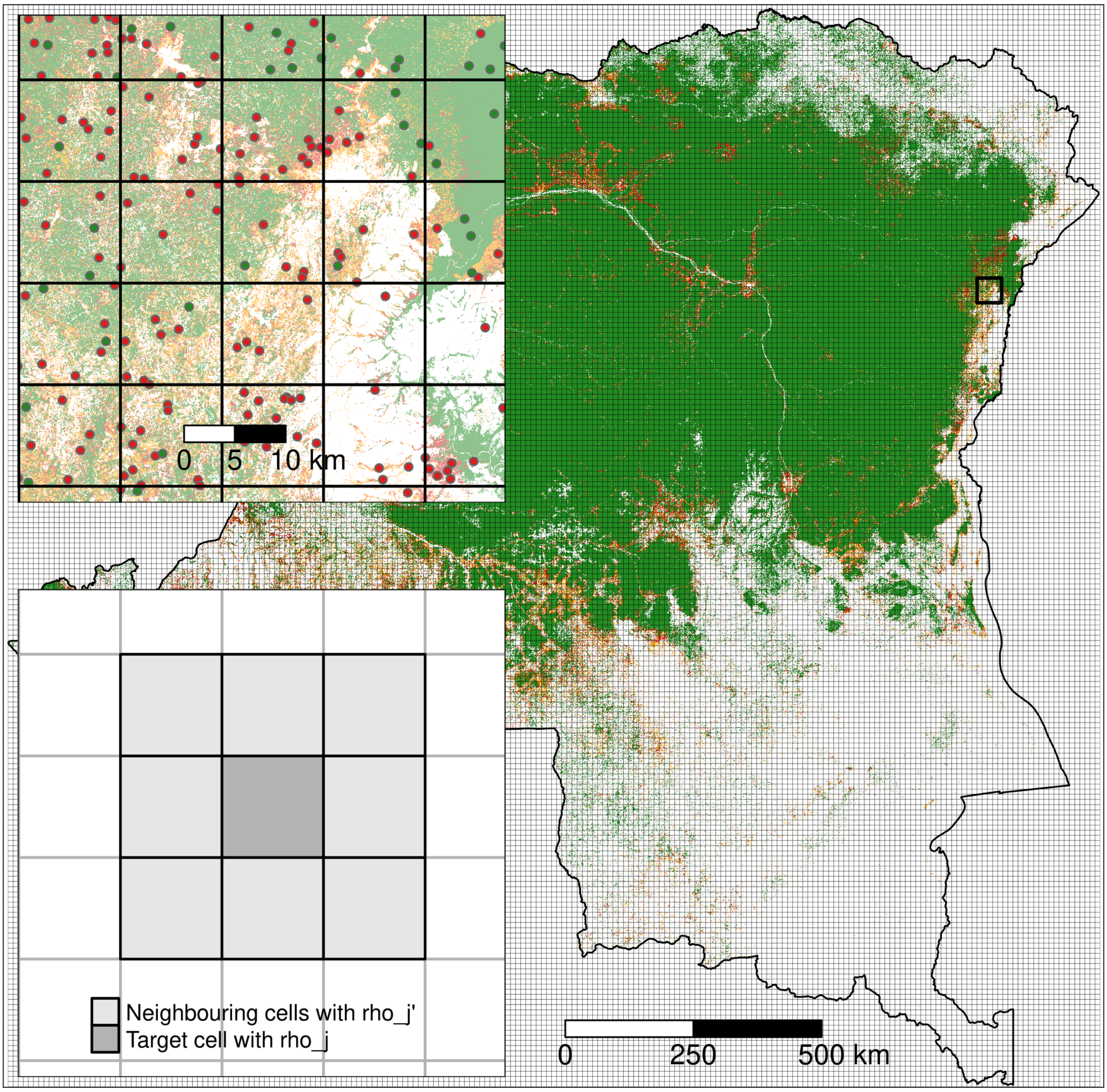

Figure S8: **Grid used to compute the spatial random effects**. *Main figure*: $10 \times 10$ km grid covering the Democratic Republic of the Congo (DRC). The grid over DRC includes 45,154 $10 \times 10$ km cells (214 cells on the $x$ axis by 211 cells on the $y$ axis). The background map shows the past forest cover change in the period 2000–2010–2020 (see Fig. S2). *Top inset*: Zoom for an area at the North-East of the country (black square) showing specific grid cells. One grid cell can include several sample points (see Fig. S7). *Bottom inset*: One random effect $\rho_j$ is estimated for each grid cell $j$. Spatial autocorrelation is taken into account through an intrinsic conditional autoregressive (iCAR) process: the value of the random effect for one cell depends on the values of the random effects $\rho_{j'}$ for the neighbouring cells $j'$ (see Eq. (S1)).

#### Fig. S9 – Estimated spatial random effects

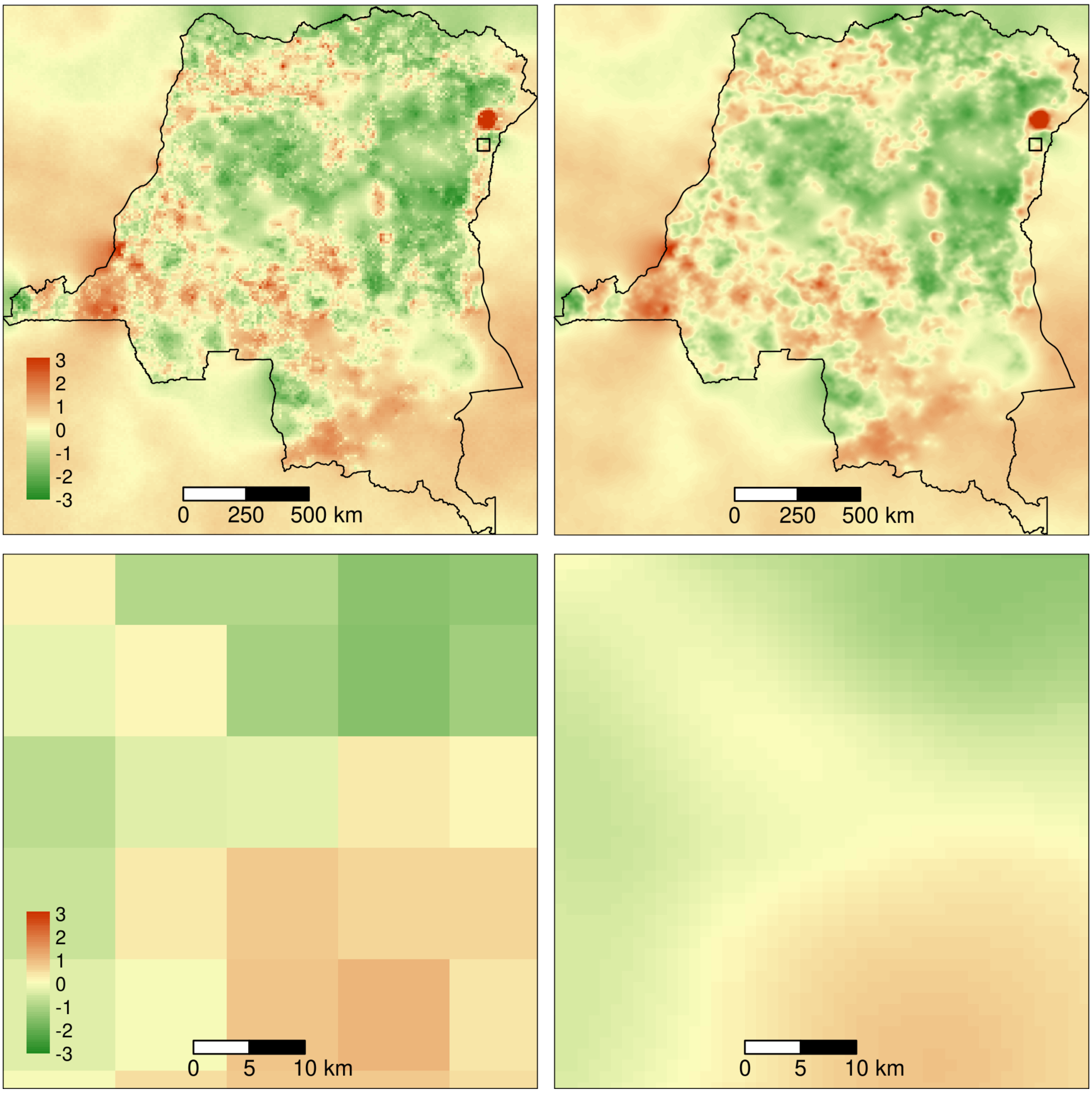

Figure S9: **Estimated spatial random effects**. *Left*: Estimated spatial random effects at 10 km resolution for the Democratic Republic of the Congo (DRC). *Right*: Interpolated spatial random effects at 1 km resolution. A bicubic interpolation method was used. *Bottom*: Zoom for an area at the North-East of the country (black square) which is close to the city of Beni and the Virunga national park. Due to the structure of the iCAR model (see Eq. (S1)), spatial random effects are also estimated for cells without sampled points. This includes cells for which there was no forest cover in the period 2000–2010–2020, and also cells outside the country's borders.

#### Fig. S10 – Spatial relative probability of deforestation

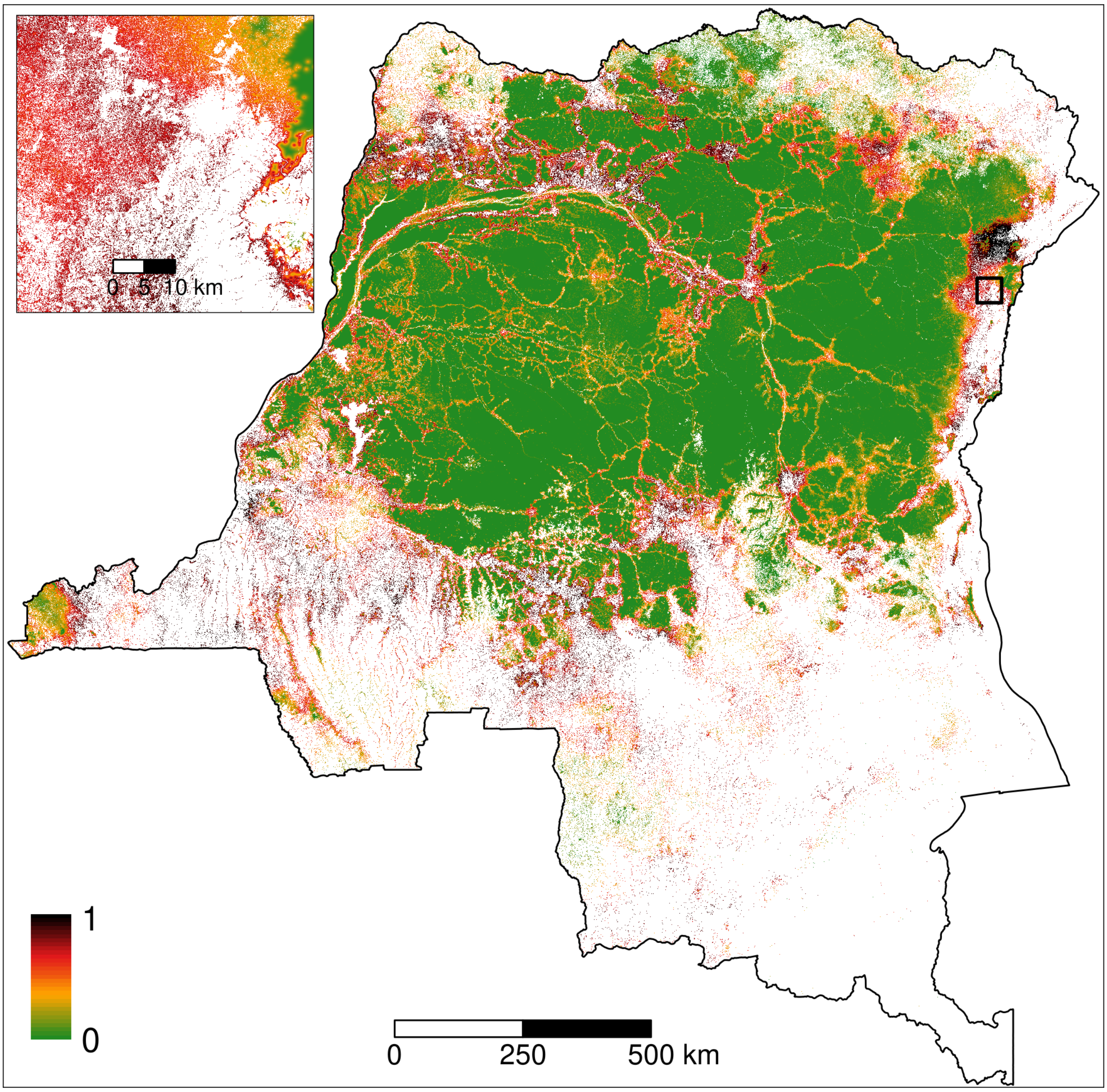

Figure S10: **Predicted spatial relative probability of deforestation**. *Main figure*: Map of the spatial probability of deforestation computed for each forest pixel in 2020 for the Democratic Republic of the Congo. On the map, we clearly see the effect of the distance to nearest town and road, and the effect of the distance to forest edge on the spatial probability of deforestation. Also, we clearly see the importance of the spatial random effects in structuring the spatial variability of the deforestation probability. For example, the area at the North of the zoom (black square) shows very high deforestation probabilities (in black). This area is politically unstable and is home to a large number of militias who survive at the expense of the forest. *Inset*: Zoom of the map for an area at the North-East of the country which is close to the city of Beni and the Virunga national park. An interactive global map of the spatial probability of deforestation is available at https://forestatrisk.cirad.fr/maps.html.

#### Fig. S11 – Projected forest cover change

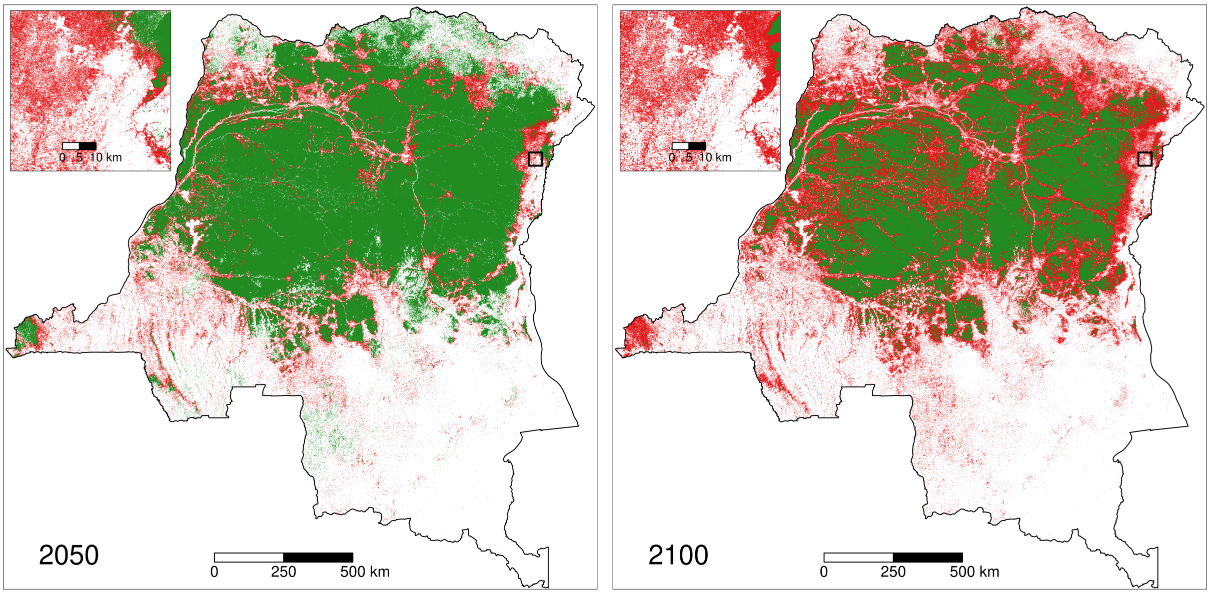

Figure S11: **Projected forest cover change**. *Main figures*: Maps of the projected forest cover change (left: 2020–2050, right: 2020–2100) for the Democratic Republic of the Congo (DRC) under a business-as-usual scenario of deforestation. red: projected deforestation, green: remaining forest cover. Besides the loss of forest cover, maps show a progressive fragmentation of the forest in the future, with an increasing number of forest patches of smaller size in DRC. *Insets*: Zoom of the map for an area at the North-East of the country (black square) which is close to the city of Beni and the Virunga national park. Interactive pantropical maps of the projected forest cover change for years 2050 and 2100 are available at https://forestatrisk.cirad.fr/maps.html.

#### Fig. S12 – Aboveground biomass maps

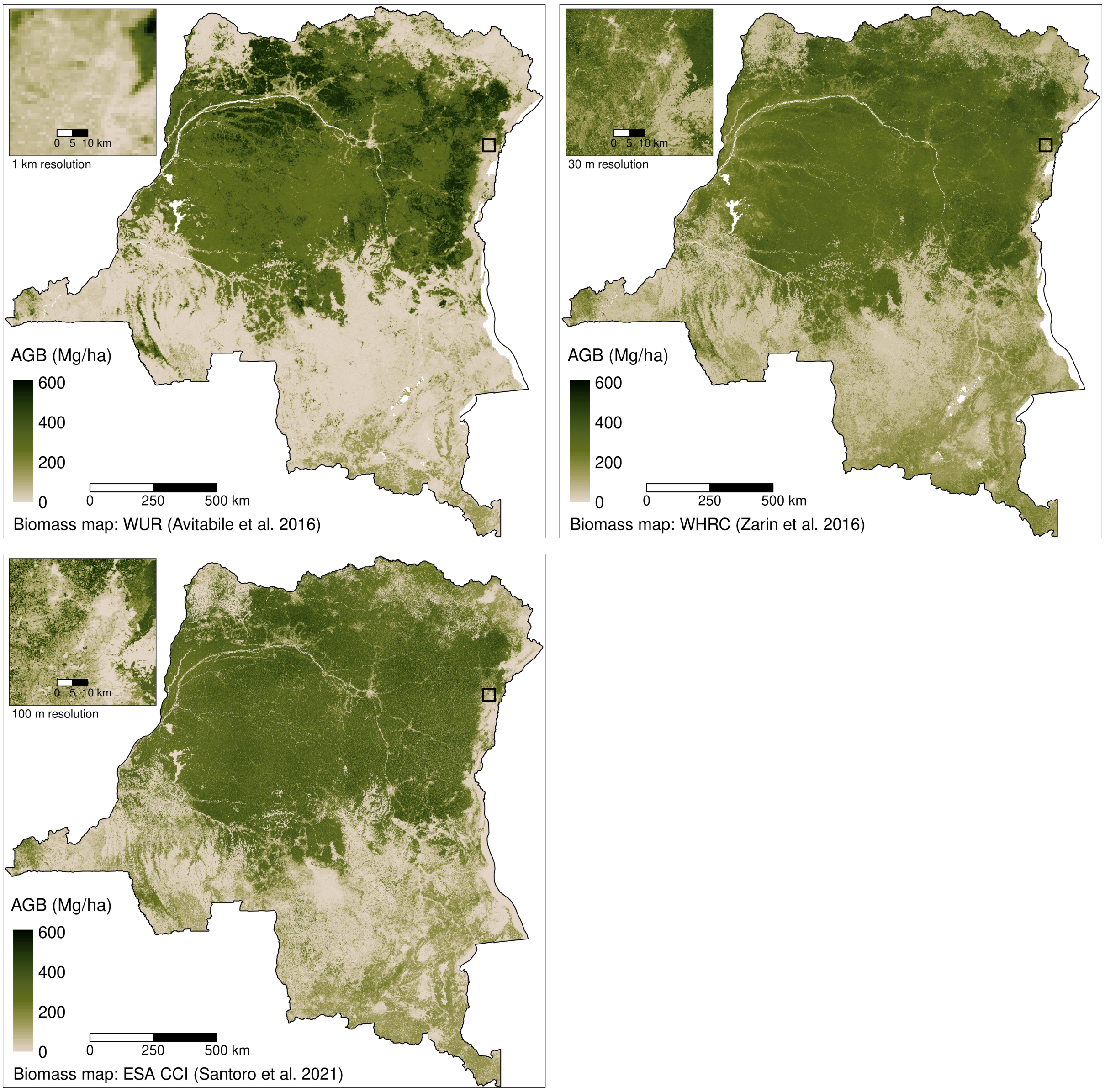

Figure S12: **Aboveground biomass maps**. This figure presents an extract of the three global or pantropical aboveground biomass (AGB in Mg/ha) maps which have been used for computing carbon emissions associated with deforestation: WUR map by Avitabile et al. (2016) for the years 2000–2010 (top left), WHRC map by Zarin et al. (2016) for the year 2000 (top right), and ESA CCI map by Santoro et al. (2021) for the year 2010 (bottom left). Extracts show biomass estimates for the Democratic Republic of the Congo (DRC). The three maps have different resolutions (1 km, 30 m, and 100 m, respectively). Insets represent zooms of the map for an area at the North-East of the country which is close to the city of Beni and the Virunga national park.

#### Fig. S13 – Change in annual carbon emissions

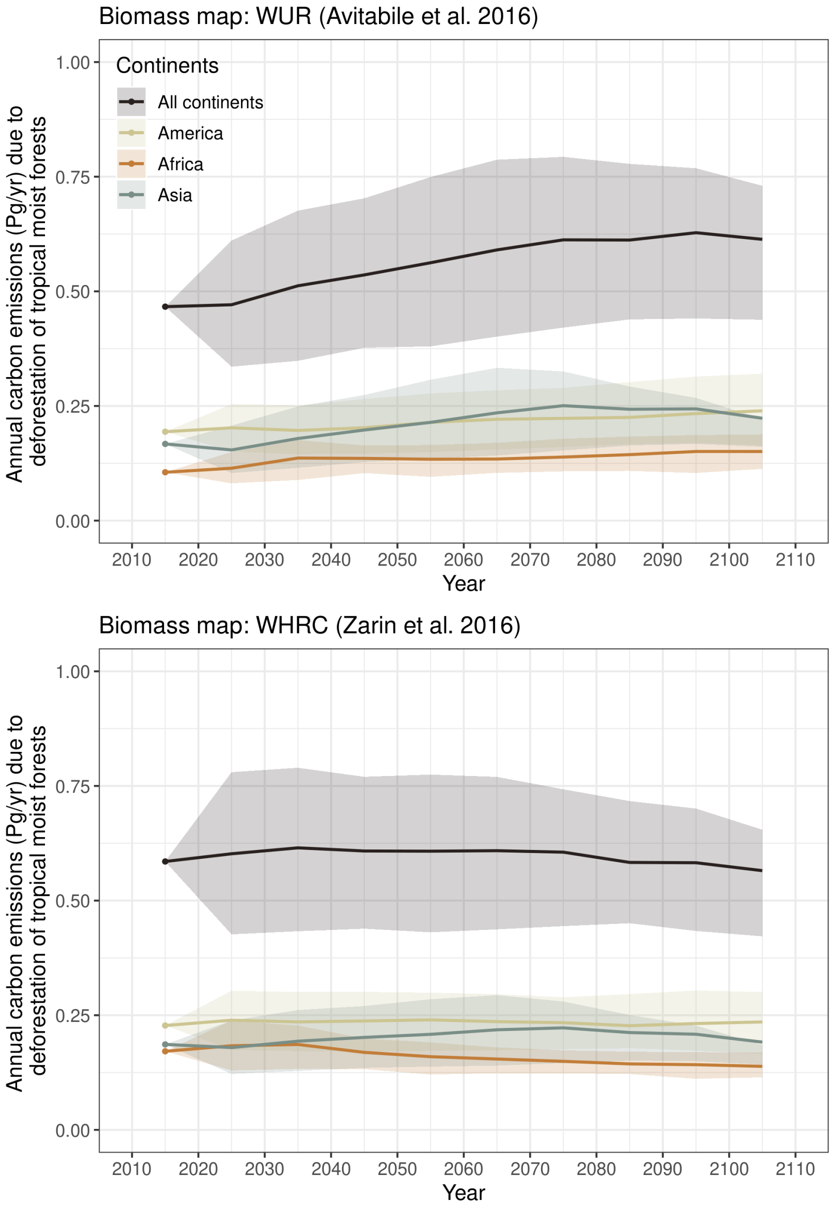

Figure S13: **Change in annual carbon emissions associated with projected deforestation**. Mean annual carbon emissions (Pg/yr) are computed for ten-year intervals from 2010–2020 to 2100–2110. The dots represent the observed mean annual carbon emissions (based on past deforestation maps) for the period 2010–2020, for the three continents (America, Africa, and Asia), and for the three continents combined. Lines represent the projected mean annual carbon emissions based on projected forest cover change maps from 2020–2030 to 2100–2110 per continent, and for all continents together. The confidence envelopes around the mean are obtained using the lower and upper bounds of the confidence intervals of the mean annual deforested areas for all study areas. *Top*: Carbon emissions obtained from the WUR aboveground biomass map by Avitabile et al. (2016) showing an increase of the annual carbon emissions from 0.467 Pg/yr in 2010–2020 to 0.628 Pg/yr (+35%) in 2090–2100 associated with the deforestation of forests with higher carbon stocks in the future. *Bottom*: Carbon emissions obtained from the WHRC aboveground biomass map by Zarin et al. (2016) showing constant annual carbon emissions of about 0.600 Pg/yr for the period 2010–2070, followed by a slight decrease of the emissions to 0.583 Pg/yr (-3%) in 2090–2100 associated with the complete loss of forest in some Asian countries.

#### Figs. S14–S15 – Uncertainty map of the future change in forest cover

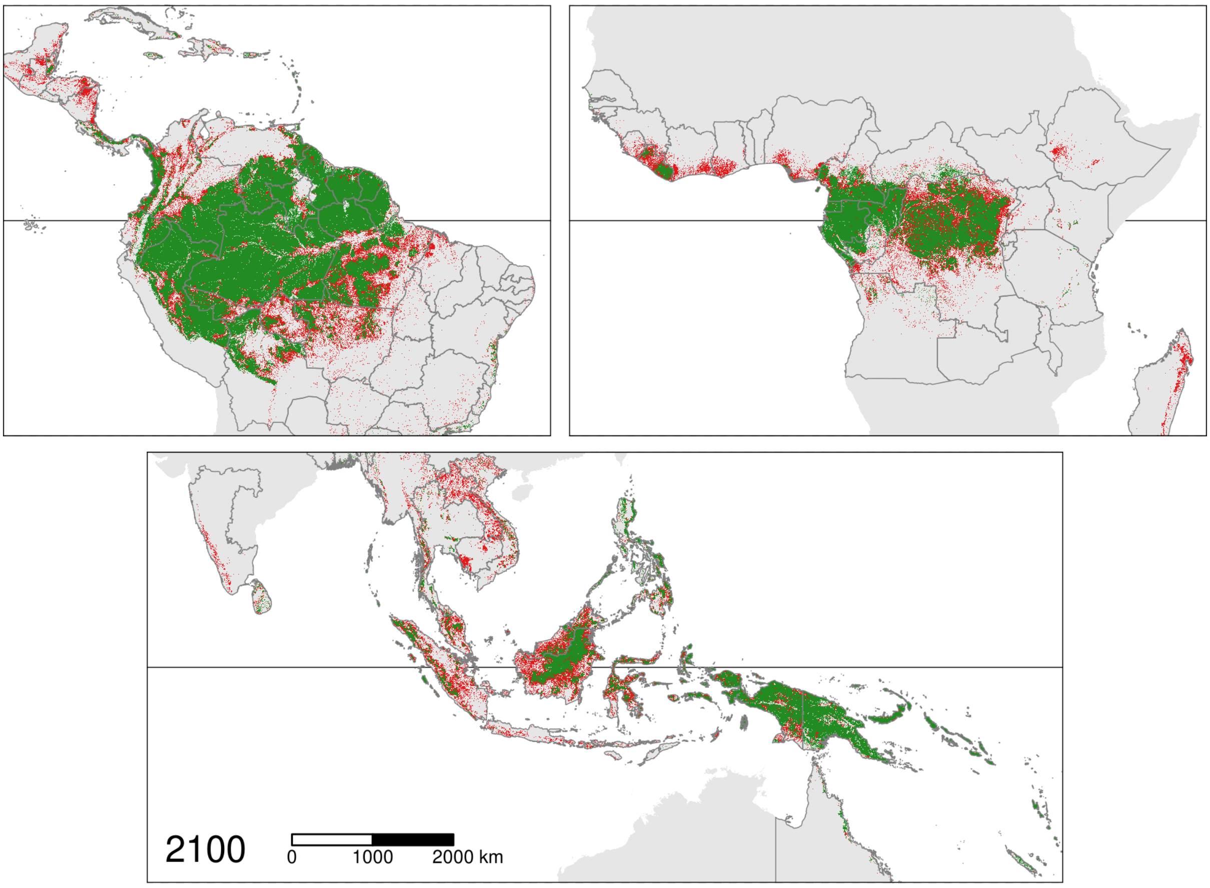

Figure S14: **Projected forest cover change map assuming a low annual deforestation**. This map was derived using $d'$, the *lower* bound of the confidence interval for the annual deforested area (in ha/yr) for each study area. This map must be compared with Fig. 1 in the main text and Fig. S15 below, which consider average and high annual deforested areas, respectively. The horizontal black line indicates the position of the Equator. The boundaries of the study areas are represented by dark grey lines. Forest areas in red are predicted to be deforested during the period 2020–2100, while forest areas in green are predicted to remain in 2100.

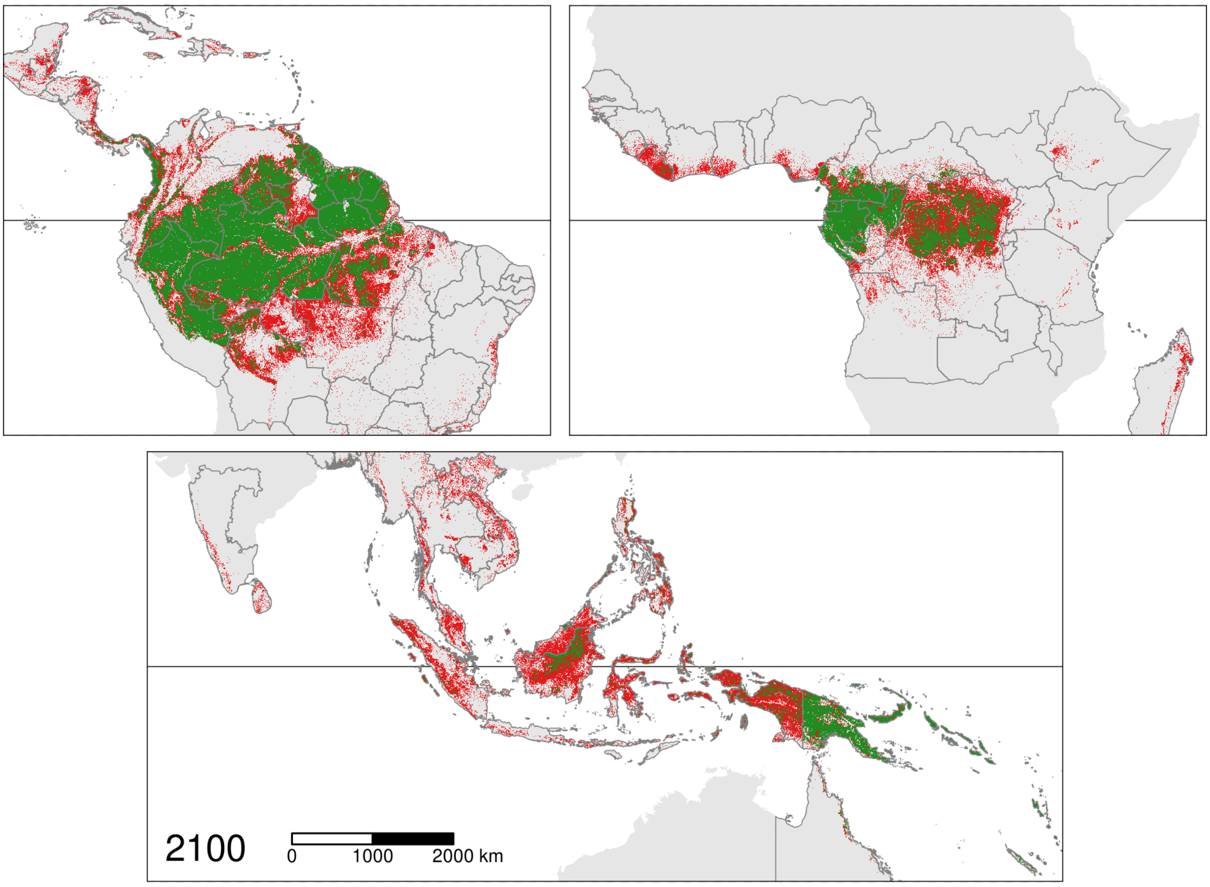

Figure S15: **Projected forest cover change map assuming a high annual deforestation**. This map was derived using $d''$, the *upper* bound of the confidence interval for the annual deforested area (in ha/yr) for each study area. This map must be compared with Fig. 1 in the main text and Fig. S14 above, which consider average and low annual deforested areas, respectively. The horizontal black line indicates the position of the Equator. The boundaries of the study areas are represented by dark grey lines. Forest areas in red are predicted to be deforested in the period 2020–2100, while forest areas in green are predicted to remain in 2100.

### 3 Supplementary tables

#### Table S1 – Pixel categories of the forest cover annual product

Table S1: **Pixel categories of the forest cover annual product by Vancutsem et al. (2021)**. The forest cover annual product v1_2020 classifies Landsat image pixels in 6 categories for each year (on the 1st of January) between 1990 and 2021 and allows identifying tropical moist forest pixels at each date.

| Class | Definition |
|---|---|
| 1 | Undisturbed Tropical Moist Forest (TMF) |
| 2 | Degraded TMF |
| 3 | Deforested land |
| 4 | Forest regrowth |
| 5 | Permanent or seasonal water |
| 6 | Other land cover |

#### Table S2 – Variables

Table S2: **Set of explanatory variables used to model the spatial probability of deforestation**. A total of height variables were tested. Variables describe topography, forest accessibility, forest landscape, deforestation history, and conservation status.

| Product | Source | Version / Date | Variable derived | Unit | Resolution (m) |
|---|---|---|---|---|---|
| Forest maps (2000–2010–2020) | Vancutsem et al. 2021 | v1_2020 | distance to forest edge | m | 30 |
| | | | distance to past deforestation | m | 30 |
| Digital Elevation Model | SRTM CSI-CGIAR | v4.1 | elevation | m | 90 |
| | | | slope | degree | 90 |
| Highways | OSM-Geofabrik | March 2021 | distance to road | m | 150 |
| Places | | | distance to town | m | 150 |
| Waterways | | | distance to river | m | 150 |
| Protected areas | WDPA | March 2021 | presence of protected area | – | 30 |

#### Table S3 – Sample size

Table S3: **Number of observations used for the spatial model of deforestation for each study area**. The table includes the number of non-deforested ("nfor") and deforested ("ndef") pixels per study area. These numbers include the forest pixels with full information regarding the explanatory variables. The corresponding number of hectares is also provided ("nfHa" and "ndHa", respectively).

| Country – Study area | Code | nfor | ndef | nfHa (ha) | ndHa (ha) |
|---|---|---|---|---|---|
| **America** | | | | | |
| Antigua and B. | ATG | 9,876 | 5,777 | 889 | 520 |
| Bahamas | BHS | 9,910 | 9,932 | 892 | 894 |
| Barbados | BRB | 9,964 | 7,004 | 897 | 630 |
| Belize | BLZ | 9,999 | 9,999 | 900 | 900 |
| Bolivia | BOL | 30,485 | 30,485 | 2,744 | 2,744 |
| Brazil – Acre | AC | 13,307 | 13,307 | 1,198 | 1,198 |
| Brazil – Alagoas | AL | 9,999 | 10,000 | 900 | 900 |
| Brazil – Amapa | AP | 11,563 | 11,564 | 1,041 | 1,041 |
| Brazil – Amazonas | AM | 50,000 | 50,000 | 4,500 | 4,500 |
| Brazil – Bahia | BA | 9,989 | 9,998 | 899 | 900 |
| Brazil – Ceara | CE | 9,996 | 9,999 | 900 | 900 |
| Brazil – Espirito Santo | ES | 9,998 | 10,000 | 900 | 900 |
| Brazil – Goias | GO | 10,000 | 10,000 | 900 | 900 |
| Brazil – Maranhao | MA | 9,985 | 9,998 | 899 | 900 |
| Brazil – Mato Grosso | MT | 33,283 | 33,283 | 2,995 | 2,995 |
| Brazil – Mato Grosso do Sul | MS | 10,000 | 10,000 | 900 | 900 |
| Brazil – Minas Gerais | MG | 10,000 | 10,000 | 900 | 900 |
| Brazil – Para | PA | 49,999 | 50,000 | 4,500 | 4,500 |
| Brazil – Paraiba | PB | 9,975 | 9,994 | 898 | 899 |
| Brazil – Parana | PR | 9,995 | 9,999 | 900 | 900 |
| Brazil – Pernambouco | PE | 9,975 | 9,997 | 898 | 900 |
| Brazil – Piaui | PI | 9,999 | 10,000 | 900 | 900 |
| Brazil – Rio de Janeiro | RJ | 9,992 | 9,980 | 899 | 898 |
| Brazil – Rio Grande do Norte | RN | 9,947 | 9,984 | 895 | 899 |
| Brazil – Rio Grande do Sul | RS | 10,000 | 9,999 | 900 | 900 |
| Brazil – Rondonia | RO | 13,800 | 13,800 | 1,242 | 1,242 |
| Brazil – Roraima | RR | 16,228 | 16,228 | 1,461 | 1,461 |
| Brazil – Santa Catarina | SC | 9,998 | 10,000 | 900 | 900 |
| Brazil – Sao Paulo | SP | 9,994 | 10,000 | 899 | 900 |
| Brazil – Sergipe | SE | 9,975 | 9,991 | 898 | 899 |
| Brazil – Tocantins | TO | 10,000 | 10,000 | 900 | 900 |
| Colombia | COL | 49,998 | 49,996 | 4,500 | 4,500 |
| Costa Rica | CRI | 9,992 | 9,987 | 899 | 899 |
| Cuba | CUB | 9,946 | 9,934 | 895 | 894 |
| Dominica | DMA | 9,991 | 9,870 | 899 | 888 |
| Dominican Rep. | DOM | 9,994 | 9,995 | 899 | 900 |
| Ecuador | ECU | 14,901 | 14,903 | 1,341 | 1,341 |
| El Salvador | SLV | 9,965 | 9,984 | 897 | 899 |
| French Guiana | GUF | 10,000 | 9,996 | 900 | 900 |
| Grenada | GRD | 9,984 | 9,923 | 899 | 893 |
| Guadeloupe | GLP | 9,972 | 9,926 | 897 | 893 |
| Guatemala | GTM | 9,999 | 9,999 | 900 | 900 |

Table S3: *(continued)*

| Country – Study area | Code | nfor | ndef | nfHa (ha) | ndHa (ha) |
|---|---|---|---|---|---|
| Guyana | GUY | 18,489 | 18,488 | 1,664 | 1,664 |
| Haiti | HTI | 9,959 | 9,969 | 896 | 897 |
| Honduras | HND | 9,998 | 10,000 | 900 | 900 |
| Jamaica | JAM | 9,998 | 9,987 | 900 | 899 |
| Martinique | MTQ | 9,985 | 9,968 | 899 | 897 |
| Mexico | MEX | 9,991 | 9,994 | 899 | 899 |
| Montserrat | MSR | 9,989 | 790 | 899 | 71 |
| Nicaragua | NIC | 9,998 | 10,000 | 900 | 900 |
| Panama | PAN | 9,994 | 9,992 | 899 | 899 |
| Paraguay | PRY | 10,000 | 10,000 | 900 | 900 |
| Peru | PER | 50,000 | 50,000 | 4,500 | 4,500 |
| Puerto Rico | PRI | 9,995 | 9,976 | 900 | 898 |
| Saint Kitts and N. | KNA | 9,978 | 3,800 | 898 | 342 |
| Saint Lucia | LCA | 9,995 | 9,968 | 900 | 897 |
| Saint Martin | MAF | 3,533 | 2,683 | 318 | 241 |
| Saint Vincent | VCT | 9,978 | 9,781 | 898 | 880 |
| Sint Maarten | SXM | 1,292 | 1,106 | 116 | 100 |
| Suriname | SUR | 13,727 | 13,726 | 1,235 | 1,235 |
| Trinidad and Tobago | TTO | 9,992 | 9,976 | 899 | 898 |
| Venezuela | VEN | 42,913 | 42,910 | 3,862 | 3,862 |
| Virgin Isl. UK | VGB | 9,905 | 9,776 | 891 | 880 |
| Virgin Isl. US | VIR | 9,914 | 9,834 | 892 | 885 |
| **Africa** | | | | | |
| Angola | AGO | 9,999 | 10,000 | 900 | 900 |
| Benin | BEN | 9,978 | 9,997 | 898 | 900 |
| Burundi | BDI | 10,000 | 10,000 | 900 | 900 |
| Cameroon | CMR | 23,543 | 23,541 | 2,119 | 2,119 |
| CAR | CAF | 10,000 | 10,000 | 900 | 900 |
| Comoros | COM | 9,993 | 9,962 | 899 | 897 |
| Congo | COG | 23,945 | 23,943 | 2,155 | 2,155 |
| DRC | COD | 50,000 | 50,000 | 4,500 | 4,500 |
| Eq. Guinea | GNQ | 9,998 | 9,985 | 900 | 899 |
| Ethiopia | ETH | 10,000 | 10,000 | 900 | 900 |
| Gabon | GAB | 24,096 | 24,077 | 2,169 | 2,167 |
| Gambia | GMB | 9,979 | 9,996 | 898 | 900 |
| Ghana | GHA | 10,000 | 10,000 | 900 | 900 |
| Guinea | GIN | 9,976 | 9,999 | 898 | 900 |
| Guinea Bissau | GNB | 9,874 | 9,969 | 889 | 897 |
| Ivory Coast | CIV | 10,000 | 9,999 | 900 | 900 |
| Kenya | KEN | 9,989 | 9,998 | 899 | 900 |
| Liberia | LBR | 10,000 | 9,997 | 900 | 900 |
| Madagascar | MDG | 9,991 | 9,996 | 899 | 900 |
| Malawi | MWI | 10,000 | 10,000 | 900 | 900 |
| Mauritius | MUS | 9,972 | 9,952 | 897 | 896 |
| Mayotte | MYT | 9,975 | 9,973 | 898 | 898 |
| Nigeria | NGA | 9,979 | 9,997 | 898 | 900 |
| Reunion | REU | 9,996 | 9,983 | 900 | 898 |
| Rwanda | RWA | 10,000 | 10,000 | 900 | 900 |
| Senegal | SEN | 9,883 | 9,972 | 889 | 897 |

Table S3: *(continued)*

| Country – Study area | Code | nfor | ndef | nfHa (ha) | ndHa (ha) |
|---|---|---|---|---|---|
| Sierra Leone | SLE | 9,993 | 9,999 | 899 | 900 |
| South Sudan | SSD | 5,682 | 7,705 | 511 | 693 |
| Tanzania | TZA | 9,978 | 9,973 | 898 | 898 |
| Togo | TGO | 10,000 | 10,000 | 900 | 900 |
| Uganda | UGA | 10,000 | 10,000 | 900 | 900 |
| Zambia | ZMB | 10,000 | 10,000 | 900 | 900 |
| **Asia** | | | | | |
| Australia – Queensland | QLD | 9,978 | 9,972 | 898 | 897 |
| Bangladesh | BGD | 9,956 | 9,972 | 896 | 897 |
| Bhutan | BTN | 10,000 | 10,000 | 900 | 900 |
| Brunei | BRN | 9,990 | 9,996 | 899 | 900 |
| Cambodia | KHM | 9,995 | 9,999 | 900 | 900 |
| Fiji | FJI | 9,984 | 9,954 | 899 | 896 |
| India – Andaman and N. | AN | 9,954 | 9,900 | 896 | 891 |
| India – North-East | NE | 9,996 | 9,999 | 900 | 900 |
| India – West. Ghats | WG | 9,996 | 9,993 | 900 | 899 |
| Indonesia | IDN | 49,976 | 49,975 | 4,498 | 4,498 |
| Laos | LAO | 10,000 | 10,000 | 900 | 900 |
| Malaysia | MYS | 22,307 | 22,310 | 2,008 | 2,008 |
| Myanmar | MMR | 15,371 | 15,376 | 1,383 | 1,384 |
| New Caledonia | NCL | 9,965 | 9,903 | 897 | 891 |
| Papua New Guinea | PNG | 39,764 | 39,670 | 3,579 | 3,570 |
| Philippines | PHL | 13,665 | 13,662 | 1,230 | 1,230 |
| Singapore | SGP | 9,902 | 9,947 | 891 | 895 |
| Solomon Isl. | SLB | 9,961 | 9,773 | 896 | 880 |
| Sri Lanka | LKA | 9,996 | 9,995 | 900 | 900 |
| Thailand | THA | 9,991 | 9,995 | 899 | 900 |
| Timor-Leste | TLS | 9,993 | 9,972 | 899 | 897 |
| Vanuatu | VUT | 9,974 | 9,890 | 898 | 890 |
| Vietnam | VNM | 9,996 | 9,996 | 900 | 900 |
| **All continents** | | | | | |
| TOTAL | | 1,610,125 | 1,587,817 | 144,914 | 142,908 |

#### Tables S4–S5 – Parameter estimates and variable importance

Table S4: **Parameter estimates for each study area**. For each study area, we computed the posterior mean of each parameter (“int”: intercept, “pa”: protected area effect, “elev”, “slope”, “ddefor”,“dedge”, “driver”, “droad”, “dtown”: slope parameters associated to elevation, slope, distance to past deforestation, distance to forest edge, distance to nearest river, distance to nearest road, and distance to nearest town, respectively, “Vrho”: variance of the spatial random effects). Continuous explanatory variables were normalized (mean=0 and standard-deviation=1), allowing us to estimate the relative importance of each variable in determining the spatial probability of deforestation from parameter values. Comparison can be done within and between study areas.

| study area | int | pa | elev | slope | ddefor | dedge | driver | droad | dtown | Vrho |
|---|---|---|---|---|---|---|---|---|---|---|
| **America** | | | | | | | | | | |
| ATG | 0.110 | -0.710 | -0.393 | – | -0.719 | -1.170 | – | – | – | 10.00 |
| BHS | -0.307 | – | -0.113 | -0.091 | -1.020 | -0.525 | -0.116 | – | – | 8.20 |
| BRB | -0.414 | -0.323 | -0.442 | -0.201 | -0.248 | -0.958 | -0.009 | 0.000 | – | 1.99 |
| BLZ | -0.905 | -0.653 | -0.058 | -0.194 | -1.270 | -1.280 | – | -0.129 | -0.383 | 6.97 |
| BOL | -0.697 | -0.163 | -0.292 | -0.223 | -1.050 | -3.070 | – | -0.306 | -0.102 | 5.36 |
| AC | -6.720 | -0.631 | – | – | -11.100 | -4.450 | – | -0.179 | -0.162 | 4.14 |
| AL | 0.735 | -0.192 | -0.333 | -0.055 | -0.632 | -1.690 | -0.139 | -0.214 | -0.140 | 3.01 |
| AP | -6.110 | -0.126 | -0.748 | -0.064 | -4.100 | -8.460 | – | -0.218 | -0.273 | 5.22 |
| AM | -3.950 | -0.806 | -0.118 | – | -2.800 | -4.330 | -0.157 | -0.898 | -0.416 | 13.70 |
| BA | 1.370 | -0.213 | -0.268 | -0.201 | -1.010 | -0.969 | – | -0.032 | – | 4.11 |
| CE | 1.200 | -0.551 | -1.150 | – | -0.214 | -0.956 | – | – | – | 8.49 |
| ES | -0.335 | -0.193 | -0.813 | – | -0.852 | -1.210 | -0.068 | 0.002 | -0.078 | 3.20 |
| GO | 0.043 | – | -0.047 | – | -0.790 | -0.447 | -0.116 | – | -0.025 | 3.43 |
| MA | -1.440 | -0.202 | – | -0.086 | -4.670 | -2.730 | – | – | -0.172 | 3.82 |
| MT | -0.506 | -0.427 | – | – | -1.530 | -0.647 | – | -0.136 | -0.240 | 10.20 |
| MS | -0.361 | – | -0.039 | -0.009 | -0.427 | -1.210 | – | – | – | 4.07 |
| MG | 0.058 | -0.153 | -0.504 | – | -0.738 | -0.515 | -0.059 | -0.058 | -0.038 | 3.86 |
| PA | -1.230 | -1.180 | – | -0.033 | -1.400 | -2.330 | -0.019 | -0.874 | -0.306 | 8.08 |
| PB | 3.030 | -0.693 | – | -0.043 | -0.821 | -1.070 | -0.244 | – | -0.094 | 4.35 |
| PR | -0.739 | -0.339 | – | -0.218 | -0.959 | -1.980 | – | – | -0.037 | 2.54 |
| PE | 7.390 | -0.558 | -0.730 | -0.173 | -0.861 | -2.100 | -0.033 | -0.066 | -0.082 | 6.55 |
| PI | 0.377 | – | – | – | -0.624 | -0.626 | -0.025 | -0.019 | -0.088 | 4.13 |
| RJ | -0.671 | -0.220 | -0.080 | -0.009 | -1.490 | -3.140 | -0.142 | -0.087 | -0.036 | 2.40 |
| RN | -0.087 | – | – | -0.084 | -0.938 | -0.625 | – | – | – | 5.73 |
| RS | -0.171 | -0.343 | -0.107 | -0.504 | -0.622 | -1.660 | -0.002 | -0.002 | – | 1.81 |
| RO | -0.719 | -1.630 | – | -0.108 | -0.598 | -1.790 | -0.003 | -0.403 | -0.141 | 7.27 |
| RR | -1.790 | -0.841 | -0.446 | – | -2.720 | -2.050 | -0.012 | -0.165 | -0.427 | 8.49 |
| SC | -0.675 | -0.303 | – | -0.548 | -0.921 | -1.560 | – | – | -0.052 | 2.00 |
| SP | -0.686 | -0.184 | -0.166 | -0.260 | -1.870 | -2.800 | -0.011 | -0.024 | -0.042 | 3.79 |
| SE | 0.669 | – | – | – | -0.918 | -0.866 | -0.132 | – | – | 3.63 |
| TO | -0.351 | -0.035 | – | – | -0.590 | -0.032 | -0.081 | -0.046 | – | 4.43 |
| COL | -2.410 | -0.449 | -0.803 | -0.330 | -4.060 | -2.050 | – | -0.591 | -0.349 | 5.62 |
| CRI | -3.510 | – | -0.065 | -0.303 | -3.050 | -6.700 | -0.009 | -0.064 | -0.136 | 2.18 |
| CUB | -0.371 | – | -0.242 | -0.105 | -0.694 | -0.707 | -0.053 | -0.126 | -0.053 | 4.77 |
| DMA | -1.470 | – | -0.216 | -0.181 | -0.840 | -2.700 | – | – | -0.407 | 3.26 |
| DOM | -0.138 | -0.331 | -0.236 | -0.075 | -1.080 | -1.390 | -0.003 | -0.130 | -0.005 | 2.85 |
| ECU | -4.340 | -0.520 | -0.097 | -0.362 | -2.790 | -10.600 | – | -0.433 | -0.107 | 2.53 |
| SLV | -0.048 | -0.476 | -0.327 | -0.213 | -0.820 | -1.590 | – | -0.073 | -0.033 | 3.38 |
| GUF | -1.220 | -0.835 | -0.584 | -0.122 | -1.200 | -2.690 | – | -1.050 | -0.880 | 18.80 |
| GRD | -0.776 | -0.396 | -1.120 | – | -1.840 | -2.700 | – | -0.266 | -0.108 | 14.20 |
| GLP | -3.960 | – | -0.171 | -0.190 | -2.020 | -7.360 | -0.036 | -0.089 | -0.090 | 2.61 |
| GTM | 0.026 | -0.229 | -0.427 | -0.252 | -1.030 | -0.145 | -0.022 | -0.180 | -0.036 | 3.15 |
| GUY | -1.970 | -0.708 | -0.452 | -0.207 | -0.479 | -4.860 | – | -0.890 | -0.330 | 13.70 |

Table S4: *(continued)*

| study area | int | pa | elev | slope | ddefor | dedge | driver | droad | dtown | Vrho |
|---|---|---|---|---|---|---|---|---|---|---|
| HTI | 0.034 | -0.038 | – | -0.120 | -0.587 | -0.751 | -0.078 | -0.256 | -0.015 | 2.87 |
| HND | -0.976 | -0.364 | -0.911 | -0.307 | -0.939 | -0.590 | -0.017 | – | – | 4.46 |
| JAM | -0.817 | -0.003 | -0.242 | -0.132 | -2.160 | -2.640 | -0.024 | -0.136 | -0.078 | 2.66 |
| MTQ | -1.410 | -0.075 | -0.627 | -0.053 | -0.900 | -4.390 | – | – | – | 2.66 |
| MEX | -0.769 | -0.228 | -0.317 | -0.277 | -1.080 | -1.150 | – | -0.056 | -0.029 | 3.59 |
| MSR | -8.160 | – | -0.213 | – | -1.740 | -6.020 | – | – | – | 4.81 |
| NIC | -1.360 | – | -0.327 | -0.088 | -0.612 | -1.680 | – | -0.158 | – | 3.19 |
| PAN | -2.510 | -0.455 | -0.157 | -0.239 | -4.100 | -2.790 | – | -0.511 | – | 2.97 |
| PRY | -0.404 | – | -0.066 | -0.109 | -0.785 | -0.126 | -0.007 | -0.096 | – | 3.93 |
| PER | -1.730 | -0.617 | -0.634 | -0.354 | -3.580 | -3.610 | -0.022 | -0.570 | -0.504 | 6.44 |
| PRI | 0.009 | – | -0.383 | -0.062 | -0.410 | -1.100 | – | – | -0.064 | 4.32 |
| KNA | -1.960 | – | -1.130 | -0.168 | -0.297 | -0.878 | – | – | – | 1.62 |
| LCA | -1.350 | – | -0.265 | -0.066 | -1.140 | -3.140 | – | -0.035 | -0.061 | 2.97 |
| MAF | 0.438 | – | – | – | -0.391 | -0.895 | -0.477 | – | – | 8.11 |
| VCT | -0.526 | -0.159 | -0.691 | -0.164 | -1.950 | -1.840 | – | -0.303 | -0.045 | 3.30 |
| SXM | -0.029 | – | – | -0.491 | -0.597 | -0.527 | -0.068 | – | -0.298 | 10.00 |
| SUR | -1.170 | -0.046 | -0.548 | -0.144 | -0.551 | -1.210 | – | -1.210 | -0.392 | 12.50 |
| TTO | -2.550 | -0.265 | -0.249 | -0.153 | -2.290 | -5.430 | -0.038 | -0.106 | -0.105 | 2.16 |
| VEN | -2.930 | -0.120 | -0.390 | -0.089 | -1.390 | -6.690 | -0.080 | -0.585 | -0.360 | 6.59 |
| VGB | 1.070 | – | -0.347 | -0.210 | -0.939 | -0.973 | – | – | – | 8.04 |
| VIR | 0.030 | -0.350 | – | -0.174 | -0.311 | -0.782 | – | – | – | 7.64 |
| **Africa** | | | | | | | | | | |
| AGO | -0.382 | – | -0.042 | -0.096 | -2.720 | -2.040 | -0.013 | -0.194 | -0.066 | 4.00 |
| BEN | 2.760 | – | – | -0.148 | -0.422 | -1.000 | – | – | – | 8.05 |
| BDI | -0.691 | -1.170 | -0.527 | – | -2.130 | -3.320 | -0.090 | – | -0.062 | 6.39 |
| CMR | -0.130 | -0.891 | – | -0.156 | -1.570 | -2.990 | -0.034 | -0.419 | -0.270 | 5.96 |
| CAF | -0.226 | -0.094 | – | -0.022 | -4.950 | -1.510 | -0.109 | -0.110 | -0.370 | 4.61 |
| COM | -2.730 | – | -0.163 | -0.314 | – | -10.100 | – | – | – | 7.22 |
| COG | -2.350 | -0.402 | -0.012 | -0.064 | -1.420 | -5.410 | -0.160 | -0.548 | -0.314 | 8.71 |
| COD | -3.910 | -0.160 | – | – | -6.450 | -4.590 | – | -0.375 | -0.295 | 4.52 |
| GNQ | -1.050 | -0.136 | -0.364 | -0.340 | -0.031 | -2.660 | -0.072 | -1.300 | -0.091 | 8.70 |
| ETH | -0.589 | -0.082 | -0.066 | -0.270 | -1.120 | -1.960 | – | -0.082 | -0.100 | 3.55 |
| GAB | -2.570 | -0.160 | -0.515 | -0.222 | -1.140 | -4.200 | – | -0.541 | -0.433 | 14.60 |
| GMB | -0.014 | -0.222 | – | -0.106 | -0.543 | -0.399 | – | -0.327 | -0.081 | 5.23 |
| GHA | 0.470 | -0.378 | -0.125 | -0.182 | -0.551 | -1.740 | – | -0.015 | -0.057 | 2.06 |
| GIN | -0.676 | -0.224 | -0.078 | -0.013 | -3.220 | -2.510 | – | -0.084 | – | 1.88 |
| GNB | 0.652 | -0.688 | – | – | -1.270 | -0.866 | – | -0.125 | -0.057 | 3.13 |
| CIV | 0.006 | – | – | -0.133 | -1.680 | -0.828 | – | -0.020 | -0.041 | 2.61 |
| KEN | -1.540 | -0.186 | -0.525 | -0.066 | -0.764 | -2.990 | -0.062 | -0.158 | -0.044 | 6.97 |
| LBR | -1.540 | -0.491 | – | -0.129 | -1.310 | -3.150 | – | -0.331 | -0.302 | 2.06 |
| MDG | -1.170 | -0.285 | -0.531 | -0.169 | -2.260 | -1.260 | – | – | -0.096 | 3.43 |
| MWI | -0.089 | -0.363 | -0.122 | -0.148 | -0.976 | -0.202 | -0.246 | -0.515 | 0.000 | 18.70 |
| MUS | -1.890 | -0.450 | -0.153 | -0.191 | -0.538 | -3.630 | -0.017 | -0.091 | -0.031 | 1.37 |
| MYT | -0.230 | -1.340 | – | – | -0.751 | -1.360 | -0.058 | – | -0.034 | 2.77 |
| NGA | 1.020 | – | – | -0.015 | -1.330 | -0.776 | – | -0.220 | – | 5.63 |
| REU | -0.932 | -0.385 | – | -0.206 | -0.184 | -4.200 | – | – | – | 2.76 |
| RWA | -2.260 | -1.300 | -0.337 | -0.215 | -1.560 | -5.180 | – | -0.138 | – | 3.98 |
| SEN | 1.690 | -0.199 | -0.012 | -0.065 | -0.794 | -0.401 | – | – | -0.062 | 6.03 |
| SLE | -0.374 | -0.247 | – | – | -1.360 | -1.190 | – | -0.038 | -0.052 | 1.31 |
| SSD | 0.850 | -0.135 | -0.182 | 0.005 | -0.266 | -0.644 | – | -0.203 | – | 4.16 |
| TZA | -0.268 | -0.239 | -0.434 | -0.090 | -1.040 | -3.160 | -0.109 | -0.184 | -0.067 | 5.93 |
| TGO | 1.960 | -0.537 | -0.190 | -0.144 | -0.365 | -0.525 | – | -0.271 | – | 4.68 |
| UGA | -1.340 | -1.190 | – | – | -2.700 | -2.480 | -0.070 | -0.036 | -0.116 | 4.72 |
| ZMB | -0.325 | -0.401 | – | – | -1.020 | -0.439 | – | -0.125 | -0.078 | 9.53 |

Table S4: *(continued)*

| study area | int | pa | elev | slope | ddefor | dedge | driver | droad | dtown | Vrho |
|---|---|---|---|---|---|---|---|---|---|---|
| **Asia** | | | | | | | | | | |
| QLD | -0.110 | -0.349 | – | -0.122 | -1.190 | -1.660 | – | 0.009 | – | 5.56 |
| BGD | 0.043 | -0.544 | -0.081 | -0.092 | -0.625 | -1.070 | -0.060 | – | – | 4.47 |
| BTN | -1.310 | – | -0.072 | -0.077 | -7.070 | -1.480 | -0.046 | – | -0.049 | 1.19 |
| BRN | -0.441 | -1.100 | -0.984 | -0.370 | -2.430 | – | – | -0.087 | -0.617 | 15.10 |
| KHM | 0.572 | -1.460 | -1.040 | -0.193 | -0.998 | -0.286 | – | -0.387 | -0.182 | 9.40 |
| FJI | -1.520 | -0.306 | -0.254 | -0.237 | -0.416 | -4.950 | – | -0.124 | -0.088 | 5.21 |
| AN | -5.620 | – | -0.223 | -0.193 | -1.120 | -17.300 | – | -0.242 | – | 4.55 |
| NE | -1.040 | -0.585 | -0.309 | -0.293 | -5.390 | -1.600 | -0.029 | -0.094 | -0.033 | 2.39 |
| WG | -0.156 | -0.369 | -0.483 | -0.084 | -0.692 | -2.060 | – | -0.170 | -0.053 | 1.93 |
| IDN | -1.330 | -0.736 | -0.312 | -0.583 | -1.660 | -1.810 | -0.037 | -0.433 | -0.088 | 8.13 |
| LAO | -0.258 | -0.442 | -0.405 | -0.340 | -1.230 | -0.941 | – | -0.162 | -0.174 | 3.08 |
| MYS | -0.687 | -2.080 | -0.385 | -0.472 | -0.830 | -1.890 | – | -0.241 | – | 7.84 |
| MMR | -0.431 | -0.167 | -0.483 | -0.215 | -1.370 | -1.420 | -0.107 | -0.139 | -0.018 | 3.10 |
| NCL | -2.000 | – | -0.368 | -0.098 | -2.340 | -8.160 | – | 0.020 | -0.077 | 4.11 |
| PNG | -1.630 | – | -0.549 | -0.478 | -0.072 | -3.840 | -0.055 | -0.358 | -0.116 | 7.46 |
| PHL | -1.570 | -0.153 | -0.226 | -0.393 | -2.010 | -2.420 | – | -0.093 | -0.182 | 2.97 |
| SGP | -0.013 | -1.350 | -0.119 | -0.108 | -1.170 | -1.330 | – | -0.616 | – | 6.01 |
| SLB | -0.544 | – | -0.802 | -0.513 | – | -0.866 | -0.221 | -0.357 | – | 5.86 |
| LKA | 0.034 | -0.516 | -0.090 | -0.291 | -1.030 | -1.480 | -0.035 | -0.068 | -0.196 | 1.71 |
| THA | -1.390 | -0.159 | -0.751 | -0.080 | -1.930 | -3.370 | -0.012 | – | – | 2.07 |
| TLS | -0.563 | -0.120 | -0.189 | -0.104 | -1.330 | -1.490 | -0.034 | -0.080 | -0.124 | 1.60 |
| VUT | -3.220 | -0.505 | -0.575 | -0.287 | -0.859 | -5.210 | – | – | 0.004 | 18.10 |
| VNM | -0.612 | -0.536 | -0.283 | -0.376 | -1.650 | -1.220 | -0.045 | -0.173 | -0.059 | 3.12 |

Table S5: **Parameter estimate weighted means per region**. We used the forest cover in 2010 to compute the parameter estimate weighted mean per region. Continuous explanatory variables were normalized (mean=0 and standard-deviation=1), allowing us to estimate the relative importance of each variable in determining the spatial probability of deforestation from parameter values. Comparison can be done within and between regions. At pantropical scale (when considering all continents together), explanatory variables can be classified in the following decreasing order of importance: distance to forest edge, distance to past deforestation, presence of a protected area, distance to nearest road, elevation, distance to nearest town, slope, and distance to nearest river.

| Region | int | pa | elev | slope | ddefor | dedge | driver | droad | dtown | Vrho |
|---|---|---|---|---|---|---|---|---|---|---|
| India | -1.074 | -0.484 | -0.354 | -0.225 | -3.742 | -2.739 | -0.019 | -0.126 | -0.037 | 2.393 |
| Brazil | -2.633 | -0.837 | -0.103 | -0.028 | -2.508 | -3.228 | -0.073 | -0.661 | -0.323 | 10.093 |
| America | -2.342 | -0.636 | -0.297 | -0.134 | -2.490 | -3.468 | -0.048 | -0.615 | -0.328 | 8.549 |
| Africa | -2.463 | -0.259 | -0.073 | -0.066 | -4.002 | -3.823 | -0.024 | -0.365 | -0.270 | 5.819 |
| Asia | -1.164 | -0.625 | -0.385 | -0.467 | -1.415 | -2.149 | -0.037 | -0.316 | -0.084 | 6.676 |
| All continents | -2.099 | -0.551 | -0.268 | -0.195 | -2.577 | -3.245 | -0.040 | -0.492 | -0.260 | 7.519 |

#### Table S6 – Effect of protected areas on deforestation

Table S6: **Effect of protected areas on deforestation**. We show here the estimated effect of the presence of a protected area on the probability of deforestation for each study area. We computed the mean (“Mean”), the standard-deviation (“Sd”), and the bayesian 95% credible interval (“CI 95%”) of the estimated parameter. Column “P” indicates the percentage of decrease in the deforestation risk inside protected areas. Column “signif” indicates (with a star) that the estimated effect was negative and significantly different from zero (zero not included in the credible interval). Out of the 119 study areas, 70 showed a significant negative effect (59% of the countries). These 70 study areas accounted for 88% of the moist tropical forest in 2010 (“fc2010” in Kha).

| Country – study area | fc2010 (Kha) | Mean | Sd | CI 95% | P % | signif |
|---|---|---|---|---|---|---|
| **America** | | | | | | |
| Antigua and B. | 4 | -0.710 | 0.232 | (-1.180, -0.257) | 33 | ⋆ |
| Bahamas | 115 | – | – | – | – | |
| Barbados | 4 | -0.323 | 0.172 | (-0.667, 0.033) | 19 | |
| Belize | 1,328 | -0.653 | 0.116 | (-0.876, -0.425) | 40 | ⋆ |
| Bolivia | 30,485 | -0.163 | 0.065 | (-0.296, -0.033) | 11 | ⋆ |
| Brazil – Acre | 13,307 | -0.631 | 0.099 | (-0.834, -0.432) | 47 | ⋆ |
| Brazil – Alagoas | 98 | -0.192 | 0.103 | (-0.401, 0.008) | 6 | |
| Brazil – Amapa | 11,565 | -0.126 | 0.105 | (-0.347, 0.073) | 12 | |
| Brazil – Amazonas | 146,852 | -0.806 | 0.060 | (-0.925, -0.692) | 55 | ⋆ |
| Brazil – Bahia | 2,097 | -0.213 | 0.075 | (-0.360, -0.072) | 5 | ⋆ |
| Brazil – Ceara | 48 | -0.551 | 0.117 | (-0.785, -0.334) | 15 | ⋆ |
| Brazil – Espirito Santo | 417 | -0.193 | 0.083 | (-0.360, -0.031) | 11 | ⋆ |
| Brazil – Goias | 481 | – | – | – | – | |
| Brazil – Maranhao | 3,930 | -0.202 | 0.096 | (-0.390, -0.024) | 15 | ⋆ |
| Brazil – Mato Grosso | 33,283 | -0.427 | 0.070 | (-0.560, -0.295) | 25 | ⋆ |
| Brazil – Mato Grosso do Sul | 736 | – | – | – | – | |
| Brazil – Minas Gerais | 1,277 | -0.153 | 0.087 | (-0.318, 0.033) | 7 | |
| Brazil – Para | 91,982 | -1.180 | 0.059 | (-1.280, -1.050) | 64 | ⋆ |
| Brazil – Paraiba | 41 | -0.693 | 0.113 | (-0.936, -0.477) | 4 | ⋆ |
| Brazil – Parana | 2,673 | -0.339 | 0.082 | (-0.506, -0.186) | 21 | ⋆ |
| Brazil – Pernambouco | 119 | -0.558 | 0.093 | (-0.733, -0.378) | 0 | ⋆ |
| Brazil – Piaui | 74 | – | – | – | – | |
| Brazil – Rio de Janeiro | 736 | -0.220 | 0.068 | (-0.369, -0.093) | 14 | ⋆ |
| Brazil – Rio Grande do Norte | 25 | – | – | – | – | |
| Brazil – Rio Grande do Sul | 2,214 | -0.343 | 0.133 | (-0.588, -0.068) | 18 | ⋆ |
| Brazil – Rondonia | 13,800 | -1.630 | 0.076 | (-1.780, -1.480) | 73 | ⋆ |
| Brazil – Roraima | 16,228 | -0.841 | 0.115 | (-1.090, -0.633) | 53 | ⋆ |
| Brazil – Santa Catarina | 2,485 | -0.303 | 0.104 | (-0.505, -0.096) | 19 | ⋆ |
| Brazil – Sao Paulo | 2,781 | -0.184 | 0.071 | (-0.312, -0.040) | 12 | ⋆ |
| Brazil – Sergipe | 61 | – | – | – | – | |
| Brazil – Tocantins | 1,341 | -0.035 | 0.090 | (-0.193, 0.152) | 2 | |
| Colombia | 66,802 | -0.449 | 0.048 | (-0.542, -0.357) | 34 | ⋆ |
| Costa Rica | 2,277 | – | – | – | – | |
| Cuba | 1,297 | – | – | – | – | |
| Dominica | 70 | – | – | – | – | |
| Dominican Rep. | 993 | -0.331 | 0.072 | (-0.482, -0.195) | 17 | ⋆ |
| Ecuador | 14,903 | -0.520 | 0.092 | (-0.696, -0.342) | 40 | ⋆ |
| El Salvador | 109 | -0.476 | 0.107 | (-0.698, -0.286) | 24 | ⋆ |
| French Guiana | 8,088 | -0.835 | 0.162 | (-1.150, -0.524) | 50 | ⋆ |

Table S6: *(continued)*

| Country – study area | fc2010 (Kha) | Mean | Sd | CI 95% | P % | signif |
|---|---|---|---|---|---|---|
| Grenada | 22 | -0.396 | 0.323 | (-1.020, 0.245) | 25 | |
| Guadeloupe | 77 | – | – | – | – | |
| Guatemala | 2,702 | -0.229 | 0.089 | (-0.401, -0.051) | 11 | ⋆ |
| Guyana | 18,489 | -0.708 | 0.234 | (-1.190, -0.285) | 47 | ⋆ |
| Haiti | 162 | -0.038 | 0.086 | (-0.200, 0.138) | 2 | |
| Honduras | 2,993 | -0.364 | 0.078 | (-0.527, -0.215) | 24 | ⋆ |
| Jamaica | 421 | -0.003 | 0.098 | (-0.189, 0.202) | 0 | |
| Martinique | 70 | -0.075 | 0.155 | (-0.386, 0.209) | 6 | |
| Mexico | 7,390 | -0.228 | 0.068 | (-0.355, -0.087) | 15 | ⋆ |
| Montserrat | 3 | – | – | – | – | |
| Nicaragua | 4,262 | – | – | – | – | |
| Panama | 4,204 | -0.455 | 0.079 | (-0.613, -0.308) | 35 | ⋆ |
| Paraguay | 1,440 | – | – | – | – | |
| Peru | 71,901 | -0.617 | 0.063 | (-0.742, -0.503) | 42 | ⋆ |
| Puerto Rico | 358 | – | – | – | – | |
| Saint Kitts and N. | 9 | – | – | – | – | |
| Saint Lucia | 47 | – | – | – | – | |
| Saint Martin | 1 | – | – | – | – | |
| Saint Vincent | 28 | -0.159 | 0.219 | (-0.591, 0.260) | 10 | |
| Sint Maarten | 0 | – | – | – | – | |
| Suriname | 13,727 | -0.046 | 0.134 | (-0.312, 0.219) | 3 | |
| Trinidad and Tobago | 330 | -0.265 | 0.085 | (-0.433, -0.097) | 22 | ⋆ |
| Venezuela | 42,913 | -0.120 | 0.059 | (-0.234, -3e-04) | 11 | ⋆ |
| Virgin Isl. UK | 3 | – | – | – | – | |
| Virgin Isl. US | 8 | -0.350 | 0.325 | (-0.969, 0.310) | 17 | |
| **Africa** | | | | | | |
| Angola | 6,044 | – | – | – | – | |
| Benin | 47 | – | – | – | – | |
| Burundi | 64 | -1.170 | 0.128 | (-1.420, -0.918) | 60 | ⋆ |
| Cameroon | 23,546 | -0.891 | 0.109 | (-1.120, -0.684) | 43 | ⋆ |
| CAR | 9,325 | -0.094 | 0.113 | (-0.327, 0.125) | 5 | |
| Comoros | 87 | – | – | – | – | |
| Congo | 23,945 | -0.402 | 0.088 | (-0.568, -0.233) | 31 | ⋆ |
| DRC | 125,605 | -0.160 | 0.070 | (-0.289, -0.015) | 15 | ⋆ |
| Eq. Guinea | 2,642 | -0.136 | 0.136 | (-0.385, 0.146) | 10 | |
| Ethiopia | 2,824 | -0.082 | 0.081 | (-0.241, 0.075) | 5 | |
| Gabon | 24,101 | -0.160 | 0.131 | (-0.407, 0.087) | 14 | |
| Gambia | 42 | -0.222 | 0.149 | (-0.491, 0.096) | 11 | |
| Ghana | 4,443 | -0.378 | 0.070 | (-0.522, -0.252) | 15 | ⋆ |
| Guinea | 1,213 | -0.224 | 0.078 | (-0.385, -0.077) | 14 | ⋆ |
| Guinea Bissau | 323 | -0.688 | 0.140 | (-0.999, -0.436) | 25 | ⋆ |
| Ivory Coast | 6,299 | – | – | – | – | |
| Kenya | 891 | -0.186 | 0.069 | (-0.322, -0.058) | 14 | ⋆ |
| Liberia | 8,653 | -0.491 | 0.117 | (-0.725, -0.259) | 34 | ⋆ |
| Madagascar | 5,541 | -0.285 | 0.067 | (-0.414, -0.150) | 20 | ⋆ |
| Malawi | 70 | -0.363 | 0.182 | (-0.783, -0.040) | 19 | ⋆ |
| Mauritius | 47 | -0.450 | 0.103 | (-0.650, -0.247) | 33 | ⋆ |
| Mayotte | 17 | -1.340 | 0.156 | (-1.640, -1.010) | 61 | ⋆ |
| Nigeria | 7,214 | – | – | – | – | |

Table S6: *(continued)*

| Country – study area | fc2010 (Kha) | Mean | Sd | CI 95% | P % | signif |
|---|---|---|---|---|---|---|
| Reunion | 142 | -0.385 | 0.116 | (-0.635, -0.156) | 25 | ⋆ |
| Rwanda | 195 | -1.300 | 0.120 | (-1.540, -1.070) | 71 | ⋆ |
| Senegal | 126 | -0.199 | 0.099 | (-0.381, 0.003) | 3 | |
| Sierra Leone | 2,260 | -0.247 | 0.096 | (-0.435, -0.064) | 14 | ⋆ |
| South Sudan | 201 | -0.135 | 0.099 | (-0.324, 0.041) | 4 | |
| Tanzania | 1,191 | -0.239 | 0.068 | (-0.372, -0.109) | 13 | ⋆ |
| Togo | 103 | -0.537 | 0.106 | (-0.729, -0.313) | 8 | ⋆ |
| Uganda | 1,087 | -1.190 | 0.074 | (-1.330, -1.040) | 64 | ⋆ |
| Zambia | 114 | -0.401 | 0.073 | (-0.545, -0.264) | 22 | ⋆ |
| **Asia** | | | | | | |
| Australia – Queensland | 1,876 | -0.349 | 0.068 | (-0.481, -0.213) | 18 | ⋆ |
| Bangladesh | 816 | -0.544 | 0.131 | (-0.790, -0.284) | 26 | ⋆ |
| Bhutan | 1,872 | – | – | – | – | |
| Brunei | 501 | -1.100 | 0.237 | (-1.550, -0.617) | 55 | ⋆ |
| Cambodia | 3,864 | -1.460 | 0.138 | (-1.720, -1.170) | 54 | ⋆ |
| Fiji | 958 | -0.306 | 0.130 | (-0.546, -0.045) | 23 | ⋆ |
| India – Andaman and N. | 591 | – | – | – | – | |
| India – North-East | 5,941 | -0.585 | 0.276 | (-1.110, -0.064) | 37 | ⋆ |
| India – West. Ghats | 2,704 | -0.369 | 0.102 | (-0.575, -0.166) | 19 | ⋆ |
| Indonesia | 126,473 | -0.736 | 0.062 | (-0.853, -0.609) | 46 | ⋆ |
| Laos | 9,690 | -0.442 | 0.076 | (-0.578, -0.273) | 24 | ⋆ |
| Malaysia | 22,315 | -2.080 | 0.092 | (-2.270, -1.900) | 82 | ⋆ |
| Myanmar | 15,380 | -0.167 | 0.081 | (-0.321, -0.017) | 10 | ⋆ |
| New Caledonia | 879 | – | – | – | – | |
| Papua New Guinea | 39,791 | – | – | – | – | |
| Philippines | 13,684 | -0.153 | 0.078 | (-0.309, -0.006) | 12 | ⋆ |
| Singapore | 15 | -1.350 | 0.263 | (-1.890, -0.815) | 59 | ⋆ |
| Solomon Isl. | 2,757 | – | – | – | – | |
| Sri Lanka | 1,735 | -0.516 | 0.062 | (-0.653, -0.404) | 25 | ⋆ |
| Thailand | 6,341 | -0.159 | 0.054 | (-0.258, -0.052) | 12 | ⋆ |
| Timor-Leste | 89 | -0.120 | 0.072 | (-0.261, 0.017) | 8 | |
| Vanuatu | 1,158 | -0.505 | 0.176 | (-0.848, -0.153) | 39 | ⋆ |
| Vietnam | 8,628 | -0.536 | 0.073 | (-0.685, -0.396) | 31 | ⋆ |

#### Table S7 – Effect of the distance to road on deforestation

Table S7: **Effect of the distance to road on deforestation**. We show here the estimated effect of the distance to the nearest road on the probability of deforestation for each study area. We computed the mean ("Mean"), the standard-deviation ("Sd"), and the bayesian 95% credible interval ("CI 95%") of the estimated parameter. Column "signif" indicates (with a star) that the estimated effect was negative and significantly different from zero (zero not included in the credible interval). Out of the 119 study areas, 61 showed a significant negative effect (51% of the countries). These 61 study areas accounted for 90% of the moist tropical forest in 2010 ("fc2010" in Kha).

| Country – study area | fc2010 (Kha) | Mean | Sd | CI 95% | signif |
|---|---|---|---|---|---|
| **America** | | | | | |
| Antigua and B. | 4 | – | – | – | |
| Bahamas | 115 | – | – | – | |
| Barbados | 4 | 0.000 | 0.022 | (-0.042, 0.043) | |
| Belize | 1,328 | -0.129 | 0.069 | (-0.270, 0.003) | |
| Bolivia | 30,485 | -0.306 | 0.052 | (-0.409, -0.199) | ⋆ |
| Brazil – Acre | 13,307 | -0.179 | 0.154 | (-0.462, 0.089) | |
| Brazil – Alagoas | 98 | -0.214 | 0.028 | (-0.270, -0.158) | ⋆ |
| Brazil – Amapa | 11,565 | -0.218 | 0.200 | (-0.597, 0.121) | |
| Brazil – Amazonas | 146,852 | -0.898 | 0.100 | (-1.080, -0.706) | ⋆ |
| Brazil – Bahia | 2,097 | -0.032 | 0.025 | (-0.080, 0.019) | |
| Brazil – Ceara | 48 | – | – | – | |
| Brazil – Espirito Santo | 417 | 0.002 | 0.025 | (-0.049, 0.049) | |
| Brazil – Goias | 481 | – | – | – | |
| Brazil – Maranhao | 3,930 | – | – | – | |
| Brazil – Mato Grosso | 33,283 | -0.136 | 0.052 | (-0.242, -0.040) | ⋆ |
| Brazil – Mato Grosso do Sul | 736 | – | – | – | |
| Brazil – Minas Gerais | 1,277 | -0.058 | 0.030 | (-0.117, -5e-04) | ⋆ |
| Brazil – Para | 91,982 | -0.874 | 0.141 | (-1.140, -0.622) | ⋆ |
| Brazil – Paraiba | 41 | – | – | – | |
| Brazil – Parana | 2,673 | – | – | – | |
| Brazil – Pernambouco | 119 | -0.066 | 0.028 | (-0.121, -0.010) | ⋆ |
| Brazil – Piaui | 74 | -0.019 | 0.037 | (-0.091, 0.053) | |
| Brazil – Rio de Janeiro | 736 | -0.087 | 0.036 | (-0.154, -0.017) | ⋆ |
| Brazil – Rio Grande do Norte | 25 | – | – | – | |
| Brazil – Rio Grande do Sul | 2,214 | -0.002 | 0.023 | (-0.049, 0.044) | |
| Brazil – Rondonia | 13,800 | -0.403 | 0.090 | (-0.576, -0.236) | ⋆ |
| Brazil – Roraima | 16,228 | -0.165 | 0.196 | (-0.535, 0.335) | |
| Brazil – Santa Catarina | 2,485 | – | – | – | |
| Brazil – Sao Paulo | 2,781 | -0.024 | 0.041 | (-0.099, 0.053) | |
| Brazil – Sergipe | 61 | – | – | – | |
| Brazil – Tocantins | 1,341 | -0.046 | 0.046 | (-0.146, 0.042) | |
| Colombia | 66,802 | -0.591 | 0.112 | (-0.818, -0.403) | ⋆ |
| Costa Rica | 2,277 | -0.064 | 0.055 | (-0.178, 0.032) | |
| Cuba | 1,297 | -0.126 | 0.068 | (-0.266, -0.004) | ⋆ |
| Dominica | 70 | – | – | – | |
| Dominican Rep. | 993 | -0.130 | 0.043 | (-0.210, -0.046) | ⋆ |
| Ecuador | 14,903 | -0.433 | 0.094 | (-0.613, -0.258) | ⋆ |
| El Salvador | 109 | -0.073 | 0.029 | (-0.131, -0.017) | ⋆ |
| French Guiana | 8,088 | -1.050 | 0.248 | (-1.560, -0.594) | ⋆ |
| Grenada | 22 | -0.266 | 0.122 | (-0.522, -0.037) | ⋆ |

Table S7: *(continued)*

| Country – study area | fc2010 (Kha) | Mean | Sd | CI 95% | signif |
|---|---|---|---|---|---|
| Guadeloupe | 77 | -0.089 | 0.047 | (-0.182, 0.003) | |
| Guatemala | 2,702 | -0.180 | 0.080 | (-0.327, -0.033) | ⋆ |
| Guyana | 18,489 | -0.890 | 0.198 | (-1.180, -0.448) | ⋆ |
| Haiti | 162 | -0.256 | 0.031 | (-0.319, -0.195) | ⋆ |
| Honduras | 2,993 | – | – | – | |
| Jamaica | 421 | -0.136 | 0.033 | (-0.199, -0.065) | ⋆ |
| Martinique | 70 | – | – | – | |
| Mexico | 7,390 | -0.056 | 0.039 | (-0.135, 0.021) | |
| Montserrat | 3 | – | – | – | |
| Nicaragua | 4,262 | -0.158 | 0.059 | (-0.275, -0.037) | ⋆ |
| Panama | 4,204 | -0.511 | 0.070 | (-0.655, -0.377) | ⋆ |
| Paraguay | 1,440 | -0.096 | 0.028 | (-0.153, -0.042) | ⋆ |
| Peru | 71,901 | -0.570 | 0.078 | (-0.756, -0.417) | ⋆ |
| Puerto Rico | 358 | – | – | – | |
| Saint Kitts and N. | 9 | – | – | – | |
| Saint Lucia | 47 | -0.035 | 0.064 | (-0.177, 0.086) | |
| Saint Martin | 1 | – | – | – | |
| Saint Vincent | 28 | -0.303 | 0.098 | (-0.507, -0.116) | ⋆ |
| Sint Maarten | 0 | – | – | – | |
| Suriname | 13,727 | -1.210 | 0.295 | (-1.700, -0.743) | ⋆ |
| Trinidad and Tobago | 330 | -0.106 | 0.040 | (-0.185, -0.033) | ⋆ |
| Venezuela | 42,913 | -0.585 | 0.147 | (-0.877, -0.338) | ⋆ |
| Virgin Isl. UK | 3 | – | – | – | |
| Virgin Isl. US | 8 | – | – | – | |
| **Africa** | | | | | |
| Angola | 6,044 | -0.194 | 0.037 | (-0.267, -0.122) | ⋆ |
| Benin | 47 | – | – | – | |
| Burundi | 64 | – | – | – | |
| Cameroon | 23,546 | -0.419 | 0.045 | (-0.506, -0.328) | ⋆ |
| CAR | 9,325 | -0.110 | 0.073 | (-0.237, 0.029) | |
| Comoros | 87 | – | – | – | |
| Congo | 23,945 | -0.548 | 0.091 | (-0.747, -0.385) | ⋆ |
| DRC | 125,605 | -0.375 | 0.029 | (-0.429, -0.318) | ⋆ |
| Eq. Guinea | 2,642 | -1.300 | 0.064 | (-1.420, -1.170) | ⋆ |
| Ethiopia | 2,824 | -0.082 | 0.037 | (-0.159, -0.007) | ⋆ |
| Gabon | 24,101 | -0.541 | 0.087 | (-0.705, -0.363) | ⋆ |
| Gambia | 42 | -0.327 | 0.029 | (-0.386, -0.271) | ⋆ |
| Ghana | 4,443 | -0.015 | 0.025 | (-0.066, 0.033) | |
| Guinea | 1,213 | -0.084 | 0.031 | (-0.144, -0.019) | ⋆ |
| Guinea Bissau | 323 | -0.125 | 0.073 | (-0.264, 0.017) | |
| Ivory Coast | 6,299 | -0.020 | 0.035 | (-0.088, 0.052) | |
| Kenya | 891 | -0.158 | 0.050 | (-0.259, -0.059) | ⋆ |
| Liberia | 8,653 | -0.331 | 0.049 | (-0.432, -0.239) | ⋆ |
| Madagascar | 5,541 | – | – | – | |
| Malawi | 70 | -0.515 | 0.064 | (-0.645, -0.395) | ⋆ |
| Mauritius | 47 | -0.091 | 0.025 | (-0.141, -0.043) | ⋆ |
| Mayotte | 17 | – | – | – | |
| Nigeria | 7,214 | -0.220 | 0.056 | (-0.330, -0.109) | ⋆ |
| Reunion | 142 | – | – | – | |

Table S7: *(continued)*

| Country – study area | fc2010 (Kha) | Mean | Sd | CI 95% | signif |
|---|---|---|---|---|---|
| Rwanda | 195 | -0.138 | 0.056 | (-0.253, -0.027) | ⋆ |
| Senegal | 126 | – | – | – | |
| Sierra Leone | 2,260 | -0.038 | 0.028 | (-0.093, 0.017) | |
| South Sudan | 201 | -0.203 | 0.068 | (-0.348, -0.072) | ⋆ |
| Tanzania | 1,191 | -0.184 | 0.053 | (-0.286, -0.078) | ⋆ |
| Togo | 103 | -0.271 | 0.041 | (-0.357, -0.194) | ⋆ |
| Uganda | 1,087 | -0.036 | 0.047 | (-0.129, 0.055) | |
| Zambia | 114 | -0.125 | 0.059 | (-0.238, -0.013) | ⋆ |
| **Asia** | | | | | |
| Australia – Queensland | 1,876 | 0.009 | 0.057 | (-0.110, 0.113) | |
| Bangladesh | 816 | – | – | – | |
| Bhutan | 1,872 | – | – | – | |
| Brunei | 501 | -0.087 | 0.228 | (-0.581, 0.315) | |
| Cambodia | 3,864 | -0.387 | 0.064 | (-0.507, -0.266) | ⋆ |
| Fiji | 958 | -0.124 | 0.084 | (-0.278, 0.062) | |
| India – Andaman and N. | 591 | -0.242 | 0.066 | (-0.362, -0.121) | ⋆ |
| India – North-East | 5,941 | -0.094 | 0.045 | (-0.177, -0.005) | ⋆ |
| India – West. Ghats | 2,704 | -0.170 | 0.039 | (-0.247, -0.093) | ⋆ |
| Indonesia | 126,473 | -0.433 | 0.041 | (-0.506, -0.350) | ⋆ |
| Laos | 9,690 | -0.162 | 0.035 | (-0.227, -0.093) | ⋆ |
| Malaysia | 22,315 | -0.241 | 0.067 | (-0.372, -0.112) | ⋆ |
| Myanmar | 15,380 | -0.139 | 0.041 | (-0.215, -0.056) | ⋆ |
| New Caledonia | 879 | 0.020 | 0.057 | (-0.088, 0.129) | |
| Papua New Guinea | 39,791 | -0.358 | 0.073 | (-0.494, -0.205) | ⋆ |
| Philippines | 13,684 | -0.093 | 0.031 | (-0.152, -0.029) | ⋆ |
| Singapore | 15 | -0.616 | 0.060 | (-0.740, -0.509) | ⋆ |
| Solomon Isl. | 2,757 | -0.357 | 0.165 | (-0.655, -0.056) | ⋆ |
| Sri Lanka | 1,735 | -0.068 | 0.042 | (-0.140, 0.029) | |
| Thailand | 6,341 | – | – | – | |
| Timor-Leste | 89 | -0.080 | 0.029 | (-0.139, -0.027) | ⋆ |
| Vanuatu | 1,158 | – | – | – | |
| Vietnam | 8,628 | -0.173 | 0.032 | (-0.233, -0.110) | ⋆ |

#### Tables S8–S9 – Back-transformed parameters

Table S8: **Back-transformed parameters for each study area**. We back-transformed the parameters using the mean and standard-deviation of each continuous variable for each study area. Doing so, we can use Eq. (S1) to compute the change in the probability of deforestation associated to a particular change in the explanatory variables, in their original units. To use this table of parameters, distances and elevation must be expressed in kilometers (Km), and slope must be expressed in hecto-degrees ($10^2$°). Note that the intercept is affected by the back-transformation but that the effect associated to protected areas ("pa") and the variance of the spatial random effects ("Vrho") are left unchanged.

| study area | int | pa | elev (Km) | slope ($10^2$°) | ddefor (Km) | dedge (Km) | driver (Km) | droad (Km) | dtown (Km) | Vrho |
|---|---|---|---|---|---|---|---|---|---|---|
| **America** | | | | | | | | | | |
| ATG | 2.558 | -0.710 | -4.124 | – | -3.965 | -19.187 | – | – | – | 10.00 |
| BHS | 1.277 | – | -18.839 | -8.903 | -5.458 | -6.158 | -0.002 | – | – | 8.20 |
| BRB | 2.022 | -0.323 | -5.203 | -4.797 | -2.039 | -15.536 | -0.004 | -0.001 | – | 1.99 |
| BLZ | 1.347 | -0.653 | -0.264 | -3.437 | -1.034 | -1.802 | – | -0.020 | -0.049 | 6.97 |
| BOL | 2.453 | -0.163 | -0.577 | -2.913 | -0.969 | -3.617 | – | -0.016 | -0.005 | 5.36 |
| AC | 2.555 | -0.631 | – | – | -2.458 | -2.615 | – | -0.004 | -0.006 | 4.14 |
| AL | 4.064 | -0.192 | -1.938 | -0.925 | -4.146 | -26.068 | -0.048 | -0.091 | -0.039 | 3.01 |
| AP | 2.649 | -0.126 | -7.514 | -1.937 | -1.228 | -3.191 | – | -0.005 | -0.005 | 5.22 |
| AM | 2.480 | -0.806 | -1.630 | – | -0.648 | -1.237 | -0.019 | -0.012 | -0.008 | 13.70 |
| BA | 3.305 | -0.213 | -1.042 | -3.206 | -4.447 | -7.344 | – | -0.011 | – | 4.11 |
| CE | 3.748 | -0.551 | -3.005 | – | -1.177 | -16.728 | – | – | – | 8.49 |
| ES | 2.116 | -0.193 | -1.960 | – | -4.125 | -10.097 | -0.023 | 0.001 | -0.017 | 3.20 |
| GO | 1.352 | – | -0.188 | – | -3.139 | -10.466 | -0.021 | – | -0.002 | 3.43 |
| MA | 1.661 | -0.202 | – | -2.437 | -4.344 | -3.215 | – | – | -0.018 | 3.82 |
| MT | 1.228 | -0.427 | – | – | -1.285 | -0.620 | – | -0.006 | -0.010 | 10.20 |
| MS | 1.035 | – | -0.237 | -0.251 | -1.231 | -18.842 | – | – | – | 4.07 |
| MG | 2.783 | -0.153 | -1.626 | – | -5.190 | -9.035 | -0.012 | -0.019 | -0.007 | 3.86 |
| PA | 1.880 | -1.180 | – | -0.815 | -0.785 | -1.552 | -0.002 | -0.013 | -0.005 | 8.08 |
| PB | 5.099 | -0.693 | – | -1.037 | -3.727 | -10.047 | -0.069 | – | -0.036 | 4.35 |
| PR | 1.190 | -0.339 | – | -3.659 | -2.700 | -8.130 | – | – | -0.006 | 2.54 |
| PE | 11.370 | -0.558 | -2.894 | -3.017 | -5.858 | -31.356 | -0.012 | -0.034 | -0.037 | 6.55 |
| PI | 1.874 | – | – | – | -3.453 | -18.889 | -0.003 | -0.004 | -0.009 | 4.13 |
| RJ | 2.906 | -0.220 | -0.195 | -0.097 | -6.354 | -19.109 | -0.052 | -0.036 | -0.009 | 2.40 |
| RN | 1.391 | – | – | -3.762 | -5.202 | -8.302 | – | – | – | 5.73 |
| RS | 2.515 | -0.343 | -0.414 | -6.759 | -3.980 | -27.546 | -0.001 | -0.001 | – | 1.81 |
| RO | 1.483 | -1.630 | – | -3.812 | -0.491 | -1.739 | 0.000 | -0.022 | -0.008 | 7.27 |
| RR | 2.037 | -0.841 | -2.035 | – | -1.091 | -1.017 | -0.002 | -0.003 | -0.009 | 8.49 |
| SC | 1.911 | -0.303 | – | -7.567 | -4.489 | -12.339 | – | – | -0.009 | 2.00 |
| SP | 2.651 | -0.184 | -0.482 | -3.604 | -3.658 | -7.871 | -0.003 | -0.007 | -0.007 | 3.79 |
| SE | 2.554 | – | – | – | -7.104 | -12.363 | -0.050 | – | – | 3.63 |
| TO | 0.249 | -0.035 | – | – | -2.011 | -0.191 | -0.009 | -0.005 | – | 4.43 |
| COL | 2.332 | -0.449 | -1.091 | -3.682 | -1.084 | -0.676 | – | -0.009 | -0.021 | 5.62 |
| CRI | 2.042 | – | -0.089 | -3.323 | -3.299 | -9.859 | -0.005 | -0.010 | -0.020 | 2.18 |
| CUB | 1.060 | – | -0.912 | -1.257 | -2.231 | -4.373 | -0.007 | -0.016 | -0.006 | 4.77 |
| DMA | 2.163 | – | -0.967 | -2.054 | -0.934 | -5.433 | – | – | -0.245 | 3.26 |
| DOM | 1.951 | -0.331 | -0.431 | -0.856 | -4.952 | -10.476 | -0.001 | -0.028 | -0.001 | 2.85 |
| ECU | 2.787 | -0.520 | -0.106 | -3.845 | -1.178 | -5.817 | – | -0.021 | -0.017 | 2.53 |
| SLV | 2.613 | -0.476 | -0.618 | -2.548 | -3.853 | -18.919 | – | -0.049 | -0.013 | 3.38 |
| GUF | 4.366 | -0.835 | -7.708 | -3.654 | -0.386 | -1.232 | – | -0.023 | -0.031 | 18.80 |
| GRD | 4.269 | -0.396 | -7.059 | – | -4.878 | -17.737 | – | -0.274 | -0.116 | 14.20 |
| GLP | 2.039 | – | -0.712 | -2.517 | -1.714 | -16.808 | -0.020 | -0.066 | -0.037 | 2.61 |
| GTM | 1.664 | -0.229 | -0.649 | -2.668 | -2.798 | -0.508 | -0.003 | -0.019 | -0.003 | 3.15 |
| GUY | 3.250 | -0.708 | -1.990 | -4.576 | -0.166 | -2.296 | – | -0.018 | -0.014 | 13.70 |

Table S8: *(continued)*

| study area | int | pa | elev (Km) | slope ($10^2$°) | ddefor (Km) | dedge (Km) | driver (Km) | droad (Km) | dtown (Km) | Vrho |
|---|---|---|---|---|---|---|---|---|---|---|
| HTI | 1.969 | -0.038 | – | -1.129 | -7.618 | -17.132 | -0.049 | -0.078 | -0.007 | 2.87 |
| HND | 1.182 | -0.364 | -1.759 | -3.511 | -0.713 | -0.529 | -0.006 | – | – | 4.46 |
| JAM | 2.408 | -0.003 | -0.905 | -1.675 | -5.545 | -9.247 | -0.008 | -0.067 | -0.035 | 2.66 |
| MTQ | 2.447 | -0.075 | -3.560 | -0.750 | -1.548 | -19.757 | – | – | – | 2.66 |
| MEX | 0.970 | -0.228 | -0.371 | -2.717 | -2.857 | -4.082 | – | -0.009 | -0.004 | 3.59 |
| MSR | 0.714 | – | -1.346 | – | -3.629 | -37.135 | – | – | – | 4.81 |
| NIC | 0.106 | – | -1.402 | -1.605 | -0.284 | -1.436 | – | -0.010 | – | 3.19 |
| PAN | 2.151 | -0.455 | -0.389 | -3.102 | -2.530 | -3.001 | – | -0.030 | – | 2.97 |
| PRY | 0.774 | – | -0.761 | -3.761 | -3.790 | -1.111 | -0.001 | -0.019 | – | 3.93 |
| PER | 4.082 | -0.617 | -0.940 | -3.981 | -0.967 | -1.322 | -0.004 | -0.014 | -0.027 | 6.44 |
| PRI | 1.697 | – | -1.600 | -0.862 | -1.663 | -7.575 | – | – | -0.024 | 4.32 |
| KNA | 1.275 | – | -5.366 | -1.959 | -0.506 | -2.931 | – | – | – | 1.62 |
| LCA | 2.153 | – | -2.175 | -0.882 | -1.738 | -10.080 | – | -0.027 | -0.052 | 2.97 |
| MAF | 3.246 | – | – | – | -4.740 | -19.250 | -0.351 | – | – | 8.11 |
| VCT | 3.544 | -0.159 | -3.215 | -2.167 | -2.031 | -7.379 | – | -0.183 | -0.028 | 3.30 |
| SXM | 4.323 | – | – | -8.031 | -6.663 | -14.834 | -0.133 | – | -0.719 | 10.00 |
| SUR | 2.359 | -0.046 | -5.104 | -3.935 | -0.288 | -0.736 | – | -0.020 | -0.014 | 12.50 |
| TTO | 2.339 | -0.265 | -2.302 | -2.393 | -3.074 | -8.572 | -0.011 | -0.055 | -0.040 | 2.16 |
| VEN | 2.877 | -0.120 | -0.854 | -1.092 | -0.626 | -3.801 | -0.010 | -0.007 | -0.014 | 6.59 |
| VGB | 3.955 | – | -2.957 | -2.658 | -7.305 | -14.010 | – | – | – | 8.04 |
| VIR | 1.208 | -0.350 | – | -2.352 | -1.401 | -7.461 | – | – | – | 7.64 |
| **Africa** | | | | | | | | | | |
| AGO | 2.012 | – | -0.107 | -1.890 | -3.736 | -6.599 | -0.002 | -0.017 | -0.004 | 4.00 |
| BEN | 4.410 | – | – | -13.207 | -1.212 | -25.389 | – | – | – | 8.05 |
| BDI | 4.709 | -1.170 | -1.027 | – | -7.115 | -12.746 | -0.025 | – | -0.031 | 6.39 |
| CMR | 3.117 | -0.891 | – | -3.409 | -0.629 | -1.679 | -0.004 | -0.048 | -0.040 | 5.96 |
| CAF | 3.070 | -0.094 | – | -1.107 | -3.620 | -1.343 | -0.010 | -0.007 | -0.028 | 4.61 |
| COM | 2.961 | – | -0.378 | -3.925 | – | -38.750 | – | – | – | 7.22 |
| COG | 2.460 | -0.402 | -0.099 | -1.779 | -0.304 | -1.428 | -0.014 | -0.024 | -0.018 | 8.71 |
| COD | 2.330 | -0.160 | – | – | -2.221 | -1.882 | – | -0.024 | -0.018 | 4.52 |
| GNQ | 2.770 | -0.136 | -1.224 | -6.923 | -0.014 | -2.320 | -0.010 | -0.367 | -0.010 | 8.70 |
| ETH | 2.076 | -0.082 | -0.113 | -3.524 | -2.713 | -5.952 | – | -0.013 | -0.011 | 3.55 |
| GAB | 2.664 | -0.160 | -2.250 | -5.677 | -0.232 | -1.678 | – | -0.028 | -0.032 | 14.60 |
| GMB | 2.030 | -0.222 | – | -12.123 | -6.070 | -8.739 | – | -0.137 | -0.044 | 5.23 |
| GHA | 2.048 | -0.378 | -1.241 | -5.014 | -0.935 | -3.447 | – | -0.006 | -0.017 | 2.06 |
| GIN | 1.598 | -0.224 | -0.311 | -0.211 | -5.232 | -5.554 | – | -0.018 | – | 1.88 |
| GNB | 2.772 | -0.688 | – | – | -7.007 | -11.120 | – | -0.012 | -0.008 | 3.13 |
| CIV | 1.029 | – | – | -4.006 | -1.090 | -0.655 | – | -0.003 | -0.011 | 2.61 |
| KEN | 2.505 | -0.186 | -0.638 | -0.905 | -1.970 | -10.784 | -0.009 | -0.026 | -0.009 | 6.97 |
| LBR | 1.575 | -0.491 | – | -4.035 | -0.915 | -3.457 | – | -0.041 | -0.040 | 2.06 |
| MDG | 1.871 | -0.285 | -1.176 | -2.307 | -3.994 | -2.523 | – | – | -0.016 | 3.43 |
| MWI | 2.617 | -0.363 | -0.316 | -2.293 | -5.284 | -3.091 | -0.052 | -0.111 | 0.000 | 18.70 |
| MUS | 1.605 | -0.450 | -0.822 | -2.421 | -2.128 | -19.086 | -0.011 | -0.102 | -0.017 | 1.37 |
| MYT | 2.020 | -1.340 | – | – | -3.575 | -16.398 | -0.033 | – | -0.041 | 2.77 |
| NGA | 2.329 | – | – | -0.348 | -1.601 | -1.520 | – | -0.023 | – | 5.63 |
| REU | 2.148 | -0.385 | – | -1.904 | -0.160 | -17.669 | – | – | – | 2.76 |
| RWA | 4.109 | -1.300 | -0.726 | -2.592 | -3.378 | -14.877 | – | -0.057 | – | 3.98 |
| SEN | 2.994 | -0.199 | -3.323 | -12.669 | -2.812 | -8.487 | – | – | -0.033 | 6.03 |
| SLE | 1.001 | -0.247 | – | – | -4.649 | -5.033 | – | -0.008 | -0.013 | 1.31 |
| SSD | 2.339 | -0.135 | -1.038 | 0.194 | -1.242 | -7.670 | – | -0.013 | – | 4.16 |
| TZA | 3.338 | -0.239 | -0.576 | -1.054 | -2.476 | -13.949 | -0.011 | -0.030 | -0.010 | 5.93 |
| TGO | 4.044 | -0.537 | -0.985 | -1.958 | -4.937 | -18.407 | – | -0.073 | – | 4.68 |
| UGA | 1.799 | -1.190 | – | – | -3.703 | -4.299 | -0.009 | -0.010 | -0.035 | 4.72 |

Table S8: *(continued)*

| study area | int | pa | elev (Km) | slope ($10^2$°) | ddefor (Km) | dedge (Km) | driver (Km) | droad (Km) | dtown (Km) | Vrho |
|---|---|---|---|---|---|---|---|---|---|---|
| ZMB | 0.808 | -0.401 | – | – | -1.383 | -13.980 | – | -0.014 | -0.005 | 9.53 |
| **Asia** | | | | | | | | | | |
| QLD | 1.812 | -0.349 | – | -1.516 | -2.393 | -3.886 | – | 0.001 | – | 5.56 |
| BGD | 1.460 | -0.544 | -0.807 | -1.947 | -2.055 | -10.087 | -0.024 | – | – | 4.47 |
| BTN | 2.309 | – | -0.063 | -0.815 | -6.571 | -15.148 | -0.017 | – | -0.008 | 1.19 |
| BRN | 2.902 | -1.100 | -7.792 | -7.695 | -1.818 | – | – | -0.013 | -0.049 | 15.10 |
| KHM | 3.637 | -1.460 | -4.035 | -3.153 | -1.803 | -0.665 | – | -0.041 | -0.022 | 9.40 |
| FJI | 2.306 | -0.306 | -1.188 | -3.469 | -0.300 | -5.792 | – | -0.015 | -0.020 | 5.21 |
| AN | 2.657 | – | -3.730 | -4.336 | -0.779 | -14.369 | – | -0.022 | – | 4.55 |
| NE | 2.278 | -0.585 | -0.314 | -2.493 | -6.283 | -8.612 | -0.009 | -0.011 | -0.006 | 2.39 |
| WG | 2.624 | -0.369 | -1.037 | -0.971 | -4.358 | -19.244 | – | -0.077 | -0.017 | 1.93 |
| IDN | 1.559 | -0.736 | -0.654 | -7.390 | -0.895 | -1.632 | -0.006 | -0.020 | -0.012 | 8.13 |
| LAO | 2.855 | -0.442 | -1.083 | -3.908 | -4.876 | -4.924 | – | -0.022 | -0.030 | 3.08 |
| MYS | 1.689 | -2.080 | -1.154 | -6.327 | -0.946 | -2.997 | – | -0.018 | – | 7.84 |
| MMR | 2.377 | -0.167 | -0.754 | -2.199 | -4.909 | -6.342 | -0.021 | -0.010 | -0.002 | 3.10 |
| NCL | 3.148 | – | -1.447 | -1.018 | -0.820 | -26.598 | – | 0.003 | -0.009 | 4.11 |
| PNG | 1.798 | – | -0.704 | -5.312 | -0.022 | -2.988 | -0.007 | -0.008 | -0.007 | 7.46 |
| PHL | 1.654 | -0.153 | -0.565 | -4.569 | -3.528 | -6.252 | – | -0.018 | -0.040 | 2.97 |
| SGP | 3.028 | -1.350 | -6.453 | -5.018 | -6.100 | -19.104 | – | -0.616 | – | 6.01 |
| SLB | 1.497 | – | -3.765 | -6.472 | – | -0.710 | -0.009 | -0.005 | – | 5.86 |
| LKA | 2.326 | -0.516 | -0.286 | -4.131 | -5.927 | -11.489 | -0.012 | -0.022 | -0.052 | 1.71 |
| THA | 2.679 | -0.159 | -2.035 | -0.968 | -5.197 | -13.353 | -0.002 | – | – | 2.07 |
| TLS | 2.450 | -0.120 | -0.401 | -1.072 | -10.936 | -17.230 | -0.011 | -0.028 | -0.029 | 1.60 |
| VUT | 1.565 | -0.505 | -1.736 | -3.475 | -0.112 | -6.924 | – | – | 0.001 | 18.10 |
| VNM | 2.477 | -0.536 | -0.645 | -4.106 | -5.390 | -5.369 | -0.012 | -0.052 | -0.012 | 3.12 |

Table S9: **Back-transformed parameters per region and continent**. We back-transformed the parameters using the mean and standard-deviation of each continuous variable for each study area. We then used the forest cover in 2010 to compute the back-transformed parameter estimate weighted mean per region. Doing so, we can use Eq. (S1) to compute the change in the probability of deforestation associated with a particular change in the explanatory variables, in their original units. To use this table of parameters, distances and elevation must be expressed in kilometers (Km), and slope must be expressed in hecto-degrees ($10^2$°).

| Region | int | pa | elev (Km) | slope ($10^2$°) | ddefor (Km) | dedge (Km) | driver (Km) | droad (Km) | dtown (Km) | Vrho |
|---|---|---|---|---|---|---|---|---|---|---|
| India | 2.403 | -0.484 | -0.744 | -2.165 | -5.368 | -12.093 | -0.006 | -0.031 | -0.009 | 2.393 |
| Brazil | 2.126 | -0.837 | -1.055 | -0.633 | -1.049 | -1.949 | -0.009 | -0.011 | -0.007 | 10.093 |
| America | 2.470 | -0.636 | -1.173 | -1.849 | -1.026 | -2.097 | -0.006 | -0.012 | -0.012 | 8.549 |
| Africa | 2.385 | -0.259 | -0.292 | -1.602 | -1.758 | -2.212 | -0.002 | -0.028 | -0.021 | 5.819 |
| Asia | 1.841 | -0.625 | -0.848 | -5.739 | -1.769 | -3.629 | -0.006 | -0.018 | -0.011 | 6.676 |
| All continents | 2.308 | -0.551 | -0.905 | -2.683 | -1.357 | -2.472 | -0.005 | -0.017 | -0.014 | 7.519 |

#### Tables S10–S11 – Mathematical formulas for accuracy indices

Table S10: **Confusion matrix used to compute accuracy indices**. A confusion matrix can be computed to compare model predictions with observations.

| | | Observations | | Total |
|---|---|---|---|---|
| | | 0 (non-deforested) | 1 (deforested) | |
| Predictions | 0 | $n_{00}$ | $n_{01}$ | $n_{0+}$ |
| | 1 | $n_{10}$ | $n_{11}$ | $n_{1+}$ |
| Total | | $n_{+0}$ | $n_{+1}$ | $n$ |

Table S11: **Formulas used to compute accuracy indices**. Several accuracy indices can be computed from the confusion matrix to estimate and compare models' predictive performance. We followed the definitions of Pontius et al. (2008) for the FOM and Liu et al. (2011) for the other indices. Note that the AUC relies on the predicted probabilities for observations 0 (non-deforested) and 1 (deforested), not on the confusion matrix.

| Index | Formula |
|---|---|
| Overall Accuracy | $\text{OA} = (n_{11} + n_{00})/n$ |
| Figure Of Merit | $\text{FOM} = n_{11}/(n_{11} + n_{10} + n_{01})$ |
| Sensitivity | $\text{Sen} = n_{11}/(n_{11} + n_{01})$ |
| Specificity | $\text{Spe} = n_{00}/(n_{00} + n_{10})$ |
| True Skill Statistics | $\text{TSS} = \text{Sen} + \text{Spe} - 1$ |
| Area Under ROC Curve | $\text{AUC} = 1/(n_{+1}n_{+0}) \sum_{i=1}^{n_{+0}} \sum_{j=1}^{n_{+1}} \phi(\delta_i, \theta_j)$ |
| | where $\phi(\delta_i, \theta_j)$ equals 1 if $\theta_j > \delta_i$, 1/2 if $\theta_j = \delta_i$, and 0 otherwise |
| | $\delta_i$ and $\theta_j$ are the predicted probabilities for $Y_i = 0$ and $Y_j = 1$ |

#### Tables S12–S13 – Accuracy indices

Table S12: **Accuracy indices' weighted means for the three statistical models**. Accuracy indices were averaged across study areas using forest cover areas in 2010 as weights. D: percentage of deviance explained, AUC: Area Under ROC Curve, OA: overall accuracy, FOM: Figure Of Merit, TSS: True Skill Statistics. Averaged accuracy indices were computed for the three statistical models: “glm”, “icar”, and “rf” model. While the “rf” model has a higher percentage of deviance explained in average (higher goodness-of-fit), the “icar” model has higher values of accuracy indices from the cross-validation (higher predictive power).

| Model | D | AUC | OA | FOM | TSS |
|---|---|---|---|---|---|
| glm | 39.3 | 88.2 | 80.8 | 68.1 | 61.8 |
| icar | 53.3 | 91.7 | 84.7 | 73.6 | 69.2 |
| rf | 87.1 | 90.8 | 83.8 | 72.3 | 67.5 |

Table S13: **Accuracy indices' weighted means for the three statistical models by continent**. Accuracy indices were averaged across study areas for each continent using forest cover areas in 2010 as weights. D: percentage of deviance explained, AUC: Area Under ROC Curve, OA: overall accuracy, FOM: Figure Of Merit, TSS: True Skill Statistics. Averaged accuracy indices were computed for the three statistical models: "glm", "icar", and "rf" model.

| Continent | Model | D | AUC | OA | FOM | TSS |
|---|---|---|---|---|---|---|
| | glm | 42.6 | 89.4 | 82.1 | 69.9 | 64.3 |
| **America** | icar | 57.0 | 92.8 | 86.2 | 75.8 | 72.1 |
| | rf | 88.6 | 92.5 | 85.9 | 75.4 | 71.4 |
| | glm | 40.3 | 88.6 | 81.5 | 69.0 | 63.1 |
| **Africa** | icar | 51.7 | 91.4 | 84.3 | 72.8 | 68.2 |
| | rf | 86.3 | 89.7 | 82.3 | 70.4 | 65.0 |
| | glm | 30.4 | 84.8 | 77.0 | 62.9 | 54.5 |
| **Asia** | icar | 46.0 | 89.5 | 81.6 | 69.0 | 63.1 |
| | rf | 84.4 | 87.5 | 79.9 | 66.7 | 60.3 |

#### Tables S14–S15 – Past forest cover change

Table S14: **Past forest cover change for each study area**. Forest cover areas are given in thousand hectares (Kha) for the years 2000, 2010 and 2020 ("fc2000", "fc2010", and "fc2020", respectively). The mean annual deforested area $d$ for the ten-year period 2010–2020 is given in hectare per year (ha/yr). The corresponding mean annual deforestation rate $p$ is also provided in percent per year (%/yr), with one decimal precision, to be able to compare the intensity of deforestation between study areas.

| Country – study area | fc2000 (Kha) | fc2010 (Kha) | fc2020 (Kha) | $d$ (ha/yr) | $p$ (%/yr) |
|---|---|---|---|---|---|
| **America** | | | | | |
| Antigua and B. | 4 | 4 | 3 | 52 | 1.6 |
| Bahamas | 152 | 115 | 98 | 1,657 | 1.5 |
| Barbados | 4 | 4 | 3 | 64 | 1.7 |
| Belize | 1,421 | 1,328 | 1,198 | 12,996 | 1.0 |
| Bolivia | 32,621 | 30,485 | 28,739 | 174,616 | 0.6 |
| Brazil – Acre | 13,912 | 13,307 | 12,860 | 44,684 | 0.3 |
| Brazil – Alagoas | 112 | 98 | 89 | 901 | 1.0 |
| Brazil – Amapa | 11,730 | 11,565 | 11,458 | 10,677 | 0.1 |
| Brazil – Amazonas | 148,095 | 146,852 | 145,346 | 150,663 | 0.1 |
| Brazil – Bahia | 2,521 | 2,097 | 1,936 | 16,103 | 0.8 |
| Brazil – Ceara | 57 | 48 | 40 | 849 | 1.9 |
| Brazil – Espirito Santo | 487 | 417 | 387 | 2,945 | 0.7 |
| Brazil – Goias | 644 | 481 | 358 | 12,241 | 2.9 |
| Brazil – Maranhao | 5,638 | 3,930 | 3,259 | 67,024 | 1.9 |
| Brazil – Mato Grosso | 40,368 | 33,283 | 30,333 | 294,945 | 0.9 |
| Brazil – Mato Grosso do Sul | 871 | 736 | 654 | 8,193 | 1.2 |
| Brazil – Minas Gerais | 1,824 | 1,277 | 954 | 32,323 | 2.9 |
| Brazil – Para | 100,168 | 91,982 | 87,764 | 421,863 | 0.5 |
| Brazil – Paraiba | 46 | 41 | 38 | 308 | 0.8 |
| Brazil – Parana | 3,202 | 2,673 | 2,475 | 19,780 | 0.8 |
| Brazil – Pernambouco | 138 | 119 | 109 | 936 | 0.8 |
| Brazil – Piaui | 104 | 74 | 55 | 1,967 | 3.0 |
| Brazil – Rio de Janeiro | 820 | 736 | 694 | 4,143 | 0.6 |
| Brazil – Rio Grande do Norte | 31 | 25 | 22 | 312 | 1.3 |
| Brazil – Rio Grande do Sul | 2,533 | 2,214 | 2,074 | 14,004 | 0.7 |
| Brazil – Rondonia | 16,500 | 13,800 | 12,436 | 136,426 | 1.0 |
| Brazil – Roraima | 16,715 | 16,228 | 15,593 | 63,551 | 0.4 |
| Brazil – Santa Catarina | 2,934 | 2,485 | 2,355 | 13,020 | 0.5 |
| Brazil – Sao Paulo | 3,029 | 2,781 | 2,653 | 12,862 | 0.5 |
| Brazil – Sergipe | 74 | 61 | 54 | 656 | 1.1 |
| Brazil – Tocantins | 1,731 | 1,341 | 954 | 38,777 | 3.4 |
| Colombia | 70,049 | 66,802 | 64,254 | 254,796 | 0.4 |
| Costa Rica | 2,412 | 2,277 | 2,132 | 14,520 | 0.7 |
| Cuba | 1,521 | 1,297 | 1,155 | 14,086 | 1.1 |
| Dominica | 71 | 70 | 67 | 254 | 0.4 |
| Dominican Rep. | 1,254 | 993 | 871 | 12,164 | 1.3 |
| Ecuador | 15,460 | 14,903 | 14,391 | 51,193 | 0.3 |
| El Salvador | 129 | 109 | 97 | 1,220 | 1.2 |
| French Guiana | 8,118 | 8,088 | 8,064 | 2,390 | 0.0 |
| Grenada | 26 | 22 | 19 | 264 | 1.3 |
| Guadeloupe | 80 | 77 | 73 | 370 | 0.5 |

Table S14: *(continued)*

| Country – study area | fc2000 (Kha) | fc2010 (Kha) | fc2020 (Kha) | $d$ (ha/yr) | $p$ (%/yr) |
|---|---|---|---|---|---|
| Guatemala | 3,449 | 2,702 | 2,275 | 42,648 | 1.7 |
| Guyana | 18,618 | 18,489 | 18,366 | 12,342 | 0.1 |
| Haiti | 247 | 162 | 122 | 4,001 | 2.8 |
| Honduras | 3,384 | 2,993 | 2,569 | 42,409 | 1.5 |
| Jamaica | 472 | 421 | 395 | 2,572 | 0.6 |
| Martinique | 73 | 70 | 66 | 400 | 0.6 |
| Mexico | 9,098 | 7,390 | 6,264 | 112,620 | 1.6 |
| Montserrat | 3 | 3 | 3 | 7 | 0.2 |
| Nicaragua | 4,926 | 4,262 | 3,408 | 85,457 | 2.2 |
| Panama | 4,423 | 4,204 | 4,012 | 19,202 | 0.5 |
| Paraguay | 2,359 | 1,440 | 1,090 | 34,950 | 2.7 |
| Peru | 73,255 | 71,901 | 70,775 | 112,691 | 0.2 |
| Puerto Rico | 423 | 358 | 321 | 3,688 | 1.1 |
| Saint Kitts and N. | 9 | 9 | 9 | 34 | 0.4 |
| Saint Lucia | 47 | 47 | 44 | 261 | 0.6 |
| Saint Martin | 1 | 1 | 0 | 24 | 5.4 |
| Saint Vincent | 29 | 28 | 27 | 106 | 0.4 |
| Sint Maarten | 0 | 0 | 0 | 10 | 6.0 |
| Suriname | 13,814 | 13,727 | 13,624 | 10,316 | 0.1 |
| Trinidad and Tobago | 349 | 330 | 307 | 2,355 | 0.7 |
| Venezuela | 44,743 | 42,913 | 41,430 | 148,279 | 0.4 |
| Virgin Isl. UK | 4 | 3 | 2 | 96 | 4.0 |
| Virgin Isl. US | 9 | 8 | 7 | 137 | 1.8 |
| **Africa** | | | | | |
| Angola | 7,065 | 6,044 | 5,341 | 70,293 | 1.2 |
| Benin | 77 | 47 | 32 | 1,463 | 3.7 |
| Burundi | 104 | 64 | 55 | 856 | 1.4 |
| Cameroon | 23,887 | 23,546 | 22,840 | 70,641 | 0.3 |
| CAR | 9,808 | 9,325 | 8,854 | 47,114 | 0.5 |
| Comoros | 87 | 87 | 83 | 263 | 0.3 |
| Congo | 24,090 | 23,945 | 23,428 | 51,689 | 0.2 |
| DRC | 131,298 | 125,605 | 118,283 | 732,153 | 0.6 |
| Eq. Guinea | 2,647 | 2,642 | 2,612 | 2,941 | 0.1 |
| Ethiopia | 3,799 | 2,824 | 2,214 | 61,036 | 2.4 |
| Gabon | 24,129 | 24,101 | 23,985 | 11,595 | 0.0 |
| Gambia | 49 | 42 | 31 | 566 | 1.4 |
| Ghana | 4,932 | 4,443 | 3,345 | 109,799 | 2.8 |
| Guinea | 1,895 | 1,213 | 837 | 37,643 | 3.6 |
| Guinea Bissau | 398 | 323 | 272 | 5,078 | 1.7 |
| Ivory Coast | 7,734 | 6,299 | 3,954 | 234,522 | 4.6 |
| Kenya | 1,199 | 891 | 768 | 12,324 | 1.5 |
| Liberia | 8,906 | 8,653 | 7,908 | 74,471 | 0.9 |
| Madagascar | 7,024 | 5,541 | 4,574 | 96,645 | 1.9 |
| Malawi | 113 | 70 | 39 | 3,147 | 5.8 |
| Mauritius | 50 | 47 | 43 | 347 | 0.8 |
| Mayotte | 18 | 17 | 12 | 511 | 3.5 |
| Nigeria | 7,770 | 7,214 | 6,197 | 101,634 | 1.5 |
| Reunion | 142 | 142 | 134 | 722 | 0.5 |
| Rwanda | 284 | 195 | 160 | 3,434 | 1.9 |

Table S14: *(continued)*

| Country – study area | fc2000 (Kha) | fc2010 (Kha) | fc2020 (Kha) | *d* (ha/yr) | *p* (%/yr) |
|---|---|---|---|---|---|
| Senegal | 136 | 126 | 109 | 1,736 | 1.5 |
| Sierra Leone | 3,440 | 2,260 | 1,414 | 84,563 | 4.6 |
| South Sudan | 265 | 201 | 167 | 3,493 | 1.9 |
| Tanzania | 1,431 | 1,191 | 1,084 | 10,689 | 0.9 |
| Togo | 160 | 103 | 66 | 3,693 | 4.3 |
| Uganda | 1,879 | 1,087 | 758 | 32,878 | 3.5 |
| Zambia | 177 | 114 | 81 | 3,294 | 3.4 |
| **Asia** | | | | | |
| Australia – Queensland | 2,055 | 1,876 | 1,764 | 11,131 | 0.6 |
| Bangladesh | 963 | 816 | 771 | 4,489 | 0.6 |
| Bhutan | 1,990 | 1,872 | 1,802 | 7,022 | 0.4 |
| Brunei | 511 | 501 | 494 | 668 | 0.1 |
| Cambodia | 4,804 | 3,864 | 2,753 | 111,024 | 3.3 |
| Fiji | 1,005 | 958 | 924 | 3,300 | 0.4 |
| India – Andaman and N. | 612 | 591 | 572 | 1,875 | 0.3 |
| India – North-East | 7,023 | 5,941 | 5,560 | 38,134 | 0.7 |
| India – West. Ghats | 3,144 | 2,704 | 2,236 | 46,740 | 1.9 |
| Indonesia | 139,358 | 126,473 | 117,072 | 939,867 | 0.8 |
| Laos | 11,607 | 9,690 | 8,308 | 138,221 | 1.5 |
| Malaysia | 25,676 | 22,315 | 20,147 | 216,762 | 1.0 |
| Myanmar | 18,279 | 15,380 | 13,728 | 165,195 | 1.1 |
| New Caledonia | 905 | 879 | 855 | 2,425 | 0.3 |
| Papua New Guinea | 40,366 | 39,791 | 39,304 | 48,691 | 0.1 |
| Philippines | 14,756 | 13,684 | 12,753 | 93,052 | 0.7 |
| Singapore | 17 | 15 | 14 | 147 | 1.0 |
| Solomon Isl. | 2,762 | 2,757 | 2,739 | 1,751 | 0.1 |
| Sri Lanka | 2,088 | 1,735 | 1,594 | 14,060 | 0.8 |
| Thailand | 7,188 | 6,341 | 5,815 | 52,603 | 0.9 |
| Timor-Leste | 131 | 89 | 78 | 1,173 | 1.4 |
| Vanuatu | 1,158 | 1,158 | 1,152 | 564 | 0.0 |
| Vietnam | 10,692 | 8,628 | 7,599 | 102,909 | 1.3 |

Table S15: **Past forest cover change per region and continent**. Areas of forest cover are given in thousand hectares (Kha) for the years 2000, 2010 and 2020 ("fc2000", "fc2010", and "fc2020", respectively). The mean annual deforested area $d$ for the ten-year period 2010–2020 is given in hectare per year (ha/yr). The corresponding mean annual deforestation rate $p$ is also provided in percent per year (%/yr), with one decimal precision, to be able to compare the intensity of deforestation between study areas. Estimates for America include Brazil, and estimates for Asia include India. Around 6.4 Mha (64,000 $km^2$, about half the size of Greece or the size of West Virginia) of natural old-growth moist tropical forest have been disappearing each year in the period 2010–2020.

| Region | fc2000 (Kha) | fc2010 (Kha) | fc2020 (Kha) | $d$ (ha/yr) | $p$ (%/yr) |
|---|---|---|---|---|---|
| India | 10,780 | 9,236 | 8,368 | 86,749 | 1.0 |
| Brazil | 374,282 | 348,650 | 334,948 | 1,370,153 | 0.4 |
| America | 687,339 | 646,685 | 621,229 | 2,545,400 | 0.4 |
| Africa | 274,993 | 258,401 | 239,681 | 1,871,233 | 0.7 |
| Asia | 297,090 | 268,058 | 248,035 | 2,001,803 | 0.8 |
| All continents | 1,259,422 | 1,173,144 | 1,108,945 | 6,418,436 | 0.6 |

#### Tables S16–17 – Forest cover projections

Table S16: **Forest cover projections for each study area**. Projected areas of forest cover are given in thousand hectares (Kha) for four years in the future (2040, 2060, 2080, and 2100). Projections were made using the forest cover in 2020 and the mean annual deforested area in the ten-year period 2010–2020 ("fc2000" and *d* respectively in Table S14), assuming a "business-as-usual" scenario of deforestation. Column "loss21" indicates the projected percentage of forest cover loss during the 21[st] century (2100 vs. 2000). Column "yrdis" indicates the estimated year at which all the forest of the study area will have disappeared.

| Country – study area | fc2040 (Kha) | fc2060 (Kha) | fc2080 (Kha) | fc2100 (Kha) | loss21 (%) | yrdis |
|---|---|---|---|---|---|---|
| **America** | | | | | | |
| Antigua and B. | 2 | 1 | 0 | 0 | 100 | 2078 |
| Bahamas | 65 | 32 | 0 | 0 | 100 | 2079 |
| Barbados | 2 | 1 | 0 | 0 | 100 | 2072 |
| Belize | 938 | 678 | 418 | 158 | 89 | 2112 |
| Bolivia | 25,246 | 21,754 | 18,262 | 14,769 | 55 | 2184 |
| Brazil – Acre | 11,967 | 11,016 | 9,943 | 8,781 | 37 | 2160 |
| Brazil – Alagoas | 71 | 0 | 0 | 0 | 100 | 2060 |
| Brazil – Amapa | 11,245 | 10,974 | 10,581 | 10,099 | 14 | 2173 |
| Brazil – Amazonas | 142,332 | 139,262 | 136,069 | 132,788 | 10 | 2264 |
| Brazil – Bahia | 1,614 | 1,235 | 733 | 143 | 94 | 2105 |
| Brazil – Ceara | 23 | 0 | 0 | 0 | 100 | 2051 |
| Brazil – Espirito Santo | 328 | 212 | 0 | 0 | 100 | 2079 |
| Brazil – Goias | 113 | 0 | 0 | 0 | 100 | 2049 |
| Brazil – Maranhao | 1,919 | 521 | 0 | 0 | 100 | 2068 |
| Brazil – Mato Grosso | 24,434 | 18,478 | 12,400 | 6,233 | 85 | 2120 |
| Brazil – Mato Grosso do Sul | 490 | 269 | 0 | 0 | 100 | 2077 |
| Brazil – Minas Gerais | 308 | 0 | 0 | 0 | 100 | 2050 |
| Brazil – Para | 79,326 | 70,832 | 62,215 | 53,510 | 47 | 2189 |
| Brazil – Paraiba | 32 | 0 | 0 | 0 | 100 | 2054 |
| Brazil – Parana | 2,080 | 1,627 | 1,052 | 388 | 88 | 2110 |
| Brazil – Pernambouco | 91 | 15 | 0 | 0 | 100 | 2063 |
| Brazil – Piaui | 15 | 0 | 0 | 0 | 100 | 2047 |
| Brazil – Rio de Janeiro | 612 | 472 | 209 | 0 | 100 | 2093 |
| Brazil – Rio Grande do Norte | 15 | 0 | 0 | 0 | 100 | 2051 |
| Brazil – Rio Grande do Sul | 1,794 | 1,457 | 997 | 449 | 82 | 2112 |
| Brazil – Rondonia | 9,707 | 6,922 | 4,014 | 1,017 | 94 | 2107 |
| Brazil – Roraima | 14,322 | 12,993 | 11,543 | 10,004 | 40 | 2161 |
| Brazil – Santa Catarina | 2,094 | 1,777 | 1,337 | 808 | 72 | 2119 |
| Brazil – Sao Paulo | 2,395 | 2,081 | 1,645 | 1,119 | 63 | 2122 |
| Brazil – Sergipe | 41 | 0 | 0 | 0 | 100 | 2055 |
| Brazil – Tocantins | 178 | 0 | 0 | 0 | 100 | 2045 |
| Colombia | 59,158 | 54,062 | 48,966 | 43,871 | 37 | 2272 |
| Costa Rica | 1,841 | 1,551 | 1,260 | 970 | 60 | 2166 |
| Cuba | 873 | 592 | 310 | 28 | 98 | 2101 |
| Dominica | 62 | 57 | 52 | 47 | 33 | 2285 |
| Dominican Rep. | 628 | 385 | 141 | 0 | 100 | 2091 |
| Ecuador | 13,367 | 12,343 | 11,319 | 10,295 | 33 | 2301 |
| El Salvador | 72 | 48 | 23 | 0 | 100 | 2099 |
| French Guiana | 8,017 | 7,969 | 7,921 | 7,873 | 3 | 5394 |

Table S16: *(continued)*

| Country – study area | fc2040 (Kha) | fc2060 (Kha) | fc2080 (Kha) | fc2100 (Kha) | loss21 (%) | yrdis |
|---|---|---|---|---|---|---|
| Grenada | 14 | 9 | 3 | 0 | 100 | 2092 |
| Guadeloupe | 66 | 59 | 51 | 44 | 45 | 2218 |
| Guatemala | 1,422 | 570 | 0 | 0 | 100 | 2073 |
| Guyana | 18,119 | 17,872 | 17,625 | 17,378 | 7 | 3508 |
| Haiti | 42 | 0 | 0 | 0 | 100 | 2050 |
| Honduras | 1,720 | 872 | 24 | 0 | 100 | 2080 |
| Jamaica | 344 | 293 | 241 | 190 | 60 | 2173 |
| Martinique | 58 | 50 | 42 | 34 | 53 | 2185 |
| Mexico | 4,012 | 1,759 | 0 | 0 | 100 | 2075 |
| Montserrat | 3 | 3 | 3 | 3 | 20 | 2490 |
| Nicaragua | 1,699 | 0 | 0 | 0 | 100 | 2059 |
| Panama | 3,628 | 3,244 | 2,860 | 2,476 | 44 | 2228 |
| Paraguay | 391 | 0 | 0 | 0 | 100 | 2051 |
| Peru | 68,521 | 66,267 | 64,013 | 61,759 | 16 | 2648 |
| Puerto Rico | 247 | 174 | 100 | 26 | 94 | 2107 |
| Saint Kitts and N. | 8 | 7 | 7 | 6 | 36 | 2273 |
| Saint Lucia | 39 | 34 | 28 | 23 | 51 | 2188 |
| Saint Martin | 0 | 0 | 0 | 0 | 100 | 2033 |
| Saint Vincent | 25 | 23 | 21 | 19 | 34 | 2278 |
| Sint Maarten | 0 | 0 | 0 | 0 | 100 | 2031 |
| Suriname | 13,418 | 13,211 | 13,005 | 12,799 | 7 | 3340 |
| Trinidad and Tobago | 259 | 212 | 165 | 118 | 66 | 2150 |
| Venezuela | 38,464 | 35,499 | 32,533 | 29,568 | 34 | 2299 |
| Virgin Isl. UK | 0 | 0 | 0 | 0 | 100 | 2039 |
| Virgin Isl. US | 4 | 1 | 0 | 0 | 100 | 2068 |
| **Africa** | | | | | | |
| Angola | 3,935 | 2,529 | 1,124 | 0 | 100 | 2095 |
| Benin | 3 | 0 | 0 | 0 | 100 | 2041 |
| Burundi | 38 | 21 | 4 | 0 | 100 | 2084 |
| Cameroon | 21,427 | 20,014 | 18,601 | 17,188 | 28 | 2343 |
| CAR | 7,911 | 6,969 | 6,027 | 5,085 | 48 | 2207 |
| Comoros | 77 | 72 | 67 | 62 | 29 | 2333 |
| Congo | 22,394 | 21,361 | 20,327 | 19,293 | 20 | 2473 |
| DRC | 103,640 | 88,997 | 74,354 | 59,711 | 55 | 2181 |
| Eq. Guinea | 2,553 | 2,495 | 2,436 | 2,377 | 10 | 2908 |
| Ethiopia | 993 | 0 | 0 | 0 | 100 | 2056 |
| Gabon | 23,753 | 23,522 | 23,290 | 23,058 | 4 | 4088 |
| Gambia | 20 | 8 | 0 | 0 | 100 | 2074 |
| Ghana | 1,149 | 0 | 0 | 0 | 100 | 2050 |
| Guinea | 84 | 0 | 0 | 0 | 100 | 2042 |
| Guinea Bissau | 171 | 69 | 0 | 0 | 100 | 2073 |
| Ivory Coast | 0 | 0 | 0 | 0 | 100 | 2036 |
| Kenya | 521 | 275 | 28 | 0 | 100 | 2082 |
| Liberia | 6,419 | 4,930 | 3,440 | 1,951 | 78 | 2126 |
| Madagascar | 2,641 | 708 | 0 | 0 | 100 | 2067 |
| Malawi | 0 | 0 | 0 | 0 | 100 | 2032 |
| Mauritius | 37 | 30 | 23 | 16 | 68 | 2145 |
| Mayotte | 2 | 0 | 0 | 0 | 100 | 2043 |
| Nigeria | 4,165 | 2,132 | 99 | 0 | 100 | 2080 |

Table S16: *(continued)*

| Country – study area | fc2040 (Kha) | fc2060 (Kha) | fc2080 (Kha) | fc2100 (Kha) | loss21 (%) | yrdis |
|---|---|---|---|---|---|---|
| Reunion | 120 | 106 | 91 | 77 | 46 | 2206 |
| Rwanda | 91 | 23 | 0 | 0 | 100 | 2066 |
| Senegal | 74 | 40 | 5 | 0 | 100 | 2082 |
| Sierra Leone | 0 | 0 | 0 | 0 | 100 | 2036 |
| South Sudan | 97 | 27 | 0 | 0 | 100 | 2067 |
| Tanzania | 870 | 657 | 443 | 229 | 84 | 2121 |
| Togo | 0 | 0 | 0 | 0 | 100 | 2037 |
| Uganda | 100 | 0 | 0 | 0 | 100 | 2043 |
| Zambia | 15 | 0 | 0 | 0 | 100 | 2044 |
| **Asia** | | | | | | |
| Australia – Queensland | 1,542 | 1,319 | 1,097 | 874 | 57 | 2178 |
| Bangladesh | 681 | 591 | 502 | 412 | 57 | 2191 |
| Bhutan | 1,661 | 1,521 | 1,380 | 1,240 | 38 | 2276 |
| Brunei | 481 | 468 | 454 | 441 | 14 | 2759 |
| Cambodia | 533 | 0 | 0 | 0 | 100 | 2044 |
| Fiji | 858 | 792 | 726 | 660 | 34 | 2300 |
| India – Andaman and N. | 534 | 497 | 459 | 422 | 31 | 2324 |
| India – North-East | 4,797 | 4,034 | 3,272 | 2,509 | 64 | 2165 |
| India – West. Ghats | 1,302 | 367 | 0 | 0 | 100 | 2067 |
| Indonesia | 98,275 | 79,478 | 60,680 | 41,883 | 70 | 2144 |
| Laos | 5,543 | 2,779 | 14 | 0 | 100 | 2080 |
| Malaysia | 15,812 | 11,477 | 7,141 | 2,806 | 89 | 2112 |
| Myanmar | 10,424 | 7,120 | 3,816 | 512 | 97 | 2103 |
| New Caledonia | 806 | 758 | 709 | 661 | 27 | 2372 |
| Papua New Guinea | 38,330 | 37,356 | 36,383 | 35,409 | 12 | 2827 |
| Philippines | 10,892 | 9,031 | 7,170 | 5,309 | 64 | 2157 |
| Singapore | 11 | 8 | 5 | 2 | 88 | 2113 |
| Solomon Isl. | 2,704 | 2,669 | 2,634 | 2,599 | 6 | 3584 |
| Sri Lanka | 1,313 | 1,032 | 751 | 470 | 78 | 2133 |
| Thailand | 4,763 | 3,711 | 2,659 | 1,607 | 78 | 2130 |
| Timor-Leste | 54 | 31 | 7 | 0 | 100 | 2086 |
| Vanuatu | 1,141 | 1,130 | 1,118 | 1,107 | 4 | 4062 |
| Vietnam | 5,541 | 3,483 | 1,424 | 0 | 100 | 2093 |

Table S17: **Forest cover projections per region and continent**. Projected areas of forest cover are given in thousand hectares (Kha) for four dates in the future (2040, 2060, 2080, and 2100). Projections were made using the forest cover in 2020 and the mean annual deforested area in the ten-year period 2010–2020 ("fc2000" and $d$ respectively in Table S15), assuming a "business-as-usual" scenario of deforestation. Column "loss21" indicates the projected percentage of forest cover loss during the 21[st] century (2100 vs. 2000). At the continental level, it makes less sense to compute the year at which all the forest will have disappeared, as some countries might conserve forest for a very long time, even though they account for a very small proportion of the total forest area at the continental scale. Instead, we computed the estimated year at which 75% of the forest cover in 2000 will have disappeared ("yr75dis").

| Region | fc2040 (Kha) | fc2060 (Kha) | fc2080 (Kha) | fc2100 (Kha) | loss21 (%) | yr75dis |
|---|---|---|---|---|---|---|
| India | 6,633 | 4,898 | 3,731 | 2,931 | 73 | 2085 |
| Brazil | 307,545 | 280,142 | 252,739 | 225,336 | 40 | 2204 |
| America | 570,321 | 519,772 | 472,135 | 427,790 | 38 | 2220 |
| Africa | 203,302 | 174,982 | 150,357 | 129,045 | 53 | 2163 |
| Asia | 207,999 | 169,650 | 132,403 | 98,922 | 67 | 2117 |
| All continents | 981,622 | 864,404 | 754,895 | 655,757 | 48 | 2192 |

#### Table S18 – Cumulative carbon emissions associated with deforestation

Table S18: **Cumulative carbon emissions associated with future deforestation**. We computed the cumulative carbon emissions associated with future deforestation from 2020 for each study area (C in Gg, $10^9$ g). To do so, we used our maps of projected forest cover change together with available global or pantropical maps of aboveground biomass (either the WUR, WHRC or CCI map). We present here the results obtained with the ESA CCI aboveground biomass map by Santoro et al. (2021).

| Country – study area | C2040 (Gg) | C2060 (Gg) | C2080 (Gg) | C2100 (Gg) |
|---|---|---|---|---|
| **America** | | | | |
| Antigua and B. | 11 | 30 | 48 | 48 |
| Bahamas | 493 | 930 | 1,327 | 1,327 |
| Barbados | 28 | 63 | 92 | 92 |
| Belize | 13,917 | 28,352 | 43,462 | 59,897 |
| Bolivia | 348,852 | 727,365 | 1,135,780 | 1,571,465 |
| Brazil – Acre | 130,941 | 287,166 | 481,319 | 700,694 |
| Brazil – Alagoas | 585 | 3,498 | 3,498 | 3,498 |
| Brazil – Amapa | 20,717 | 50,081 | 98,636 | 163,592 |
| Brazil – Amazonas | 421,635 | 828,613 | 1,250,705 | 1,687,342 |
| Brazil – Bahia | 16,232 | 40,364 | 76,668 | 124,938 |
| Brazil – Ceara | 405 | 996 | 996 | 996 |
| Brazil – Espirito Santo | 3,761 | 11,565 | 28,282 | 28,282 |
| Brazil – Goias | 13,396 | 19,393 | 19,393 | 19,393 |
| Brazil – Maranhao | 95,874 | 219,895 | 302,967 | 302,967 |
| Brazil – Mato Grosso | 580,235 | 1,219,046 | 1,918,403 | 2,713,595 |
| Brazil – Mato Grosso do Sul | 8,599 | 20,026 | 35,051 | 35,051 |
| Brazil – Minas Gerais | 58,150 | 92,244 | 92,244 | 92,244 |
| Brazil – Para | 1,071,058 | 2,187,322 | 3,387,051 | 4,636,768 |
| Brazil – Paraiba | 197 | 1,272 | 1,272 | 1,272 |
| Brazil – Parana | 27,483 | 58,610 | 100,392 | 151,484 |
| Brazil – Pernambouco | 604 | 3,334 | 3,970 | 3,970 |
| Brazil – Piaui | 1,162 | 1,631 | 1,631 | 1,631 |
| Brazil – Rio de Janeiro | 6,987 | 19,748 | 47,673 | 73,653 |
| Brazil – Rio Grande do Norte | 138 | 494 | 494 | 494 |
| Brazil – Rio Grande do Sul | 13,313 | 29,906 | 54,549 | 86,708 |
| Brazil – Rondonia | 366,527 | 749,292 | 1,182,271 | 1,648,580 |
| Brazil – Roraima | 157,889 | 323,817 | 498,378 | 682,608 |
| Brazil – Santa Catarina | 17,094 | 37,932 | 67,251 | 103,633 |
| Brazil – Sao Paulo | 20,225 | 47,756 | 89,045 | 142,571 |
| Brazil – Sergipe | 377 | 1,703 | 1,703 | 1,703 |
| Brazil – Tocantins | 49,936 | 63,510 | 63,510 | 63,510 |
| Colombia | 434,390 | 888,778 | 1,357,504 | 1,888,621 |
| Costa Rica | 12,782 | 31,103 | 52,548 | 76,496 |
| Cuba | 6,249 | 14,199 | 23,038 | 32,821 |
| Dominica | 142 | 313 | 492 | 675 |
| Dominican Rep. | 13,142 | 28,693 | 46,157 | 56,949 |
| Ecuador | 60,190 | 141,016 | 235,274 | 339,528 |
| El Salvador | 1,017 | 1,984 | 2,885 | 3,631 |
| French Guiana | 6,529 | 12,515 | 18,709 | 24,714 |
| Grenada | 193 | 377 | 572 | 706 |
| Guadeloupe | 199 | 375 | 576 | 818 |

Table S18: *(continued)*

| Country – study area | C2040 (Gg) | C2060 (Gg) | C2080 (Gg) | C2100 (Gg) |
|---|---|---|---|---|
| Guatemala | 49,603 | 100,908 | 135,598 | 135,598 |
| Guyana | 30,975 | 59,519 | 89,605 | 120,609 |
| Haiti | 2,635 | 4,024 | 4,024 | 4,024 |
| Honduras | 62,847 | 122,192 | 186,338 | 188,247 |
| Jamaica | 2,255 | 4,490 | 6,631 | 8,716 |
| Martinique | 159 | 327 | 512 | 716 |
| Mexico | 96,494 | 196,664 | 283,494 | 283,494 |
| Montserrat | 3 | 7 | 11 | 15 |
| Nicaragua | 102,340 | 231,755 | 231,755 | 231,755 |
| Panama | 28,372 | 56,353 | 84,951 | 115,047 |
| Paraguay | 37,533 | 58,591 | 58,591 | 58,591 |
| Peru | 314,567 | 627,961 | 946,086 | 1,272,433 |
| Puerto Rico | 3,810 | 8,119 | 12,835 | 17,905 |
| Saint Kitts and N. | 14 | 33 | 55 | 78 |
| Saint Lucia | 124 | 260 | 410 | 568 |
| Saint Martin | 6 | 6 | 6 | 6 |
| Saint Vincent | 41 | 79 | 120 | 162 |
| Sint Maarten | 3 | 3 | 3 | 3 |
| Suriname | 29,981 | 57,109 | 84,601 | 113,033 |
| Trinidad and Tobago | 2,696 | 6,165 | 10,200 | 14,505 |
| Venezuela | 229,052 | 513,123 | 837,485 | 1,180,325 |
| Virgin Isl. UK | 72 | 72 | 72 | 72 |
| Virgin Isl. US | 109 | 225 | 269 | 269 |
| **Africa** | | | | |
| Angola | 112,054 | 235,388 | 365,560 | 493,217 |
| Benin | 424 | 517 | 517 | 517 |
| Burundi | 933 | 2,216 | 3,316 | 3,878 |
| Cameroon | 132,303 | 290,031 | 458,714 | 637,848 |
| CAR | 99,960 | 200,920 | 306,863 | 420,044 |
| Comoros | 168 | 338 | 514 | 693 |
| Congo | 105,366 | 222,716 | 351,680 | 488,487 |
| DRC | 1,539,813 | 3,305,004 | 5,296,020 | 7,444,442 |
| Eq. Guinea | 6,766 | 13,989 | 21,596 | 29,644 |
| Ethiopia | 111,380 | 214,296 | 214,296 | 214,296 |
| Gabon | 25,833 | 53,060 | 81,886 | 111,061 |
| Gambia | 231 | 435 | 591 | 591 |
| Ghana | 130,416 | 252,021 | 252,021 | 252,021 |
| Guinea | 47,702 | 56,084 | 56,084 | 56,084 |
| Guinea Bissau | 3,774 | 7,390 | 10,214 | 10,214 |
| Ivory Coast | 319,052 | 319,052 | 319,052 | 319,052 |
| Kenya | 11,128 | 23,251 | 36,556 | 38,106 |
| Liberia | 99,738 | 241,138 | 425,959 | 641,118 |
| Madagascar | 128,144 | 272,655 | 330,993 | 330,993 |
| Malawi | 2,253 | 2,253 | 2,253 | 2,253 |
| Mauritius | 130 | 294 | 500 | 738 |
| Mayotte | 558 | 656 | 656 | 656 |
| Nigeria | 118,606 | 265,210 | 417,524 | 427,625 |
| Reunion | 386 | 806 | 1,256 | 1,740 |
| Rwanda | 3,599 | 11,537 | 11,537 | 11,537 |

Table S18: *(continued)*

| Country – study area | C2040 (Gg) | C2060 (Gg) | C2080 (Gg) | C2100 (Gg) |
|---|---|---|---|---|
| Senegal | 511 | 1,034 | 1,488 | 1,499 |
| Sierra Leone | 79,457 | 79,457 | 79,457 | 79,457 |
| South Sudan | 3,999 | 6,595 | 6,595 | 6,595 |
| Tanzania | 10,535 | 22,736 | 36,436 | 51,049 |
| Togo | 4,584 | 4,584 | 4,584 | 4,584 |
| Uganda | 38,594 | 45,772 | 45,772 | 45,772 |
| Zambia | 5,202 | 6,203 | 6,203 | 6,203 |
| **Asia** | | | | |
| Australia – Queensland | 12,555 | 23,142 | 34,972 | 49,748 |
| Bangladesh | 2,585 | 4,666 | 6,622 | 8,494 |
| Bhutan | 6,777 | 13,306 | 19,858 | 26,433 |
| Brunei | 787 | 1,684 | 2,584 | 3,527 |
| Cambodia | 136,413 | 170,714 | 170,714 | 170,714 |
| Fiji | 1,593 | 3,647 | 5,822 | 8,235 |
| India – Andaman and N. | 1,051 | 2,284 | 3,610 | 4,982 |
| India – North-East | 20,049 | 45,075 | 71,686 | 98,948 |
| India – West. Ghats | 15,581 | 38,396 | 48,217 | 48,217 |
| Indonesia | 1,215,891 | 2,796,927 | 4,738,167 | 6,910,733 |
| Laos | 127,455 | 252,199 | 389,729 | 389,729 |
| Malaysia | 283,988 | 660,935 | 1,077,382 | 1,516,728 |
| Myanmar | 116,201 | 236,921 | 361,399 | 491,056 |
| New Caledonia | 1,308 | 2,780 | 4,348 | 6,005 |
| Papua New Guinea | 96,558 | 185,984 | 280,257 | 380,681 |
| Philippines | 93,000 | 196,682 | 301,730 | 414,931 |
| Singapore | 95 | 214 | 346 | 485 |
| Solomon Isl. | 1,687 | 4,017 | 6,523 | 9,071 |
| Sri Lanka | 4,795 | 10,940 | 17,564 | 24,842 |
| Thailand | 34,602 | 75,562 | 119,061 | 164,791 |
| Timor-Leste | 1,974 | 3,806 | 5,612 | 6,123 |
| Vanuatu | 425 | 956 | 1,475 | 1,962 |
| Vietnam | 81,330 | 159,950 | 247,926 | 328,684 |

#### Table S19 – Confidence interval of the annual deforested area

Table S19: **Confidence interval of the annual deforested area**. We computed the 95% confidence interval of the annual deforested area $d$ (in ha/yr) for each study area. We used the deforestation observations $d_t$ for the ten years $t$ from 2010 to 2020. The lower and upper bounds of the confidence interval were denoted $d'$ and $d''$, respectively. Forest cover areas (in Kha) for the years 2010 and 2020 ("fc2010" and "fc2020") are also provided for comparison.

| Country – study area | fc2010 (Kha) | fc2020 (Kha) | $d$ (ha/yr) | $d'$ (ha/yr) | $d''$ (ha/yr) |
|---|---|---|---|---|---|
| **America** | | | | | |
| Antigua and B. | 4 | 3 | 52 | 22 | 83 |
| Bahamas | 115 | 98 | 1,657 | 813 | 2,502 |
| Barbados | 4 | 3 | 64 | 18 | 110 |
| Belize | 1,328 | 1,198 | 12,996 | 7,657 | 18,334 |
| Bolivia | 30,485 | 28,739 | 174,616 | 96,497 | 252,735 |
| Brazil – Acre | 13,307 | 12,860 | 44,684 | 38,303 | 51,065 |
| Brazil – Alagoas | 98 | 89 | 901 | 477 | 1,325 |
| Brazil – Amapa | 11,565 | 11,458 | 10,677 | 7,654 | 13,699 |
| Brazil – Amazonas | 146,852 | 145,346 | 150,663 | 104,470 | 196,856 |
| Brazil – Bahia | 2,097 | 1,936 | 16,103 | 9,030 | 23,176 |
| Brazil – Ceara | 48 | 40 | 849 | 454 | 1,243 |
| Brazil – Espirito Santo | 417 | 387 | 2,945 | 1,202 | 4,689 |
| Brazil – Goias | 481 | 358 | 12,241 | 8,147 | 16,335 |
| Brazil – Maranhao | 3,930 | 3,259 | 67,024 | 37,602 | 96,446 |
| Brazil – Mato Grosso | 33,283 | 30,333 | 294,945 | 236,938 | 352,951 |
| Brazil – Mato Grosso do Sul | 736 | 654 | 8,193 | 6,016 | 10,370 |
| Brazil – Minas Gerais | 1,277 | 954 | 32,323 | 16,580 | 48,067 |
| Brazil – Para | 91,982 | 87,764 | 421,863 | 346,357 | 497,370 |
| Brazil – Paraiba | 41 | 38 | 308 | 122 | 494 |
| Brazil – Parana | 2,673 | 2,475 | 19,780 | 12,721 | 26,839 |
| Brazil – Pernambouco | 119 | 109 | 936 | 432 | 1,439 |
| Brazil – Piaui | 74 | 55 | 1,967 | 846 | 3,087 |
| Brazil – Rio de Janeiro | 736 | 694 | 4,143 | 1,141 | 7,146 |
| Brazil – Rio Grande do Norte | 25 | 22 | 312 | 195 | 428 |
| Brazil – Rio Grande do Sul | 2,214 | 2,074 | 14,004 | 9,402 | 18,607 |
| Brazil – Rondonia | 13,800 | 12,436 | 136,426 | 114,420 | 158,431 |
| Brazil – Roraima | 16,228 | 15,593 | 63,551 | 20,622 | 106,479 |
| Brazil – Santa Catarina | 2,485 | 2,355 | 13,020 | 9,833 | 16,208 |
| Brazil – Sao Paulo | 2,781 | 2,653 | 12,862 | 6,930 | 18,794 |
| Brazil – Sergipe | 61 | 54 | 656 | 226 | 1,086 |
| Brazil – Tocantins | 1,341 | 954 | 38,777 | 22,144 | 55,410 |
| Colombia | 66,802 | 64,254 | 254,796 | 212,523 | 297,069 |
| Costa Rica | 2,277 | 2,132 | 14,520 | 9,291 | 19,748 |
| Cuba | 1,297 | 1,155 | 14,086 | 7,725 | 20,446 |
| Dominica | 70 | 67 | 254 | 126 | 383 |
| Dominican Rep. | 993 | 871 | 12,164 | 6,964 | 17,364 |
| Ecuador | 14,903 | 14,391 | 51,193 | 31,232 | 71,154 |
| El Salvador | 109 | 97 | 1,220 | 796 | 1,645 |
| French Guiana | 8,088 | 8,064 | 2,390 | 1,922 | 2,857 |
| Grenada | 22 | 19 | 264 | 57 | 470 |
| Guadeloupe | 77 | 73 | 370 | 237 | 502 |

Table S19: *(continued)*

| Country – study area | fc2010 (Kha) | fc2020 (Kha) | $d$ (ha/yr) | $d'$ (ha/yr) | $d''$ (ha/yr) |
|---|---|---|---|---|---|
| Guatemala | 2,702 | 2,275 | 42,648 | 33,295 | 52,000 |
| Guyana | 18,489 | 18,366 | 12,342 | 8,474 | 16,210 |
| Haiti | 162 | 122 | 4,001 | 2,655 | 5,346 |
| Honduras | 2,993 | 2,569 | 42,409 | 31,042 | 53,775 |
| Jamaica | 421 | 395 | 2,572 | 1,403 | 3,741 |
| Martinique | 70 | 66 | 400 | 229 | 571 |
| Mexico | 7,390 | 6,264 | 112,620 | 85,558 | 139,682 |
| Montserrat | 3 | 3 | 7 | 3 | 11 |
| Nicaragua | 4,262 | 3,408 | 85,457 | 47,013 | 123,901 |
| Panama | 4,204 | 4,012 | 19,202 | 13,907 | 24,496 |
| Paraguay | 1,440 | 1,090 | 34,950 | 19,137 | 50,764 |
| Peru | 71,901 | 70,775 | 112,691 | 95,602 | 129,780 |
| Puerto Rico | 358 | 321 | 3,688 | 1,566 | 5,809 |
| Saint Kitts and N. | 9 | 9 | 34 | 12 | 56 |
| Saint Lucia | 47 | 44 | 261 | 124 | 398 |
| Saint Martin | 1 | 0 | 24 | 4 | 44 |
| Saint Vincent | 28 | 27 | 106 | 54 | 159 |
| Sint Maarten | 0 | 0 | 10 | 2 | 18 |
| Suriname | 13,727 | 13,624 | 10,316 | 8,044 | 12,589 |
| Trinidad and Tobago | 330 | 307 | 2,355 | 382 | 4,328 |
| Venezuela | 42,913 | 41,430 | 148,279 | 99,388 | 197,169 |
| Virgin Isl. UK | 3 | 2 | 96 | 28 | 164 |
| Virgin Isl. US | 8 | 7 | 137 | 32 | 242 |
| **Africa** | | | | | |
| Angola | 6,044 | 5,341 | 70,293 | 43,994 | 96,591 |
| Benin | 47 | 32 | 1,463 | 836 | 2,090 |
| Burundi | 64 | 55 | 856 | 251 | 1,461 |
| Cameroon | 23,546 | 22,840 | 70,641 | 57,279 | 84,002 |
| CAR | 9,325 | 8,854 | 47,114 | 33,224 | 61,005 |
| Comoros | 87 | 83 | 263 | 136 | 390 |
| Congo | 23,945 | 23,428 | 51,689 | 37,980 | 65,399 |
| DRC | 125,605 | 118,283 | 732,153 | 562,240 | 902,066 |
| Eq. Guinea | 2,642 | 2,612 | 2,941 | 2,173 | 3,710 |
| Ethiopia | 2,824 | 2,214 | 61,036 | 32,558 | 89,514 |
| Gabon | 24,101 | 23,985 | 11,595 | 7,866 | 15,324 |
| Gambia | 42 | 31 | 566 | 98 | 1,035 |
| Ghana | 4,443 | 3,345 | 109,799 | 62,116 | 157,481 |
| Guinea | 1,213 | 837 | 37,643 | 28,448 | 46,838 |
| Guinea Bissau | 323 | 272 | 5,078 | 3,768 | 6,387 |
| Ivory Coast | 6,299 | 3,954 | 234,522 | 153,578 | 315,466 |
| Kenya | 891 | 768 | 12,324 | 5,084 | 19,564 |
| Liberia | 8,653 | 7,908 | 74,471 | 55,838 | 93,105 |
| Madagascar | 5,541 | 4,574 | 96,645 | 66,586 | 126,704 |
| Malawi | 70 | 39 | 3,147 | 1,891 | 4,404 |
| Mauritius | 47 | 43 | 347 | 18 | 675 |
| Mayotte | 17 | 12 | 511 | 148 | 874 |
| Nigeria | 7,214 | 6,197 | 101,634 | 66,695 | 136,573 |
| Reunion | 142 | 134 | 722 | 480 | 964 |
| Rwanda | 195 | 160 | 3,434 | 1,296 | 5,572 |

Table S19: *(continued)*

| Country – study area | fc2010 (Kha) | fc2020 (Kha) | $d$ (ha/yr) | $d'$ (ha/yr) | $d''$ (ha/yr) |
|---|---|---|---|---|---|
| Senegal | 126 | 109 | 1,736 | 41 | 3,431 |
| Sierra Leone | 2,260 | 1,414 | 84,563 | 59,064 | 110,062 |
| South Sudan | 201 | 167 | 3,493 | 2,248 | 4,738 |
| Tanzania | 1,191 | 1,084 | 10,689 | 6,573 | 14,804 |
| Togo | 103 | 66 | 3,693 | 1,930 | 5,456 |
| Uganda | 1,087 | 758 | 32,878 | 19,529 | 46,226 |
| Zambia | 114 | 81 | 3,294 | 1,931 | 4,658 |
| **Asia** | | | | | |
| Australia – Queensland | 1,876 | 1,764 | 11,131 | 6,349 | 15,912 |
| Bangladesh | 816 | 771 | 4,489 | 2,907 | 6,071 |
| Bhutan | 1,872 | 1,802 | 7,022 | 4,133 | 9,912 |
| Brunei | 501 | 494 | 668 | 499 | 838 |
| Cambodia | 3,864 | 2,753 | 111,024 | 79,030 | 143,017 |
| Fiji | 958 | 924 | 3,300 | 1,767 | 4,832 |
| India – Andaman and N. | 591 | 572 | 1,875 | 388 | 3,363 |
| India – North-East | 5,941 | 5,560 | 38,134 | 24,962 | 51,305 |
| India – West. Ghats | 2,704 | 2,236 | 46,740 | 28,945 | 64,536 |
| Indonesia | 126,473 | 117,072 | 939,867 | 646,351 | 1,233,384 |
| Laos | 9,690 | 8,308 | 138,221 | 108,579 | 167,863 |
| Malaysia | 22,315 | 20,147 | 216,762 | 161,714 | 271,809 |
| Myanmar | 15,380 | 13,728 | 165,195 | 108,006 | 222,384 |
| New Caledonia | 879 | 855 | 2,425 | 1,490 | 3,359 |
| Papua New Guinea | 39,791 | 39,304 | 48,691 | 29,465 | 67,917 |
| Philippines | 13,684 | 12,753 | 93,052 | 49,915 | 136,189 |
| Singapore | 15 | 14 | 147 | 90 | 204 |
| Solomon Isl. | 2,757 | 2,739 | 1,751 | 1,065 | 2,437 |
| Sri Lanka | 1,735 | 1,594 | 14,060 | 8,139 | 19,981 |
| Thailand | 6,341 | 5,815 | 52,603 | 33,106 | 72,100 |
| Timor-Leste | 89 | 78 | 1,173 | 422 | 1,925 |
| Vanuatu | 1,158 | 1,152 | 564 | 256 | 873 |
| Vietnam | 8,628 | 7,599 | 102,909 | 73,716 | 132,102 |

### 4 Legends for Supplementary Data

Supplementary Data are available at https://forestatrisk.cirad.fr/data-s.html.

##### Data S1 – Uncertainty around projected forest cover

Data S1: **Uncertainty around projected forest cover**. Past and projected forest cover change by study area are given in thousand hectares (Kha). The mean annual deforested area $d$ for the ten-year period 2010–2020 is given in hectare per year (ha/yr). The corresponding mean annual deforestation rate $p$ is also provided in percent per year (%/yr), with one decimal precision, to be able to compare the intensity of deforestation between study areas. Projections were made using the forest cover in 2020 ("fc2000") and either (i) the mean annual deforested area $d$ for the ten-year period 2010–2020, (ii) the lower bound $d'$ of the confidence interval for the annual deforested area, or (iii) the upper bound $d''$ of the confidence interval for the annual deforested area. We considered a business-as-usual scenario of deforestation (deforestation constant through time) for the projections. Column "loss21" indicates the projected percentage of forest cover loss during the $21^{st}$ century (2100 vs. 2000). Column "yrdis" indicates the estimated year at which all the forest of the study area will have disappeared.

##### Data S2 – Uncertainty around projected carbon emissions

Data S2: **Uncertainty around projected carbon emissions**. We combined our maps of projected forest cover change together with the aboveground biomass map by Avitabile et al. (2016) to compute the cumulative carbon emissions associated with future deforestation from 2020 for each study area (C in Gg=$10^9$ g). Maps of projected forest cover change were derived using either (i) the mean annual deforested area $d$ for the ten-year period 2010–2020, (ii) the lower bound $d'$ of the confidence interval for the annual deforested area, or (iii) the upper bound $d''$ of the confidence interval for the annual deforested area. Column "C2020" indicates the carbon emissions associated with past deforestation during the period 2010–2020 for comparison.

IPCC. 2014. Climate Change 2014: Synthesis Report. Contribution of Working Groups I, II and III to the Fifth Assessment Report of the Intergovernmental Panel on Climate Change. Page 151 (R. K. Pachauri and L. A. Meyer, Eds.). The Intergovernmental Panel on Climate Change, IPCC, Geneva, Switzerland.

Liu, C., M. White, and G. Newell. 2011. Measuring and comparing the accuracy of species distribution models with presence-absence data. Ecography 34:232–243.

Ludeke, A. K., R. C. Maggio, and L. M. Reid. 1990. An analysis of anthropogenic deforestation using logistic regression and GIS. Journal of Environmental Management 31:247–259.

Mas, J.-F., M. Paegelow, B. de Jong, G. Guerrero, M. Follador, M. Olgien, J. R. Díaz, M. A. Castillo, and T. García. 2007. Modelling tropical deforestation: A comparison of approaches. Symposium Remote Sensing of Environment:3.

McGroddy, M. E., T. Daufresne, and L. O. Hedin. 2004. Scaling of C:N:P stoichiometry in forests worldwide: implications of terrestrial redfield-type ratios. Ecology 85:2390–2401.

Pedregosa, F., G. Varoquaux, A. Gramfort, V. Michel, B. Thirion, O. Grisel, M. Blondel, P. Prettenhofer, R. Weiss, V. Dubourg, J. Vanderplas, A. Passos, D. Cournapeau, M. Brucher, M. Perrot, and E. Duchesnay. 2011. Scikit-Learn: Machine learning in Python. Journal of Machine Learning Research 12:2825–2830.

Ploton, P., F. Mortier, M. Réjou-Méchain, N. Barbier, N. Picard, V. Rossi, C. Dormann, G. Cornu, G. Viennois, N. Bayol, A. Lyapustin, S. Gourlet-Fleury, and R. Pélissier. 2020. Spatial validation reveals poor predictive performance of large-scale ecological mapping models. Nature Communications 11.

Pontius, R. G. J., and M. Millones. 2011. Death to Kappa: Birth of quantity disagreement and allocation disagreement for accuracy assessment. International Journal of Remote Sensing 32:4407–4429.

Pontius, R., W. Boersma, J.-C. Castella, K. Clarke, T. de Nijs, C. Dietzel, Z. Duan, E. Fotsing, N. Goldstein, K. Kok, E. Koomen, C. Lippitt, W. McConnell, A. Mohd Sood, B. Pijanowski, S. Pithadia, S. Sweeney, T. Trung, A. Veldkamp, and P. Verburg. 2008. Comparing the input, output, and validation maps for several models of land change. The Annals of Regional Science 42:11–37.

Python Software Foundation. 2020. The Python language reference. Python Software Foundation.

R Core Team. 2020. R: A language and environment for statistical computing. R Foundation for Statistical Computing, Vienna, Austria.

Rosa, I. M. D., D. Purves, J. M. B. Carreiras, and R. M. Ewers. 2014. Modelling land cover change in the brazilian amazon: Temporal changes in drivers and calibration issues. Regional Environmental Change 15:123–137.

Saatchi, S. S., N. L. Harris, S. Brown, M. Lefsky, E. T. A. Mitchard, W. Salas, B. R. Zutta, W. Buermann, S. L. Lewis, S. Hagen, S. Petrova, L. White, M. Silman, and A. Morel. 2011. Benchmark map of forest carbon stocks in tropical regions across three continents. Proceedings of the National Academy of Sciences 108:9899–9904.

Santoro, M., O. Cartus, N. Carvalhais, D. M. A. Rozendaal, V. Avitabile, A. Araza, S. de Bruin, M. Herold, S. Quegan, P. Rodríguez-Veiga, H. Balzter, J. Carreiras, D. Schepaschenko, M. Korets, M. Shimada, T. Itoh, Álvaro Moreno Martínez, J. Cavlovic, R. C. Gatti, P. da Conceição Bispo, N. Dewnath, N. Labrière, J. Liang, J. Lindsell, E. T. A. Mitchard, A. Morel, A. M. P. Pascagaza, C. M. Ryan, F. Slik, G. V. Laurin, H. Verbeeck, A. Wijaya, and S. Willcock. 2021. The global forest above-ground biomass pool for 2010 estimated from high-resolution satellite observations. Earth System Science Data 13:3927–3950.

Schwartz, N. B., T. M. Aide, J. Graesser, H. R. Grau, and M. Uriarte. 2020. Reversals of reforestation across Latin America limit climate mitigation potential of tropical forests. Frontiers in Forests and Global Change 3.

Soares-Filho, B. S., G. C. Cerqueira, and C. L. Pennachin. 2002. DINAMICA - a stochastic cellular automata model designed to simulate the landscape dynamics in an Amazonian colonization frontier. Ecological Modelling 154:217–235.

Thuiller, W., B. Lafourcade, R. Engler, and M. B. Araujo. 2009. BIOMOD - a platform for ensemble forecasting of species distributions. Ecography 32:369–373.

UNEP-WCMC, and IUCN. 2020. Protected Planet: The World Database on Protected Areas (WDPA). On-line; 01-03/2020, Cambridge, UK: UNEP-WCMC and IUCN. Available at: www.protectedplanet.net. https://www.protectedplanet.net.

Valle, D., J. Hyde, M. Marsik, and S. Perz. 2020. Improved inference and prediction for imbalanced binary big data using case-control sampling: A case study on deforestation in the Amazon region. Remote Sensing 12:1268.

Vancutsem, C., F. Achard, J.-F. Pekel, G. Vieilledent, S. Carboni, D. Simonetti, J. Gallego, L. E. O. C. Aragão, and R. Nasi. 2021. Long-term (1990–2019) monitoring of forest cover changes in the humid tropics. Science Advances 7.

Vieilledent, G. 2021. `forestatrisk`: A Python package for modelling and forecasting deforestation in the tropics. Journal of Open Source Software 6:2975.

Vieilledent, G., C. Grinand, and R. Vaudry. 2013. Forecasting deforestation and carbon emissions in tropical developing countries facing demographic expansion: A case study in Madagascar. Ecology and Evolution 3:1702–1716.

Vieilledent, G., M. Nourtier, C. Grinand, M. Pedrono, A. Clausen, T. Rabetrano, J.-R. Rakotoarijaona, B. Rakotoarivelo, F. A. Rakotomalala, L. Rakotomalala, A. Razafimpahanana, J. M. Ralison, and F. Achard. 2020. It's not just poverty: Unregulated global market and bad governance explain unceasing deforestation in Western Madagascar. bioRxiv 2020.07.30.229104.

Zanella, L., A. M. Folkard, G. A. Blackburn, and L. M. T. Carvalho. 2017. How well does random forest analysis model deforestation and forest fragmentation in the Brazilian Atlantic forest? Environmental and Ecological Statistics 24:529–549.

Zarin, D. J., N. L. Harris, A. Baccini, D. Aksenov, M. C. Hansen, C. Azevedo-Ramos, T. Azevedo, B. A. Margono, A. C. Alencar, C. Gabris, A. Allegretti, P. Potapov, M. Farina, W. S. Walker, V. S. Shevade, T. V. Loboda, S. Turubanova, and A. Tyukavina. 2016. Can carbon emissions from tropical deforestation drop by 50. Global Change Biology 22:1336–1347.
